# *miR-34b* associates with epithelial-mesenchymal transition and regulates *BMP7, CAV1, ID2* and *FN1* in human cervical cancer

**DOI:** 10.1101/2021.09.02.458804

**Authors:** Nalini Venkatesan, Ashley Xavier, Sindhu K.J., Sandhya Sundhram, Himanshu Sinha, Devarajan Karunagaran

## Abstract

**Purpose:** We aimed to identify the epithelial-mesenchymal transition and associated putative targets of the *miR-34* family in transcriptomic data of cervical epithelial squamous cell carcinoma (CESC) to find new therapeutic targets for better disease management.

**Methods:** A combined computational analysis of the *miR-34* family; gene expression in heterogeneous primary CESCs derived from TCGA; and the integration of *miR-34b* and EMT-regulated genes was performed. Four EMT-associated *miR-34b* gene targets were analysed in primary human CESC and non-cancerous cervical tissues by qRT-PCR. Effects of endogenous *miR-34b* expression and its associated gene modulations in cervical cancer cells (C33A and HeLa) were analysed using qRT-PCR, western blotting and immunofluorescence, transwell migration and invasion assays.

**Results:** The results showed that the *miR-34* family might regulate the mTOR pathway, cell cycle (*CCND2*) and cell adhesion functions (*FZD4*). Further, we showed that a low negative correlation (r^2^= −0.07) between *miR-34b/EMĨ* score and four EMT signature genes (*E>MP7, CAV1, ID2, FN1*) were significantly regulated by *miR-34b*. Also, *FN1* was indicated as its putative target with the highest negative EMT score and high binding energy between MRE/3’UTR. Further, these transcriptomic signatures in CESC revealed a significant inverse correlation across the stages of primary human CESC. These genes were repressed at transcriptional and translational levels in *miR-34b-3p* expressing C33A and HeLa cells, contributing to their reduced cell migration and invasive properties.

**Conclusions:** Our studies revealed the potential targets of the *miR-34* family, especially *miR-34b*, that can emerge as potential biomarkers and promising therapeutic targets in CESC disease management.

## 1. Introduction

Carcinoma of the uterine cervix, the second largest cancer in women, is on a steady rise, leading to cancer-related deaths worldwide^1,2^. Despite the awareness of cervical cancer prevention and improvement in screening systems, there is a higher incidence of cervical cancer in developing countries, especially in India, accounting for 16% of total global cases^3^. Indian women develop cervical cancer with a cumulative risk of 1.6% and face a cumulative death risk of 1.0%, contributing to about one-third of the global cervical cancer deaths^4^. Advanced and recurrent cervical cancer results in a poor prognosis with a one-year survival rate of 10-20%^5^. The existence of invasion, metastasis and drug resistance in cervical cancer results in treatment failure and induces most of its related deaths. A major contributor to the poor prognosis of cervical cancer is Epithelial-Mesenchymal Transition (EMT) initiation, triggered by the cellular processes, which include the dysregulation of certain tumour suppressors, oncogenes, transcription factors, growth factor signalling, and miRNAs.EMT, a pivotal and intricate process, plays an essential role in tumour cell invasion and migration by involving numerous cytokines, transcription factors and cellular pathways such as *NF-κB, PI3K*, Wnt, and TGfβ, etc.^6^ During the EMT process, the tumour cells undergo a “cadherin switch”, lose their epithelial molecule (E-cadherin) and gain the mesenchymal molecule (N-cadherin), attributing to tumour invasion and metastasis^7^. So, the dysregulation of these EMT markers indicates the poor prognosis of cancer patients, especially epithelial cancer cell types like cervical cancer. A positive correlation exists between EMT phenotype and increased tumour progression, invasion, and metastasis in primary cervical cancers^8^. Despite many studies showing stimulation of*IL6*^9^ and increased *REXI*^10^ and *ATG5*^11^ expression in the EMT induction, there is a lack of concrete molecular insights involved in the EMT regulatory factors to halt cervical cancer progression. In this context, microRNAs (miRNAs), small 18-22 nucleotides long non-coding RNAs, as potential modulators of tumour suppressors and oncogenes acquire attention as they act as RNA silencers or post-transcriptional regulators of messenger RNAs (mRNAs)^12,13^. Changes in miRNA and gene expression have been documented in tumour initiation and progression^14^, such as decreased expression of miRNAs, explaining its intrinsic role in tumour suppression^15^. Multiple miRNAs target the EMT regulatory factors inhibiting or promoting EMT in cervical cancers. These multiple miRNAs include *miR-200* family, *miR-141, miR-10a, miR-19a, miR-29a, miR-361-5p, miR-429* and *miR-155*^16–19^.Earlier, several studies have indicated the role of *miR-34b* in many cancers, including cervical cancer^20–22^. In addition, differential expression of *miR-34b* was reported between high-grade cervical intraepithelial neoplasia and normal cervical epithelial samples^23^. *miR-34b*, along with *miR-34a* and *miR-34c*, belong to the *miR-34* family. *miR-34a* is encoded by exon 2 of chromosome 1 (ch1p36.22), while *miR-34b/c* exists as the polycistronic transcript located within intron 1 and exon 2 on chromosome 11 (ch11q23.1)^24^. Interestingly, the genes of the *miR-34* family harbour a significant conserved sequence in the putative promoter regions of *p53* binding sites contributing to its pro-apoptotic/anti-proliferative functions^25–27^. The switching of the inverse regulation between the *p53* inactivation and *miR-34a/b/c* contributes to the mesenchymal induction and, thereby, the metastatic state of the cancer cells^28^. Further, a significant association exists between *miR-34b* and phospho-Met, *p53* (phospho S392), and *MDM2* in non-small cell lung cancer (NSCLC)^29^. In a recent report, *miR-34b* was shown to mediate TGF-β1 regulation of cervical cancer cell proliferation and apoptosis^23^. In yet another study, Lu et al.^30^ have indicated the significance of *miR-34b* as a diagnostic biomarker and a potential molecular target for ovarian cancer treatment by using exosomal *miR-34b*-mediated inhibition of cell proliferation and EMT. Also, Rivas et al.^20^ have suggested the involvement of other regulatory genes in the cellular process of cervical cancer mediated by *miR-34*. While all these reports show that *miR-34b* plays a vital role in the EMT regulation of solid tumours, including cervical cancer, exploring the list of *miR-34b/gene* modulations and their underlying mechanisms would contribute to the identification of newer therapeutic targets for better disease management. To address this research gap, we applied a computational approach that includes three processes to identify targets of *miR-34b* contributing to EMT progression in cervical cancer from The Cancer Genome Atlas (TCGA), TCGA-CESC dataset, as shown in Figure 1a. Further, we validated the identified candidate *miR-34b* targets in primary CESC derived from the South Asian Indian population and cervical cancer cell lines to show the correlation between the expression of *miR-34b* and target genes. The study also revealed the repression of candidate genes/proteins in the ectopically *miR-34b* expressing stable cervical cancer cells indicating the role of *miR-34b* in cervical cancer progression. As many gene aberrations contribute to cancer progression and metastasis, advanced treatment regimens targeting multiple gene modulations acquire importance in treating cervical cancer. As EMT offers insight into novel strategies for cancer treatment, the outcome of this study could aid in identifying the multigene aberrations contributing to the advancement of biotherapy and adjuvant therapy in cervical cancer.

**Figure 1.**
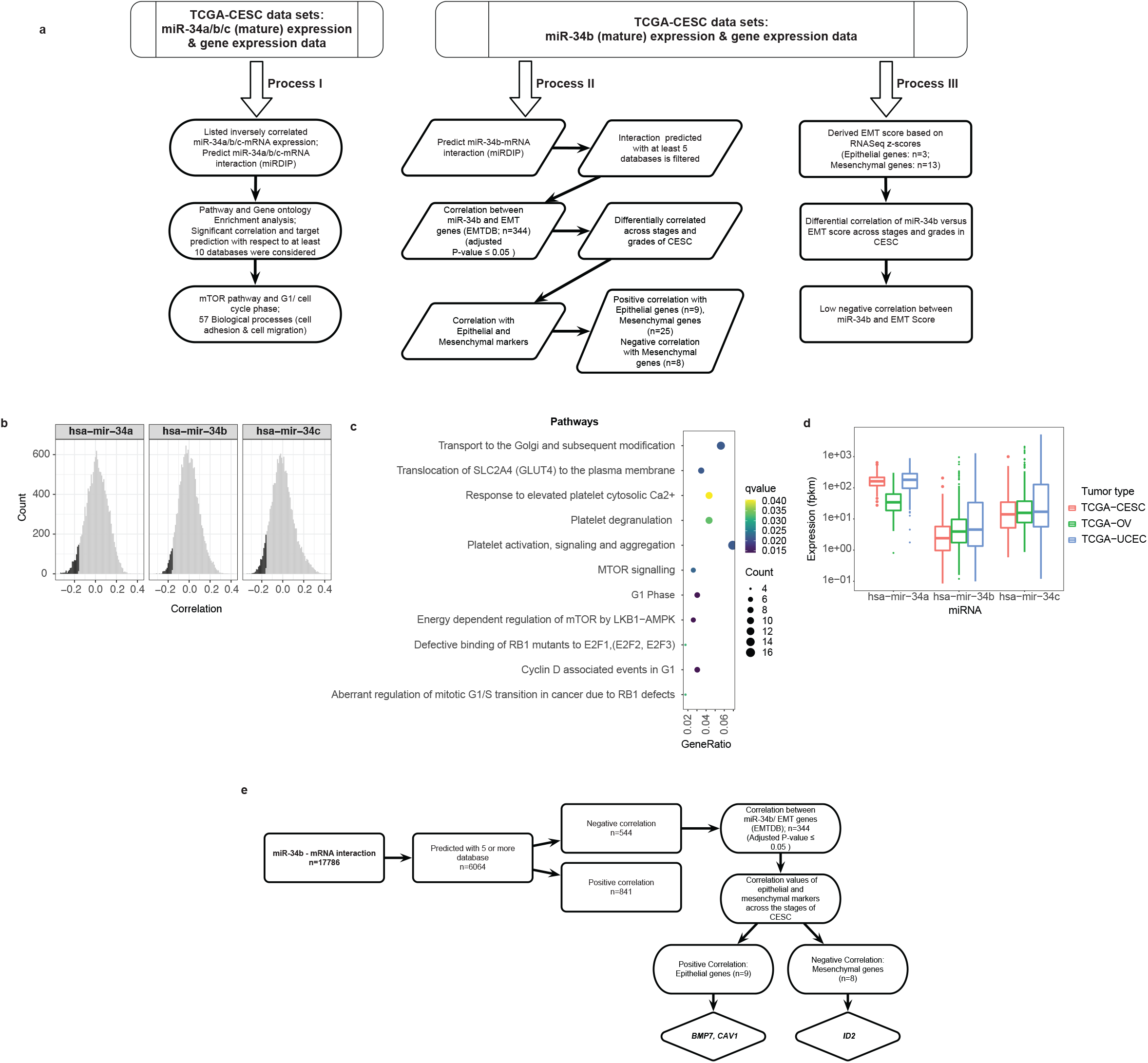
(a) Schematic representation describing the steps followed in the current study. The three processes followed in the pipeline are briefly presented, (b) Distribution of negative correlation of *miR-34* family (a/b/c) with transcripts of TCGA-CESC from TCGA datasets. The black colour in the graph indicates the cut-off value of the negative correlation considered in the study, (c) The pathway enrichment analysis using Reactome is presented. A total of 10 different significantly enriched pathways were observed (q-value ≤ 0.05). (d) Expression of *miR-34* family (*a/b/c*) in cervical cancer (CESC), ovarian cancer (OV) and uterine corpus endometrial carcinoma (UCEC) using TCGA datasets. (e) Illustration of the steps followed in the correlation analysis of miR-34b with epithelial/mesenchymal genes in cervical cancer.

## 2. Resultsand Discussion

### 2.1. *miR-34* family regulates the candidate genes in the mTOR pathway, cell cycle (*CCND2*) and cell adhesion functions (*FZD4*)

miRNA target prediction strategies alone do not indicate whether a particular pair of miRNA and mRNA interactions are biologically relevant. Therefore, a negative correlation between the team in independent TCGA-CESC cancer samples was used as a proxy for a regulatory effect (Figure 1b). This study used the mirDIP4.1 target prediction database, which includes 30 different databases for microRNA-mRNA target prediction to identify targets of the *miR-34(a/b/c*) family. A set of 23 databases had targets for all the three *miR-34* family miRNAs, and to obtain a confident set of target prediction genes, we iterated a number of databases in which a gene was predicted as a target. About 19,675 predicted genes were obtained with the cut-off of exactly five databases, of which 1,648 genes revealed a significant negative correlation after the Benjamini-Hochberg correction. However, about 4,029 predicted targets exhibited a significant negative correlation out of 27,591 genes with a cut-off of ten databases and these 4,029 genes were considered for further analysis.

Gene ontology analyses for the biological process for the above gene list identified 18 biological processes, including epithelial cell migration and adhesion (q-value ≤ 0.05, Supplementary File la). Using Reactome, a total of 10 different pathways were obtained significantly, of which the four biologically significant pathways included mTOR-associated pathways, G1 cell cycle, epithelial cell migration (ECM) and cell adhesion (q-value ≤ 0.05, Figure 1c, Supplementary File 1b). *miR-34a* was predicted to interact with the mTOR pathway genes *PRKAB2, CAB39*, and *PPM1A; miR-34b* with genes such as *PRKAA2, CAB39L* and *miR-34c* with genes like *PRKAA2, CAB39L*, and *STRADA*(Supplementary Figure S1).

We observed that *miR34a* and *miR-34c* regulated the key cell cycle genes *CCND1, CCND2, CDKN1B, E2F3, E2F5*, and *ABL1* (Supplementary Figure S2). *CCND2*, a regulator of cell cycle G1/S transition, is also expressed in cancers^31^, and miRNA-mediated repression of *CCND2* lowered the proliferation, migration and invasion properties of cervical cancer cells^32^. Our findings support the scope of studying the *miR-34a/c* and *CCND2* interactions in cervical cancer. The genes involved in ECM and *miR-34* family showed the interaction of *miR-34a* with many vital genes such as *TGFB1, TGFB2, GATA3, MAP3K3, MAP4K4* and *CDH13*. Also, *miR-34b* and *miR-34c* regulated the genes like *PTPRTG, ARSB*, and *PIK3R3* (Supplementary Figure S3). Further, the regulatory networks with the genes involved in cell adhesion function revealed a unique gene cluster for *miR-34a*. At the same time, *miR-34b* and *miR-34c* regulate *FZD4* and *LIMCH1* genes (Supplementary Figure S4). This unique cluster of *miR-34a* includes the key genes, namely *SMAD3, JAG1, ITGA6, CD44*, and *CDH13*. All these genes were widely studied for their role in the progression of various cancers and are also inversely correlated with the *miR-34* family in cervical cancer. Thus, these *mRNA/miR-34* family interactions can be further explored by the experimental validation contributing to gene therapy.

### 2.2. Epithelial genes show a positive correlation, while mesenchymal genes exhibit a negative correlation with *miR-34b* in cervical cancer

Further, the regulatory targets of *miR-34b* were specifically explored as its expression was observed to be lower than *miR-34a* and *miR-34c* in TCGA-CESC datasets (Figure 1d). Interestingly, similar results were observed in two other epithelial cancer types: ovarian cancer (TCGA-OV) and uterine corpus endometrial carcinoma (TCGA-UCEC, Figure 1d). Next, a similar strategy was applied to the *miR-34* family using the mirDIP4.1 target prediction database. About 17,786 genes out of 27,591 in 26 databases and 6,064 in more than five databases were predicted for any given interaction. Out of 6,064 genes, we observed a significant correlation of *miR-34b* with 1,385 genes, of which 544 genes showed a significant negative correlation. In comparison, 841 genes showed a significant positive correlation after the Benjamini-Hochberg correction for multiple comparisons. Out of 1,385 genes, a total of 344 genes were known to be involved in regulating the EMT pathway. Figure 1e shows the schematic representation of the steps followed in this analysis. As decreased *miR-34b* levels in the three epithelial cancers (TCGA-CESC, TCGA-OV and TCGA-UCEC) were observed, the interaction between *miR-34b* versus epithelial and mesenchymal markers was the focus of the study.

EMT is a complex process involving morphological changes and a cascade of molecular events contributing to metastasis in cervical cancer, but its regulatory networks are poorly understood. To better understand the *miR-34b* dependent regulatory network modulating the EMT process, we analysed TCGA-CESC data sets and CESC primary tumour tissues (n = 21) obtained from the Asian Indian population along with non-cancerous cervical tissues (n = 5) as control. First, qPCR analyses were performed for *miR-34b, CDH1* (epithelial signature gene) and *VIM* (mesenchymal signature gene) expressions in these select CESC and non-cancerous cervical tissues. A decreased expression in the median of *miR-34b* (Figure 2a), *CDH1* gene (Figure 2b), and increased expression in the median of *VIM* gene (Figure 2c) in Stage III of the Asian Indian population-derived CESC was observed. These results corroborate with the ones obtained from the TCGA-CESC datasets as described in the section below (*miR-34b* has a positive correlation with epithelial gene and negatively correlates with mesenchymal gene, Figures 2d, 2e).

**Figure 2.**
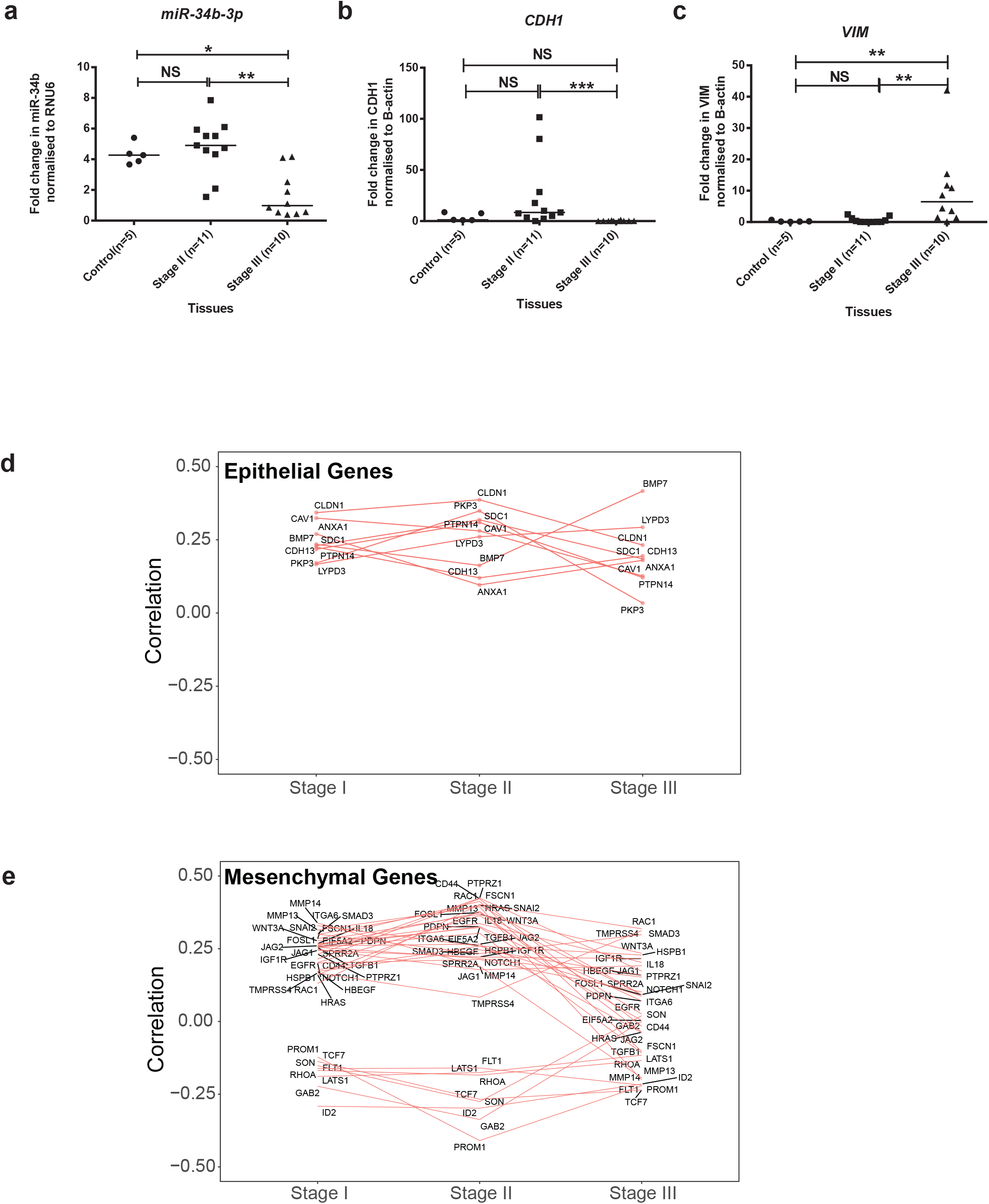
(a) Cluster graph of miRNA expression across control and Stages (II/III) in the South Asian Indian population for the *miR-34b*. (b) Cluster graph of gene expression across control and Stages (II/III) in the South Asian Indian population for the *CDH1* gene, (c) Cluster graph of gene expression across control and Stages (II/III) in the South Asian Indian population for the *VIM* gene, (d) Correlation of *miR-34b* and epithelial genes across Stages (I/II/III) of TCGA-CESC (e) Correlation of *miR-34b* and mesenchymal genes across Stages (I/II/III) of TCGA-CESC. The symbols “NS” denotes nonsignificance; * denotes p-value ≤ 0.05; ** denotes p-value ≤ 0.01 and *** denotes p-value ≤ 0.001.

Upon comparing correlation values of EMT markers across the stages of TCGA-CESC, we observed about nine epithelial markers such as *ANAX1, BMP7, CAV1, CDH13, CLDN1, LYPD3, PKP3, PTPN14* and *SDC1*, had a positive correlation across the stages of CESC (Figure 2d, Supplementary File 2). *BMP7* was known to inactivate the EMT-associated genes, thereby contributing to the reduced TGfβl-mediated cell growth and metastasis in breast cancer^33^ and a significant association of decreased *BMP7* in primary breast cancer cells with clinically overt bone metastasis was observed earlier^34^. Similarly, *CAV1* positively correlated with E-cadherin (*CDH1*) gene expression, thereby regulating EMT in gastric cancers^6,35^. Therefore, based on the high positive correlation across stages, gene expression and significant expression in the survival analysis, *BMP7* and *CAV1* were selected out of the nine genes identified for further study (Supplementary File 2).

First, we analysed the expression of *BMP7* across the TCGA-CESC data sets and found no significant change across the stages (Figure 3a). In contrast, a significant difference was observed across G2 versus G3 grades of TCGA-CESC (Figure 3d). We decided to analyse *BMP7* expression in CESC primary tumour tissues (n = 21). The results showed a significant decrease in *BMP7* expression in Stage III CESC, while no significant change was observed in Stage II CESC compared to the control cervical tissues Figure 3g). Also, a significant decrease in the survival period was observed in patients having lower *BMP7* expression (p-value = 0.017, Figure 3j). Similarly, a previous study has shown that treating cervical cancer cells with human recombinant *BMP7* resulted in tumour growth arrest by triggering hTERT gene repression^36^. Therefore, there is significant evidence for miR-34b-mediated *BMP7* modulation, which has the potential to contribute to CESC growth arrest.

**Figure 3.**
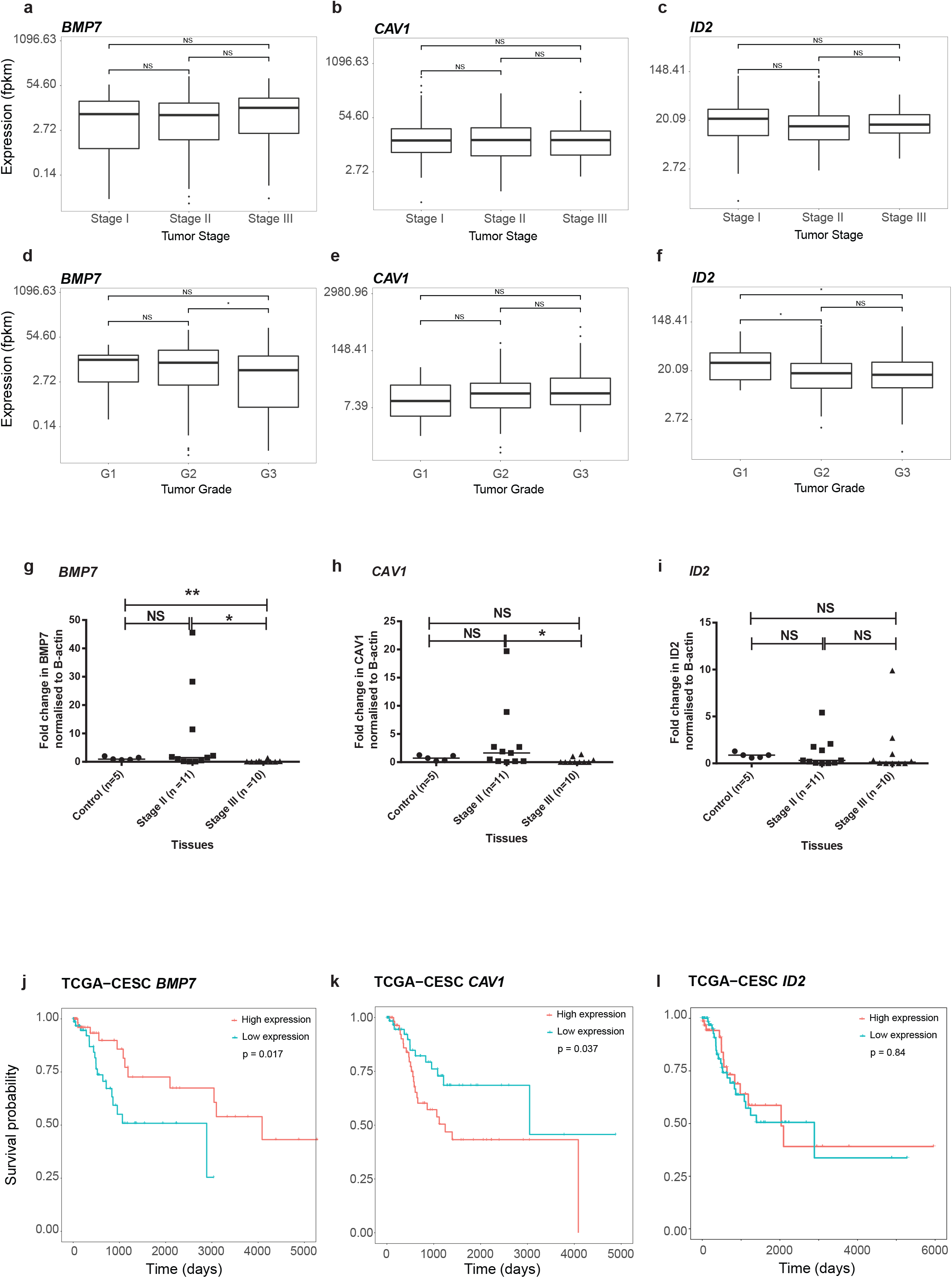
(a-f) Box plot graph across Stages (I/II/III) and grades (G1/G2/G3) of genes’ expression in TCGA-CESC; (a) *BMP7*-stages; (b) *CAV1*-stages; (c)*ID2*-stages; (d) *BMP7*-grades; (e) *C4V1*-grades; (f) *ID2*-grades; (g-i) Cluster graph of genes’ expression across control and Stages (II/III) in the South Asian Indian population: (g) *BMP7*, (h) *CAV1*, (i) *ID2;* (j-l) Survival analysis (KM plot) of TCGA-CESC based on their median genes expressions of (j) *BMP7*, (k) *CAV1*, (l) *ID2*. The symbol “NS” denotes non-significance; * indicates p-value ≤ 0.05, and ** denotes p-value ≤ 0.01.

Next, we analysed the expression of the *CAV1* gene across the TCGA-CESC datasets; the results showed an insignificant change across stages and grades of TCGA-CESC datasets (Figures 3b, 3e). Further, qPCR analysis of *CAV1*gene expression in CESC revealed a significant decrease in Stage III than Stage II cervical cancer tissues (Figure 3h). A recent study on cervical cancer cell lines has reported a decreased level of *CAV1* gene expression, and the m/7?-96-mediated restoration of the *CAV1* has reduced cell proliferation, migration, and invasion^37^. Also, other studies have indicated the tumour suppressor role of *CAV1* in various cancers, including cervical cancer^38–40^. However, in the present study, a significant decrease in survival period was observed in patients having higher *CAV1* gene expression in TCGA-CESC datasets (Figure 3k). So, as the current study indicates a positive correlation between *miR-34b* and *CAV1*, further experimental validation of *miR-34b* mediated *CAV1* modulation would aid in understanding their role in CESC tumour management.

Following epithelial markers, we observed a significant negative correlation of eight mesenchymal markers with *miR-34b*, namely *RHOA, GAB2*, SON, *PR0M1, TCF7, ID2, FLT1*, and *LATS1* (Figure 2e, Supplementary File 2). All genes, except *PR0M1*, had the MRE in their 3’UTR for *miR-34b*, indicating that they may be directly regulated by *miR-34b*.

Among these eight mesenchymal markers, the *ID2* gene, which had a relatively high negative correlation (r^2^= −0.27; p-value < 0.001) than other genes, was further analysed to understand its significance in CESC tumour progression. *ID2* has been reported as an EMT attenuator in triplenegative breast cancers^41^ and multiple myeloma^42^. Also, *ID2* has been stated as a potential therapeutic target in salivary gland carcinoma where its repression has reduced Vimentin, N-cadherin, and Snail with induction of E-cadherin expression leading to a more differentiated phenotype^43^. In the present study, an insignificant change in *ID2* gene expression was observed between the stages (Figure 3c), while a significant difference was observed between the grades of TCGA-CESC datasets (Figure 3f). Further, the qPCR analysis in primary cervical cancer tissues of the South Asian Indian population showed an insignificant change in *ID2* gene expression across stages of CESC (Figure 3i). However, a minor difference in the patient’s survival between the low and high *ID2* gene expressions was also observed (Figure 3l). Therefore, these results indicate sufficient evidence for exploring further the *miR-34b*-mediated *ID2* modulation and their potential to contribute to tumour regression in CESC.

In addition to mesenchymal markers showing a negative correlation, a total of 25 mesenchymal markers had a significant positive correlation (r^2^= 0.2-0.25, p-value ≤ 0.01), including genes such as *HSPB1, IGF1R, SNAI2*, and *SMAD3* (Figure 2e) and this observation can be further studied with experimental validation on its association with *miR-34b*.

To further understand the relationship between putative EMT markers regulated by *miR-34b*, we analysed EMT scores, indicating the distribution of hybrid Epithelial/Mesenchymal (E/M) cells in the primary tumours and aiding in identifying the putative therapeutic targets and therapy resistance in cervical cancer. Previous studies have discussed the various methods to derive EMT score^44^. A differential correlation analysis of *miR-34b* expression with EMT scores across the stages and grades of TCGA-CESC datasets was performed in the present study (Supplementary File 3). As this analysis precisely involves the quantitative measurement of EMT signature genes, its outcome would aid us in identifying relevant genes regulated by *miR-34b* involved in EMT. The overall correlation revealed a significant low negative correlation (r^2^= −0.07) between *miR-34b* and EMT score in TCGA-CESC. Also, a significant low negative correlation was observed across stages (Figure 4a) and a varied correlation across grades (Figure 4b). These results indicate that if there is an increase in *miR-34b* expression, we may obtain a more negative EMT score, showing an increase in epithelial signature genes than the mesenchymal genes. So *miR-34b* can be further explored for its potential to regulate the EMT signature genes, thereby improving the management of CESC metastasis.

**Figure 4.**
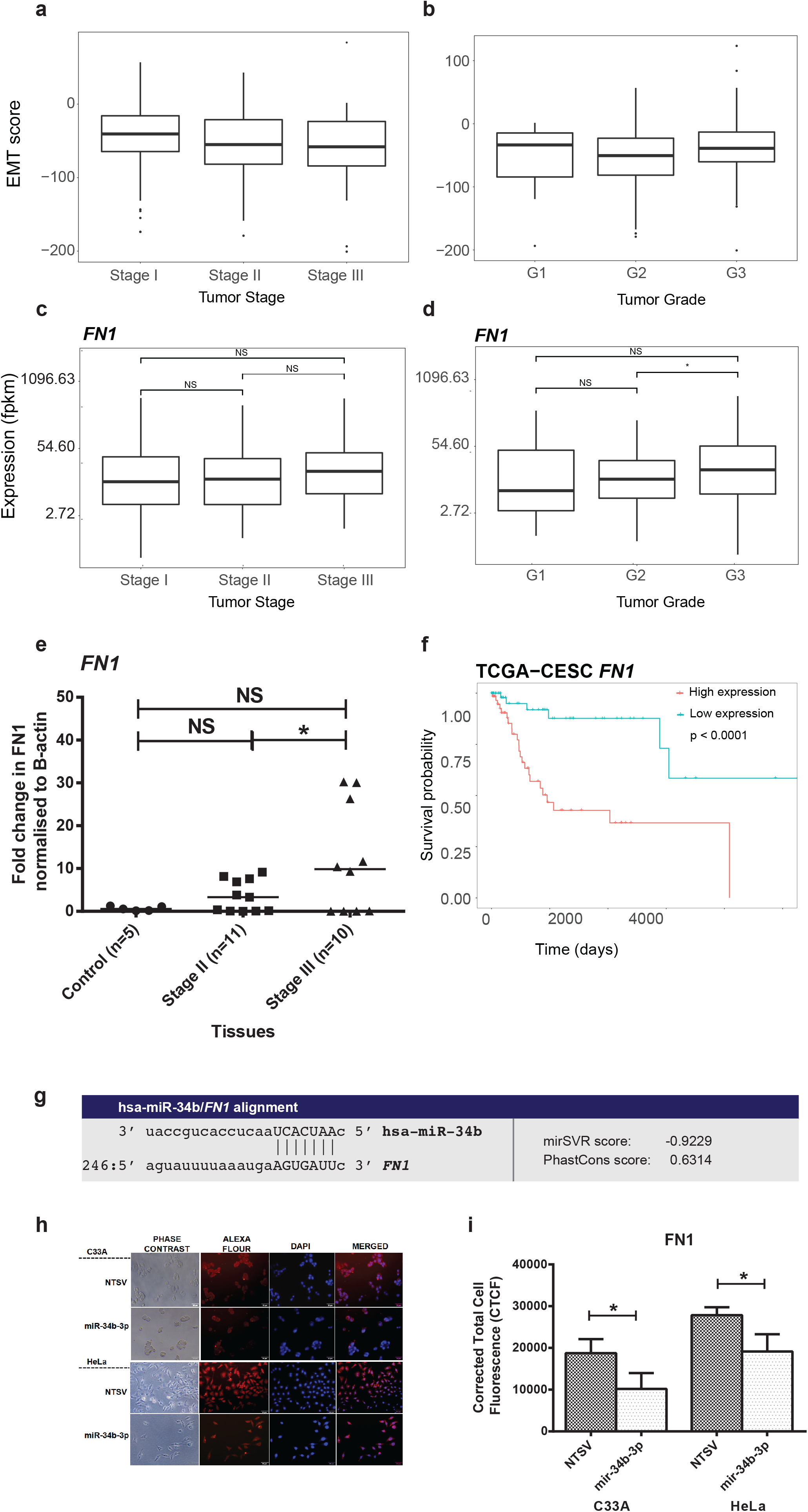
(a) Correlation analysis between *miR-34b* and EMT score in TCGA-CESC across Stages (I/II/III), (b) Correlation analysis between *miR-34b* and EMT score in TCGA-CESC across grades (G1/G2/G3), (c) Box plot graph across Stages (I/II/III) of *FN1* expression in TCGA-CESC, (d) Box plot graph across grades (G1/G2/G3) of *FN1* expression in TCGA-CESC, (e) Cluster graph of gene expression across control and Stages (II/III) in the South Asian Indian population, (f) Survival analysis (KM plot) of TCGA-CESC based on *FN1* median expression, (g) Sequence alignment analysis between *miR-34b* and 3’UTR of *FN1*, (h) Microscopic observation of immunofluorescence in transfected control (NTSV) and *miR-34b-3p* expressing cervical cancer cells (C33A and HeLa), (i) Bar graph of invasion assay in transfected control (NTSV) and *miR-34b-3p* expressing cervical cancer cells (C33A and HeLa). The black shaded bar shows the *miR-34b-3p/gene* expressions in cervical cancer cells carrying non-target sequence vector (NTSV), and the grey shaded bar shows the *miR-34b-3p/gene* expressions in cervical cancer cells carrying *miR-34b-3p* expressing vector. The symbol “NS” denotes non-significance; * indicates p-value ≤0.05.

Based on the correlation analysis between *miR-34b* and these EMT signature genes, we found Fibronectin 1 (*FN1*), a mesenchymal marker, had a significant high negative correlation (r^2^= 0.0028). *FN1* expression in TCGA-CESC datasets showed an insignificant difference in expression across stages (Figure 4c) while a significant change across the grades (Figure 4d). Also, the qPCR analysis in primary cervical cancer tissues of the South Asian Indian population showed a significant change in *FN1* gene expression across stages of CESC (Figure 4e). In addition, a significant decrease in the patient’s survival with high *FN1* expression was also observed (Figure 4f). Also, *FN1* has MRE in its 3’UTR for *miR-34b* (Figure 4g), indicating that it is inversely regulated by *miR-34b*. Further, the deregulation of *FN1* protein by *miR-34b* was analysed by immunofluorescence stain. Here, a significant decrease of Corrected Total Cell Fluorescence (CTCF) in the presence of *miR-34b* expressing cervical cancer cells compared to the non-target sequence vector (NTSV) transfected control cells (Figure 4h, 4i) indicated the repression of *FN1* protein expression. These results prove that *FN1* could be a potential target of *miR-34b* in CESC.

To ascertain the select candidate gene expression in the presence of *miR-34b*, the cervical cancer cells (C33A and HeLa) were stably expressed with the *miR-34b-3p* expression construct. The stable cells expressed *miR-34b-3p* up to 4-5 folds in cervical cancer cells (C33A and HeLa) after 48h of induction (Figure 5a). At this time point, a significant increase in the *CDH1* (Figure 5b) and a decrease in *VIM* (Figure 5c) compared to the NTSV transfected control cells was observed. This change in *CDH1* and *VIM* gene expression was also supported by their protein expressions (Figure 5h). Thus, the increase of epithelial marker (*CDH1*) and a decrease of mesenchymal marker (*VIM*) in the presence of *miR-34b* indicate its association with EMT regulation of cervical cancer cells.

**Figure 5.**
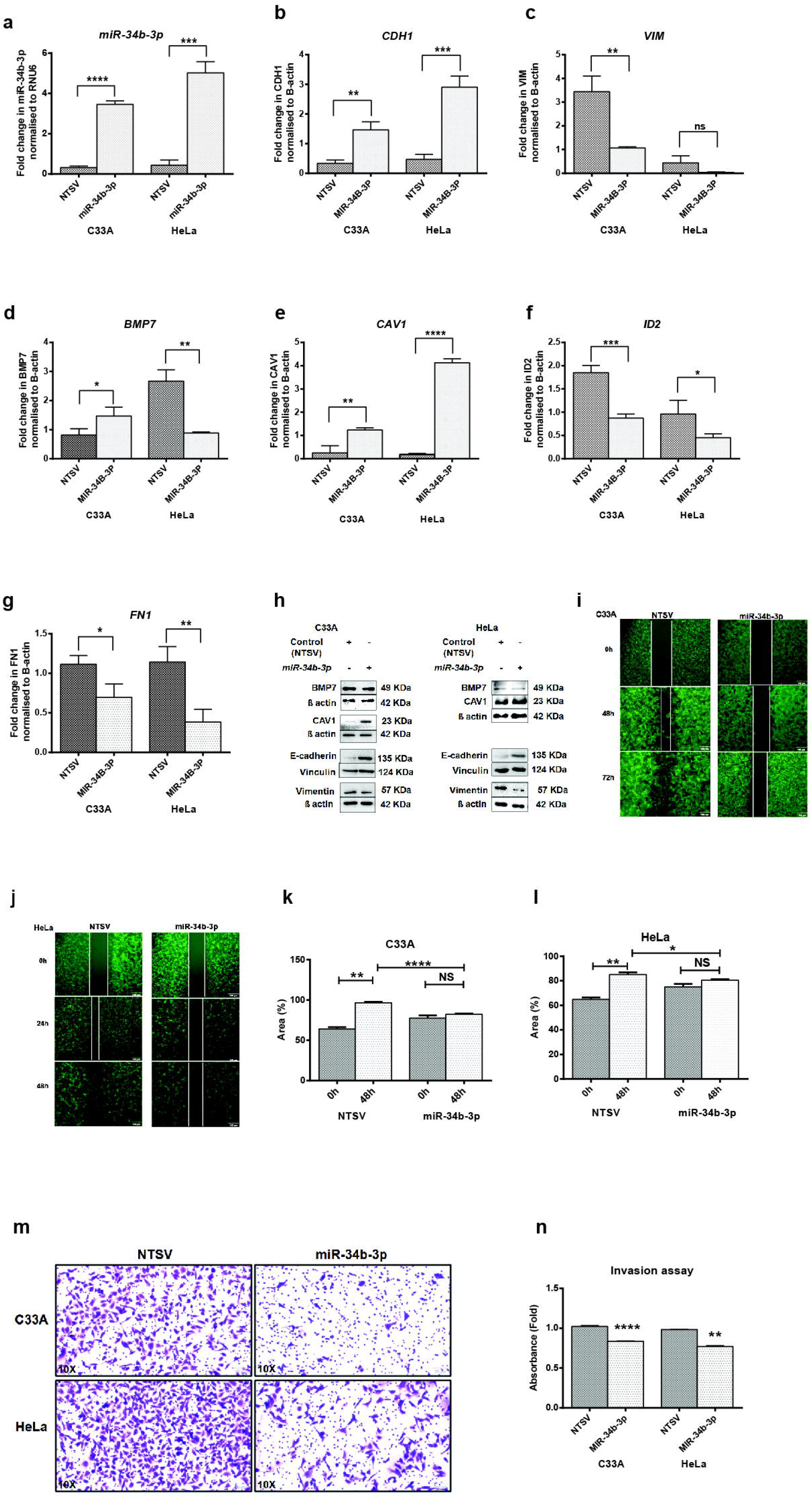
(a-g) Bar graph of *miR-34b-3p/gene* expressions in *miR-34b-3p* expressing cervical cancer cells (C33A and HeLa). (a) *miR-34b-3p*, (b) *CDH1*, (c) *VIM*,(d) *BMP7*, (e) *CAV1*, (f) *ID2*, (g) *FN1*, (h) Western analysis of *BMP7, CAV1*, E-cadherin, *VIM*, Vinculin and β-actin proteins in transfected control (NTSV) and *miR-34b-3p* expressing cervical cancer cells (C33A and HeLa), (i and j) Microscopic observation of migration assay in transfected control (NTSV) and *miR-34b-3p* expressing cervical cancer cells at 0h, 48h and 72h (i: C33A and j: HeLa), (k) and (l) Bar graph of migration assay in transfected control (NTSV) and *miR-34b-3p* expressing cervical cancer cells at 48h (k:C33A and l: HeLa), (m) Microscopic observation of invasion assay in transfected control (NTSV) and *miR-34b-3p* expressing cervical cancer cells at 48h (C33A and HeLa), (n) Bar graph of invasion assay in transfected control (NTSV) and *miR-34b-3p* expressing cervical cancer cells at 48h (C33A and HeLa). The black shaded bar shows the *miR-34b-3p/geπe* expressions in cervical cancer cells carrying non-target sequence vector (NTSV), and the grey shaded bar shows the *miR-34b-3p/geπe* expressions in cervical cancer cells carrying *miR-34b-3p* expressing vector. The symbols “NS” denotes non-significance;* denotes p-value ≤ 0.05; ** denotes p-value ≤ 0.01;*** denotes p-value ≤ 0.001, and **** denotes p-value ≤ 0.0001.

Further, the *miR-34b-3p* expressing cervical cancer cells revealed a significant increase of *BMP7* gene expression in C33A cells (1.5-fold) while a decreased expression in HeLa cells (0.9-fold) compared to the transfected control cells (Figure 5d). A similar result was also significantly observed for the Bmp7 protein expression (Figure 5h). The difference in the cell lines phenotype may contribute to the variation of *BMP7* expression. The observed decrease in *BMP7* gene expression in the HeLa cells corroborated an earlier report where the *BMP7* knockdown inhibited EMT by inducing E-cadherin and decreasing Vimentin^45^. However, the increased *BMP7* gene levels observed in C33A trigger further evaluation of the molecular insights as *BMP7* was reported to inactivate the EMT-associated genes resulting in reduced TGFâ1-mediated breast cancer cell growth and metastasis^33^. Next, a significant increase of *CAV1* at the mRNA level (Figure 5e) and protein levels (Figure 5h) was observed. As *CAV1* is reported for tumour suppressor activity in other cancers, its modulation by *miR-34b* in the present study may contribute to the increased tumour suppressor activity in cervical cancer cells. Next, a significant decrease of *ID2* (Figure 5f) was observed in the cervical cancer cells in the presence of *miR-34b. ID2*, being reported as an EMT attenuator in other cancers, its suppression by *miR-34b* in cervical cancer cells may contribute to better tumour management. Also, the decrease of *FN1* mRNA levels in the cervical cancer cells expressing *miR-34b-3p* strongly indicates the regulation of *FN1* by *miR-34b-3p* (Figure 5g).

Thus, the inverse changes of select candidate gene expressions in the ectopically *miR-34b-3p* expressing cervical cancer cells (Figure 5) to that of primary cervical cancers (Figures 2, 3) indicate their association. In addition, the regulation of these genes or proteins resulting in a significant decrease of migration and invasion properties of *miR-34b-3p* expressing cervical cancer cells suggests that the select candidate genes are potentially modulated by *miR-34b* to contribute to tumour regression in CESC (Figures 5i-5n). However, further experimental validation needs to be carried out to understand the regulation of these candidate genes by *miR-34b* in a cascade.

## 3. Conclusions

A wide range of genetic aberrations regulated by the *miR-34* family involving EMT in cervical cancer has been indicated in this study. This study reveals the evidence for regulating candidate genes in the mTOR pathway, cell cycle (*CCND2*) and cell adhesion functions (*FZD4*) by the *miR-34* family. In addition, the correlation of *miR-34b* with the signature genes involved in EMT modulation has identified the new putative targets, namely *CDH1, BMP7, CAV1*, and *ID2*. Further, using EMT scores, *we* show that *FN1* could be a direct target regulated by *miR-34b*. Following this identification, the experimental validation using gene expression studies in ectopically expressed *miR-34b-3p* cervical cancer cells proves their association with each other. In the present study, an integrated approach was used to reveal the novel targets for the *miR-34* family, especially *miR-34b* and experimentally showed the relationship between *miR-34b* and its target genes in CESC. Also, the derived observations depend entirely on the absolute gene expressions of primary TCGA-CESC datasets, as they are robust to be carried out for further experimental validation. In addition, these results were supported by the evidence obtained from a small subset of the Asian Indian population. Finally, the results obtained from this integrated analysis provide convincing evidence for the role of *miR-34b* in EMT modulation by regulating the candidate signature genes (*BMP7, CAV1, ID2*, and *FN1*) and pave the way for a more detailed study to understand their interactions and help in epithelial cancer cell metastasis management of cervical cancer.

## 4. Methods

### 4.1. Identify miRNAs whose expression is correlated with cervical cancer

Previously generated miRNA and mRNA expression levels by RNASeq within TCGA Research Network (http://cancergenome.nih.gov/) were used in the study. The processed count and raw read data were downloaded for cervical cancer from TCGA Genomics Data Commons (GDC) portal (https://gdc.cancer.gov/).TCGA-CESC expression data were derived from 307 cervical carcinomas (epithelial squamous cell and endocervical adenocarcinoma) and two non-cancerous cervical epithelial cells. The samples were identified using the TCGA barcode associated with each sample. The clinical metadata files such as tumour stage, patient’s age, gender, and survival statistics of each TCGA-CESC sample were downloaded through the GDC Application Programming Interface (API). Supplementary Table 1 summarises the clinical metadata of all the CESC samples (n = 307) from the TCGA project. Gene expression (Fragments per Kilo-base of exon per Million mapped, FPKM) and π⋂iRNA expression (Counts per Million, CPM) quantification data files available from cervical epithelial squamous cell carcinoma (TCGA-CESC) were utilised from TCGA.

### 4.2. Identify potential gene targets of the *miR-34* family and *miR-34b* through TCGA cervical cancer gene expression dataset

RNA-sequence expression data generated by the TCGA project available from the Genomics Data Common repository (https://portal.gdc.cancer.gov)was used. A TCGA file list, ‘TCGA-CESC sample sheet’, containing information such as sample ID and file ID for each CESC sample, was also downloaded by querying the GDC API. This file list sample sheet was used to download the files in Ras input for the pipeline in R. The library sizes for gene expressions and miRNA expressions were 60,483 and 1,881, respectively. Expression criterion of genes with FPKM ? 1.0 and miRNA with CPM ? 1.0 across all samples were applied as described earlier^46^. The cut-off values represent a mean frequency ?l in a million reads mapped for microRNA and a mean frequency ?l in a million reads after transcript length normalisation for mRNA. There were 14,509 genes and 407 miRNAs for which the expression was above the cut-off in TCGA-CESC datasets.

The pipeline was built using R version 3.6.1, which includes data downloading, correlation analysis, target prediction database lookup, and functional enrichment. The instructions for running the pipeline are presented as R script and can be accessed from the GitHub repository (https://github.com/HimanshuLab/miR-34-in-pan-Gvnaecological-Cancers). The entire mirDIP4.1 database was downloaded and used to compare the target prediction across 30 individual resources^47^. The mirDIP4.1 database has been used to compare 26 microRNA-mRNA target prediction tools databases. These databases cover a wide variety of target prediction strategies, including seed region matching, evolutionary conservation of regulatory region on mRNA, binding energy, site accessibility, CLIP experimental evidence, and machine learning methods. DIANA-TarBase (v8.0; http://www.microrna.gr/tarbase) was used to lookup experimentally validated π⋂icroRNA-mRNA interactions^48^. Additionally, TargetScan (v7.0; www.targetscan.org), MiRanda-miRSVR (v3.3a; http://www.microrna.org) and DIANA-microT-CDS (v5.0; http://diana.imis.athenainnovation.gr/DianaTools/index.php) databases were utilised^49,50^. Only predictions categorised as either good or better according to the miDIP database were used. As microRNA target prediction databases are known to have many false positives, we have considered microRNA-mRNA interactions predicted to occur in at least five databases as a weak indicator and being predicted in over ten bases as a strong indicator.

### 4.3. miRNA:mRNA correlation analysis with epithelial and mesenchymal genes

As one of the key functions of miRNAs is to potentially destabilise the mRNAs, we focussed on the significant miRNA-mRNA negative correlation as an indication of a regulatory effect in subsequent analysis. Spearman rank correlations were calculated for *miR-34(a/b/c*) family versus gene expression using the rcorr function from the Hmisc R package. Hmisc::rcorrfunction also provides asymptotic p-values for the correlations, corrected for multiple comparisons using the Benjamini-Hochberg method. *miR-34(a/b/c*) family and gene pairs were only considered if they were anticorrelated (Spearman correlation; *r*^2^ <0) and if the corrected p-value was significant (p-value < 0.01).

The EMT genes from the first literature-based database^51^ for EMT human genes (dbEMT) was used in this study. The correlations of *miR-34b* with EMT genes were generated with adjusted p-values using the Benjamini-Hochberg method. The correlations with a p-value ≤ 0.01 were selected and analysed across stages/grades of CESC. The analysed results are presented in Supplementary File 1 and derived a co-expression network using Cytoscape v3.7.1. The International Federation of Gynaecology and Obstetrics (FIGO) staging system based on clinical examination and specific procedures (imaging) was used in the study^52^. The tumour stage of each sample was classified into Stages I, II, III, and IV by binning the various subtypes in each stage. Independent of the staging system, the histopathologic parameter based on the tumour cell differentiation was also used to study the candidate gene expression levels. The tumour grade of each sample was classified as GX (grade cannot be assessed), G1 (well-differentiated), G2 (moderately differentiated) and G3 (poorly or undifferentiated)^52^. The candidate gene expressions were checked across Stages I/II/III and grade G1/G2/G3 as the increase in stage and grade directly proportionate to the metastasis and poor prognosis of CESC.

### 4.4. Calculation of EMT scores from TCGA cervical cancer gene expression dataset

EMT score is a quantitative measure of the sample characteristic on the Epithelial-Mesenchymal spectrum. The EMT score for each sample was derived by subtracting the mean gene expression of epithelial markers from the mean gene expression of mesenchymal markers. TCGA-CESC datasets were used to derive the gene expression RNA sequencing z-scores for the epithelial markers (n = 3) and mesenchymal markers (n = 13)^53^. The epithelial markers were E-Cadherin (*CDH1), DSP*, and *TIPI;*while the mesenchymal markers were Fibronectin (*FN1*), Vimentin (*VIM*), N-Cadherin (*CDH2*),Integrin beta 6 (*ITBG6*), Forkhead box protein C2 (*FOXC2*), Matrix metalloproteinase 2 (*MMP2*), Snail family transcriptional repressor 1 (*SNAI1*), Matrix metalloproteinase 3 (*MMP3*), Snail family transcriptional repressor *2 (SNAI2*), Matrix metalloproteinase 9 (*MMP9*), Twist-related protein *l(TWISTl*), SRY-Box transcription *ĩactorlO(SOXl0*), and Goosecoid homeobox (GSC). The z-scores for three mesenchymal genes (*ITBG6, SOX10*, and *GSC*) were not obtained as no data was available in TCGA-CESC datasets. A positive EMT score was thus associated with the mesenchymal cell type, while a negative score was associated with the epithelial cell type. A subgroup analysis using grades and stages of CESC for correlation between *miR-34b* expressions and EMT score was also derived. A negative correlation indicates that as the gene’s expression increased, the sample was more likely to take on epithelial characteristics (a more negative EMT score). The derived EMT scores for each sample of TCGA-CESC are given in Supplementary File 3.

### 4.5. Gene ontology and pathway analysis

Functional enrichment was done using Reactome Pathways (https://reactome.org/)using the R/Reactome PA package. Ontology enrichment was done using gene ontology (GO) using the R/cluster Profiler package. The Reactome and gene ontology results are filtered using q-values calculated using adjusted p-value and false discovery rate (FDR). A cut-off q-value of 0.05 was used. The obtained results are presented in Supplementary File 1. The resulting network was plotted using Cytoscape v3.7.1 network analysis of gene interactions between the genes of interest was done using the STRING-db v11.O webpage (https://string-db.org/cgi/).

### 4.6. Survival analysis

Survival Analysis was done using Kaplan-Meier (KM) plots using the R packages survminer, survival, and RTCGA.clinical. The survival probability was compared between the samples in the upper and lower quartiles of expression for a given gene, and the adjusted p-values were derived using the Benjamini-Hochberg method.

### 4.7. Primary cervical cancer tumour tissues of the South Asian population

Twenty-one primary cervical cancer tumour tissues and five normal cervical tissues collected from the South Asian population reported at the regional institutes were included in the study. These primary tumour tissues (n = 21) include fresh human cervical cancer tissues (n= 10) and formalin-fixed paraffin-embedded tissues (n = 11) with a median age of 55. Ten of these tumour samples were Stage II, while eleven were Stage III (Supplementary Table 2, Supplementary Figure S5). Respective non-cancerous cervical biopsy tissues were used as normalised controls (n = 3).

### 4.8. *miR-34b* over-expression in cervical cancer cell lines

The shMIMIC inducible Lentiviral microRNA, tetracycline-inducible expression system (Tet-On 3G) with TurboGFP as reporter and mCMV as promoter for both *hsa-miR-34b-3p* (MIMAT0004676; 5’CAAUCACUAACUCCACUGCCAU3’; Cat# GSH11929-224638820; Dharmacon™) and for nontargeting sequence control (Cat# VSC11651; Dharmacon™) was purchased. The packaging lentiviral particle, transduction and puromycin selection were carried out per the manufacturer’s protocol. The stable cell lines were established for HeLa (RRID: CVCL_0030) and C33A (RRID: CVCL_I094) cervical cancer cell lines (*Mycoplasma sps* free). Both the cell lines were maintained in a growth medium containing Dulbecco’s Modified Eagle’s Medium (DMEM, Life Technologies, Carlsbad, CA, USA); 10% FBS (Life Technologies) and antibiotics (l00⍰U⍰ml^-1^ penicillin and l00⍰μg⍰ml^-1^streptomycin) at 37⍰°C with a humidified atmosphere of 5% CO_2_. Both cell lines have been authenticated using STR profiling within the last three years.

### 4.9. Immunofluorescence

About 1 x 10^5^ stable cells of C33A/HeLa lines were seeded in a 12-well cell culture plate and incubated overnight at 37⍰°C with a humidified atmosphere of 5% CO_2_. Following incubation, doxycycline (C33A:750ng/ml/ HeLa: 800ng/ml was added and incubated for 48h at 37⍰°C with 5% CO_2_. Next, the cells were fixed with chilled methanol (10 min), washed with ice-cold 1XPBS (3 times) and incubated with 1% BSA in PBST (PBS+ 0.1% Tween 20). The cells were incubated with the diluted FN1 antibody (1:50, Cat# SC-9068, Santacruz) in a humidified chamber overnight at 40⍰°C. Then, the cells were probed with diluted Alexa flour conjugated secondary antibody (Alexa Flour™ 568, Cat# A-11011, Invitrogen) for 1h in the dark. The cells were intermittently washed three times with chilled 1XPBST. Finally, the cells were stained with DAPI (DNA stain) for 1 min, and the cells were imaged using an Olympus IX-73 microscope and analysed using ImageJ software. The corrected total cell fluorescence (CTCF) was calculated using the equation, *CTCF = Integrated Density — (Area of selected cell × Mean fluorescence of background readings*), described by Jakic et al.^54^

### 4.10. RNA isolation, cDNA synthesis, and real-time quantitative PCR (qRT PCR)

The total RNA is extracted from the stable cell lines (C33A/HeLa) using a trizol reagent (RNA Iso Plus reagent, Takara) and from paraffin-embedded cervical cancer tumour tissue sections using the RNeasy FFPE kit (catalogue number 73504, Qiagen) according to the manufacturer’s protocol. The total RNA was quantified using a NanoDrop spectrophotometer (ND2000, Thermo Scientific, MA, USA).and converted to cDNA using a Primescript RT reagent kit (Cat# RR037A, DSS Takara Bio India Private Ltd) as per the manufacturer’s instructions. The stem-loop primers for miRNA cDNA conversion and oligodT primers for mRNA cDNA conversion were used. The converted cDNA was amplified using gene-specific primers and TB Green^®^ Premix Ex Taq^™^ II (Cat# RR820A; DSS Takara Bio India Private Ltd) to quantify the expression of the genes using a real-time PCR system (Applied Biosystem Quant Studio 7 Flex) with the following protocol (Denaturation: 95⍰°C/5 min; Amplification for 40 cycles: 95⍰°C/30 sec, 60⍰°C/45 sec, 72 °C/30 sec). The primers used in this study are given in Supplementary File 4. The relative C_t_ (cycle threshold) quantification of genes in tumour and normal cell lines was analysed for expression analysis. All samples were assayed in triplicates. The unpaired t-test was performed using GraphPad Prism 6.

### 4.11. Western Analysis

Protein isolation was carried out by resuspending the PBS-washed cells in radioimmunoprecipitation assay (RIPA) buffer and vortexing intermittently. The protein lysate was subjected to high-speed centrifugation, resolved by sodium dodecyl sulphate-polyacrylamide gel electrophoresis (SDS-PAGE) and transferred onto a polyvinylidene difluoride membrane (Cat# 1620177, BioRad). Following the protein transfer, blots were blocked using bovine serum albumin (5%) and probed with the primary antibody at 4 °C overnight and on the next day with an HRP-linked secondary antibody for 1h at room temperature. The primary antibodies against *BMP7* (Cat# A0697, ABclonal), *CAV1* (Cat# A1555, ABclonal), E-cadherin (Cat# SC-8426, G-10, Santacruz), Vimentin (Cat# SC-33322, RV-202, Santacruz), Vinculin (Cat# 4650S, Cell signalling technologies), and β-actin (Cat# A2228, Ac-74, Sigma Aldrich) proteins were used in the study. Intermittent washing of the blots was carried out using Tris-buffered saline containing 1% Tween-20 (1x TTBS). Following the incubation of antibodies, the immunoblot was subjected to imaging using an Enhanced Chemiluminescence Kit (Cat# 1705061, Bio-Rad) and ChemiDoc (BioRad).

### 4.12. Migration and invasion assay

About 2 × 10^6^ pre-induced cells were plated in each well of Culture-Insert 3 Well in μ-Dish 35 mm (Cat# 80366, ibidi). After overnight incubation, a fresh medium containing 5 μg/ml mitomycin-c and doxycycline was added, and the three-well silicone insert was removed to form two defined cell-free gaps. The migration of cells was imaged using an Olympus IX-73 microscope at 0h, 24h, and 72h of incubation. The percentage of area covered by the cells was analysed using ImageJ software.

Trans-well cell culture inserts coated with GelTrex (24-well inserts: Cat# 140629, GelTrex: Cat# A15696-01, Thermo Fischer Scientific) were used for the study. About 50 × 10^3^ pre-induced cells were seeded on the upper chamber of the cell culture insert with growth media containing 1% FBS, while the lower one had a complete medium (10% FBS). Cells were incubated to migrate for 48 h. After incubation, the cells were fixed with 100% chilled methanol, stained with 0.1% crystal violet solution, and imaged using an Olympus IX-73 microscope. After imaging, the insert membranes carrying cells were destained in 10% acetic acid solution, and absorbance was measured at 595 nm to quantify migrated cells^55^.

## Ethical approval

The study followed the protocol as per the Declaration of Helsinki. The Institutional Ethics Committee of Sri Ramachandra Medical College and Research Institute (SRMC&RI), Chennai and the Institute of Obstetrics and Gynaecology, Chennai (No:36433/E1/2008-1) approved this study.

## Informed consent

All individual participants of the study have provided informed consent.

## Authors’ contributions

The authors confirm their contribution to the paper as follows: VN, AX, HS, DK: Study conception and design; VN, AX, KJS, SS: Data collection; SS, VN: Interpretation of primary tumour samples and sample selection; VN, AX, HS, DK: Analysis and interpretation of results; VN, AX, HS, DK: Draft manuscript preparation: All authors reviewed the results and approved the final version of the manuscript.

## Acknowledgements

The authors would like to thank the Indian Institute of Technology Madras and the Indian Council of Medical Research for the postdoctoral fellowship of Nalini Venkatesan (Project no. 2019-4566/CMBBMS); Dr A.K. Munirajan, Professor and Head, Department of Genetics, University of Madras, Chennai, for the cDNA of CESC from the South Asian Indian population.

## Availability of data and materials

Supplementary data is available for the manuscript. The TCGA data is publicly available. The instructions for running the pipeline are presented as R script and can be accessed from the GitHub repository (https://github.com/HimanshuLab/miR-34b-in-pan-Gvnaecological-Cancers).

This manuscript is available in the bioRxiv preprint server https://doi.org/10.1101/2021.09.02.458804.

## Supplementary Tables

**Supplementary Table 1**

Summary of the clinical metadata, including median age, five-year survival statistics, tumour stage, and grade associated with each sample.

**Supplementary Table 2**

Details of the non-cancerous cervical tissue and primary CESC, of which *miR-34b-3p/gene* expression data presented in the study, are listed.

## Supplementary Files

**Supplementary File**1: Results from functional enrichment analysis - GO (1a) and Reactome (1b).

**Supplementary File 2:**Results obtained from the correlation analysis between EMT genes and *miR-34b*.

**Supplementary File 3:**Derived EMT scores of TCGA-CESC.

**Supplementary File 4:**Primer details used in the study.

## Supplementary Figures

Network analysis was derived using STRING-db v11.0 for pathways, networks and functions below. The gene names in each figure are coloured based on the intensity directly proportional to their expression, as shown in the colour bar.

**Supplementary Figure S1:**mTOR pathway

**Supplementary Figure S2:**G1/Cell cycle regulation

**Supplementary Figure S3:**Epithelial cell migration (ECM)

**Supplementary Figure S4:**Cell adhesion

**Supplementary Figure S5:**Representative microscopic images of well-differentiated, moderately differentiated and poorly differentiated CESC (Haematoxylin and Eosin stain) included in the study are presented.

