## Supplementary File for "*miR-34b* associates with epithelial-mesenchymal transition and regulates *BMP7, CAV1, ID2* and *FN1* in human cervical cancer"

**SUPPLEMENTARY TABLES**

**Supplementary Table 1:** Summary of the clinical metadata, including median age, five-year survival statistics, tumour stage, and grade associated with each sample.

**Supplementary Table 2:** Details of the non-cancerous cervical tissue and primary CESC, of which *miR-34b-3p*/gene expression data presented in the study, are listed.

**SUPPLEMENTARY FIGURES**

Network analysis derived by using STRING-db v11.0 for pathways, networks and functions given below. The gene names in each figure are coloured based on the intensity directly proportional to its expression, as shown in the colour bar.

**Supplementary Figure S1**: mTOR pathway
**Supplementary Figure S2**: G1/Cell cycle regulation
**Supplementary Figure S3**: Epithelial cell migration (ECM)
**Supplementary Figure S4**: Cell adhesion

**Supplementary Figure S5**: Representative microscopic images of well-differentiated, moderately differentiated and poorly differentiated CESC (Haematoxylin and Eosin stain) included in the study are presented.

**SUPPLEMENTARY FILES**

**Supplementary File 1**: Results obtained from functional enrichment analysis (a) GO, (b) Reactome.
**Supplementary File 2**: Results obtained from the correlation analysis between EMT genes and *miR-34b*.
**Supplementary File 3**: Derived EMT scores of TCGA-CESC.
**Supplementary File 4**: Primers used in the study.

#### Supplementary Table 1

Summary of the clinical metadata such as median age, five-year survival statistic, tumour stage, and grade associated with each of the samples.

| S.No. | Parameters | TCGA-CESC<br>N = 307 |
|---|---|---|
| **1.** | **Median Age**<br>**Inter Quartile Range** | 46.73<br>17.9 |
| **2.** | **Alive** | 247 |
| **3.** | **Dead** | 60 |
| **4.** | **Non-Available** | - |
| **5.** | **Metastasis** | 10 |
| **6.** | **Non-Metastasis** | 116 |
| **7.** | **Unknown metastasis** | 131 |
| **8.** | **Radiation**<br>**Yes**<br>**No** | <br>110<br>48 |
| **9.** | **Stage Classification** | |
| A. | Stage I | 163 |
| B. | Stage II | 70 |
| C. | Stage III | 46 |
| D. | Stage IV | 21 |
| E. | Non-Available | 7 |
| **10.** | **Tumour Grade** | |
| A. | GX (Grade cannot be assessed) | 24 |
| B. | G1 (Well differentiated) | 18 |
| C. | G2 (Moderately differentiated) | 136 |
| D. | G3 (Poorly or undifferentiated) | 20 |
| E. | Non-Available | 8 |

**Supplementary Table 2**

| Sample Name | Type of sample | Age /Sex | Clinical Symptoms |
|---|---|---|---|
| C1 | FT | 65/F | Third Degree Uterus prolapse, cystocele |
| C2 | FT | 45/F | Third degree prolapse, cystocele |
| C3 | FT | 55/F | Third Degree Uterus prolapse, cystocele |
| C4 | FT | 40/F | Third degree prolapse, cystocele |
| C5 | FT | 55/F | Third degree prolapse, cystocele |
| T1 | FT | 63/F | IIB, MD,CESC |
| T2 | FT | 50/F | IIB, MD,CESC |
| T3 | FT | 55/Y | IIB, MD,CESC |
| T4 | FT | 60/F | IIB, MD,CESC |
| T5 | FT | 42/F | IIB, PD,CESC |
| T6 | FT | 50/F | IIIB, MD,CESC |
| T7 | FT | 55/F | IIIB, MD,CESC |
| T8 | FT | 70/F | IIIB, MD,CESC |
| T9 | FT | 60/F | IIIB, MD,CESC |
| T10 | FT | 45/F | IIIB, MD,CESC |
| T11 | FFPE | 37/F | IIB, WD,CESC |
| T12 | FFPE | 65/F | IIB, MD,CESC |
| T13 | FFPE | 69/F | IIA, MD,CESC |
| T14 | FFPE | 45/F | IIA, MD,CESC |
| T15 | FFPE | 60/F | IIB,M D,CESC |
| T16 | FFPE | 40/F | IIIB,MD, CESC |
| T17 | FFPE | 43/F | IIIB, MD,CESC |
| T18 | FFPE | 46/F | IIIB, MD,CESC |
| T19 | FFPE | 65/F | IIIB, MD,CESC |
| T20 | FFPE | 62/F | IIIA, MD,CESC |
| T21 | FFPE | 65/F | IIIB,MD, CESC |

C: Non-cancerous cervical tissue; T: CESC tumour tissue; FT: Fresh frozen tissue; FFPE: Formalin fixed paraffin embedded tissue; F: female; WD: Well differentiated; MD: Moderately differentiated; PD: Poorly differentiated; CESC: Cervical epithelial squamous carcinoma.

Figure S1

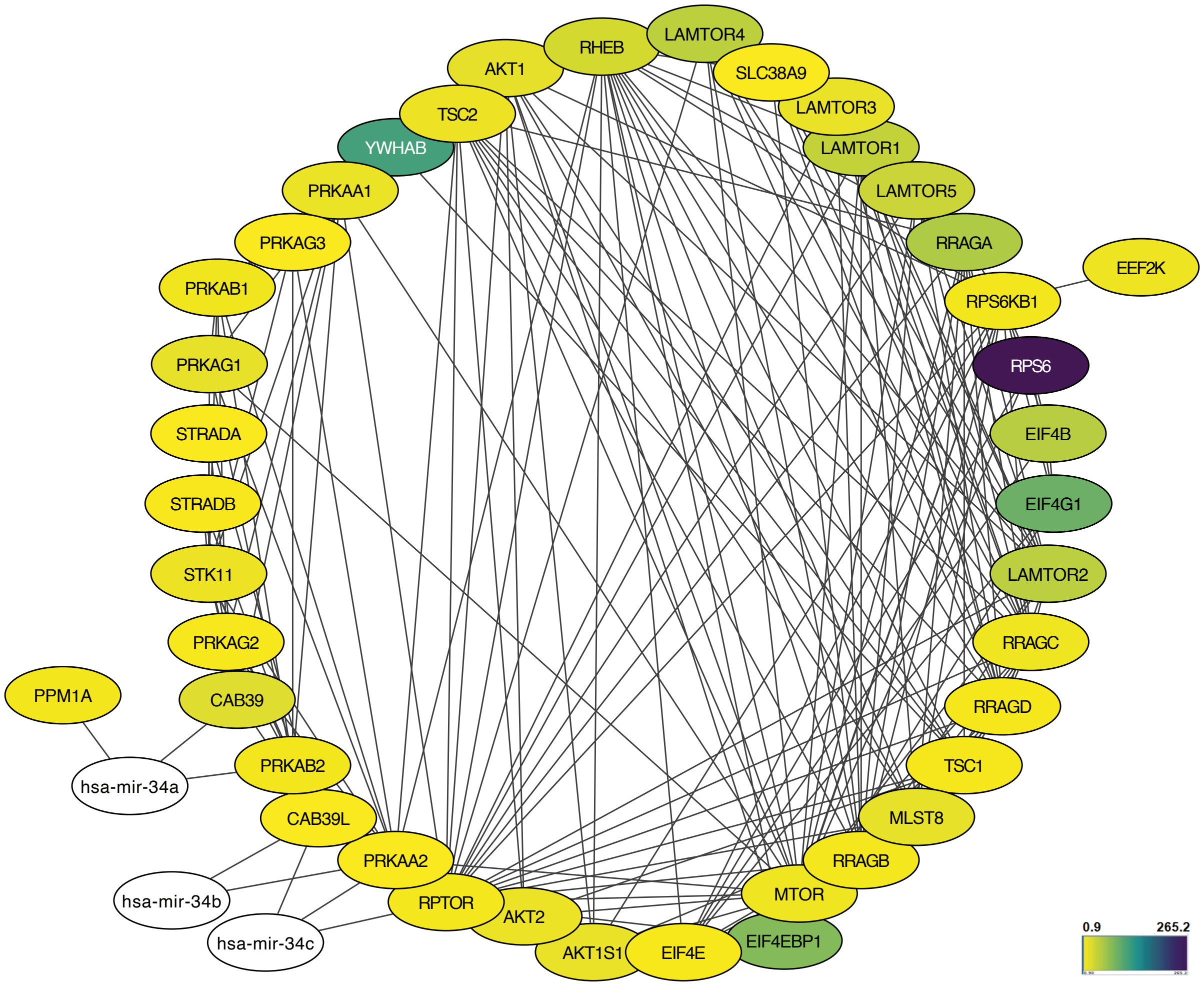

AKT1
RHEB
LAMTOR4
SLC38A9
TSC2
LAMTOR3
YWHAB
LAMTOR1
PRKAA1
LAMTOR5
PRKAG3
RRAGA
EEF2K
PRKAB1
RPS6KB1
RPS6
PRKAG1
STRADA
EIF4B
STRADB
EIF4G1
STK11
LAMTOR2
PRKAG2
RRAGC
PPM1A
CAB39
RRAGD
PRKAB2
TSC1
hsa-mir-34a
CAB39L
MLST8
PRKAA2
RRAGB
hsa-mir-34b
RPTOR
AKT2
MTOR
hsa-mir-34c
AKT1S1
EIF4E
EIF4EBP1
0.9
265.2

Figure S2

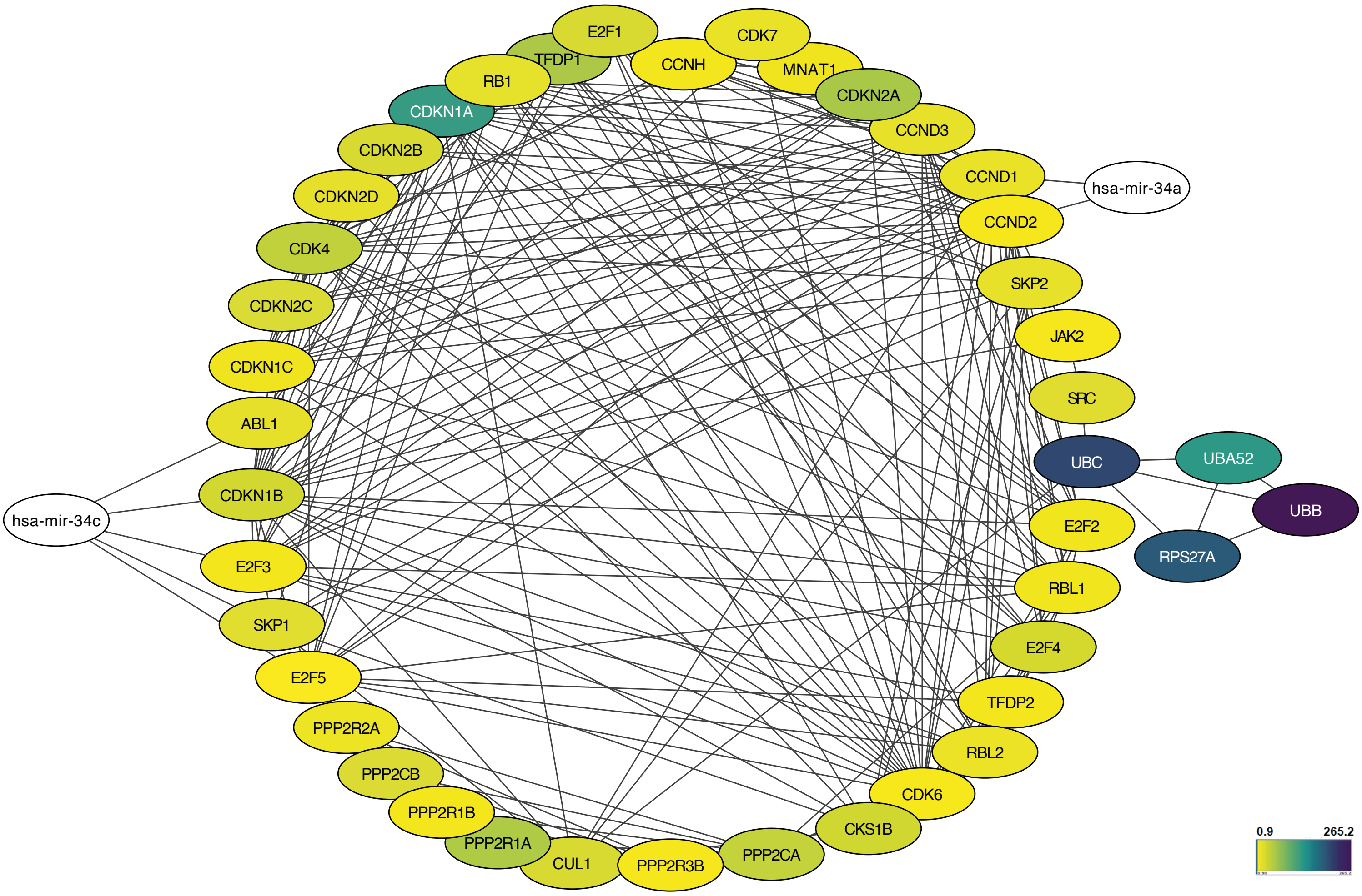
E2F1
TFDP1
CDK7
CCNH
MNAT1
RB1
CDKN2A
CDKN1A
CCND3
CDKN2B
CCND1
hsa-mir-34a
CDKN2D
CCND2
CDK4
SKP2
CDKN2C
JAK2
CDKN1C
SRC
ABL1
UBC
UBA52
UBB
CDKN1B
hsa-mir-34c
E2F2
RPS27A
E2F3
RBL1
SKP1
E2F4
E2F5
TFDP2
PPP2R2A
RBL2
PPP2CB
CDK6
PPP2R1B
CKS1B
PPP2R1A
CUL1
PPP2R3B
PPP2CA
0.9
265.2

Figure S3

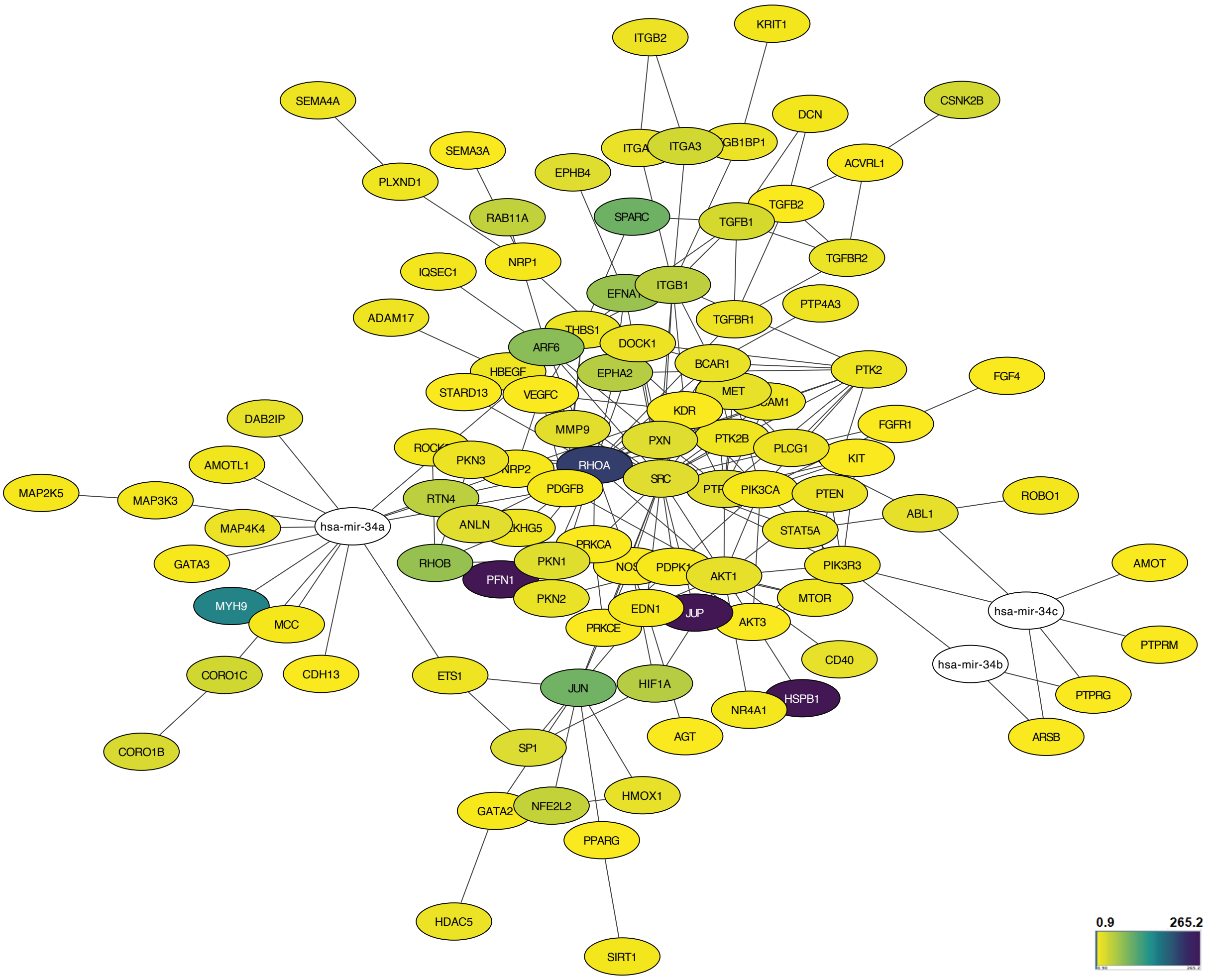

ITGB2
KRIT1
SEMA4A
CSNK2B
DCN
SEMA3A
ITGA3
ITGB1BP1
ACVRL1
PLXND1
EPHB4
TGFB2
RAB11A
SPARC
TGFB1
TGFBR2
IQSEC1
NRP1
ITGB1
PTP4A3
ADAM17
THBS1
TGFBR1
ARF6
DOCK1
HBEGF
EPHA2
BCAR1
PTK2
FGF4
STARD13
VEGFC
MET
KDR
DAB2IP
MMP9
FGFR1
PXN
PTK2B
PLCG1
AMOTL1
PKN3
KIT
RHOA
MAP2K5
MAP3K3
NRP2
SRC
RTN4
PDGFB
PIK3CA
PTEN
ROBO1
ABL1
MAP4K4
hsa-mir-34a
ANLN
STAT5A
PRKCA
AMOT
GATA3
RHOB
PKN1
PDPK1
PIK3R3
PFN1
AKT1
MYH9
PKN2
MTOR
hsa-mir-34c
EDN1
JUP
AKT3
MCC
PRKCE
PTPRM
CD40
hsa-mir-34b
CORO1C
CDH13
ETS1
JUN
HIF1A
HSPB1
PTPRG
NR4A1
AGT
ARSB
CORO1B
SP1
HMOX1
GATA2
NFE2L2
PPARG
HDAC5
SIRT1
0.9
265.2

Figure S4

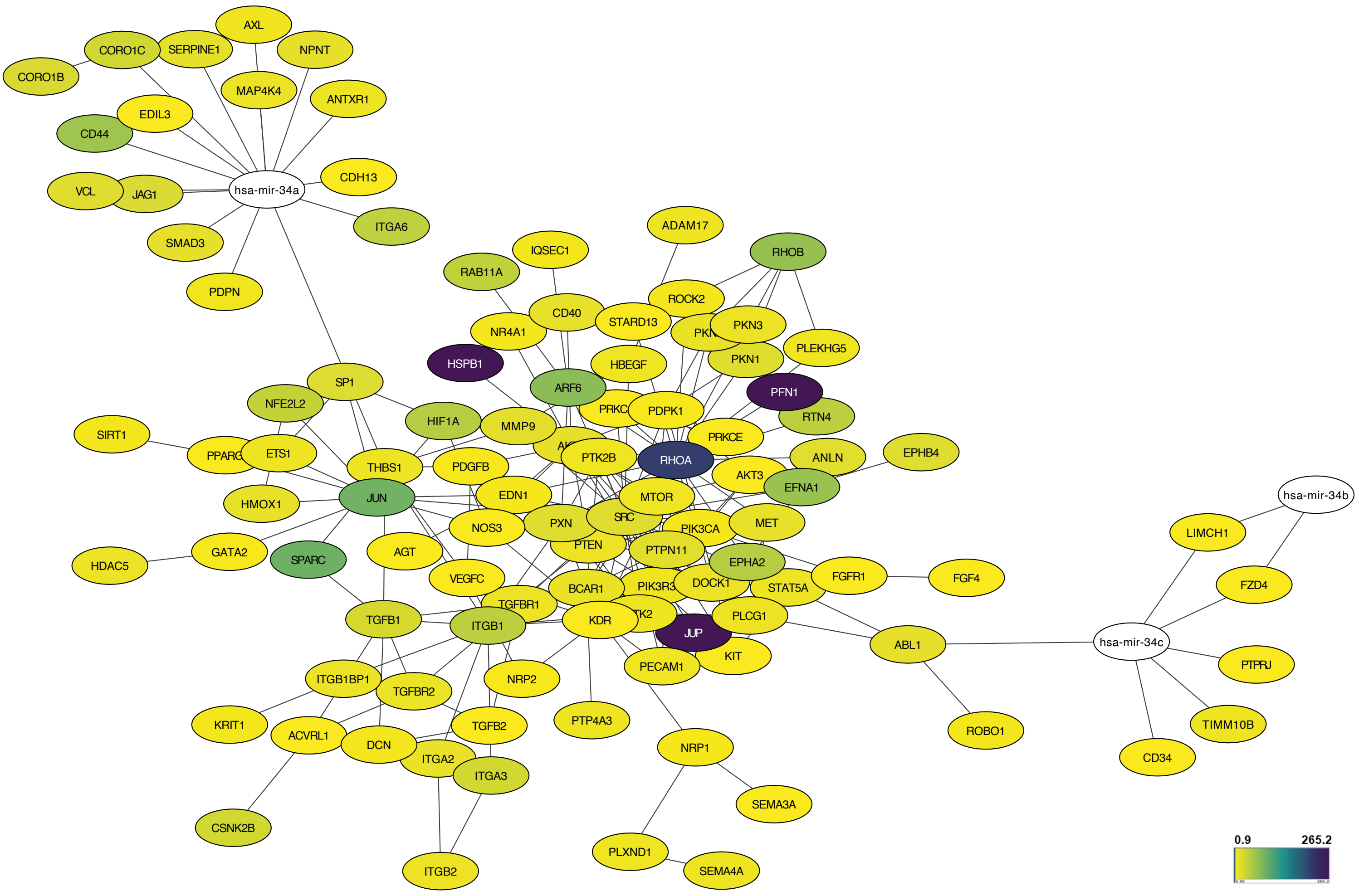

AXL
CORO1C
SERPINE1
NPNT
CORO1B
MAP4K4
ANTXR1
EDIL3
CD44
CDH13
VCL
JAG1
hsa-mir-34a
ITGA6
SMAD3
PDPN
ADAM17
IQSEC1
RHOB
RAB11A
ROCK2
CD40
STARD13
PKN3
NR4A1
PKN1
PLEKHG5
HSPB1
HBEGF
SP1
ARF6
PFN1
NFE2L2
PDPK1
RTN4
HIF1A
MMP9
SIRT1
PRKCE
ANLN
EPHB4
PTK2B
RHOA
ETS1
THBS1
PDGFB
AKT3
EFNA1
HMOX1
JUN
EDN1
MTOR
hsa-mir-34b
SRC
LIMCH1
NOS3
PXN
MET
PIK3CA
HDAC5
GATA2
SPARC
AGT
PTEN
PTPN11
EPHA2
FZD4
FGFR1
FGF4
VEGFC
BCAR1
DOCK1
STAT5A
TGFBR1
PLCG1
TGFB1
ITGB1
KDR
JUP
ABL1
hsa-mir-34c
KIT
PECAM1
PTPRJ
ITGB1BP1
NRP2
TGFBR2
KRIT1
ACVRL1
TGFB2
PTP4A3
ROBO1
TIMM10B
DCN
NRP1
ITGA2
ITGA3
CD34
SEMA3A
CSNK2B
PLXND1
SEMA4A
ITGB2
0.9
265.2

### Figure S5

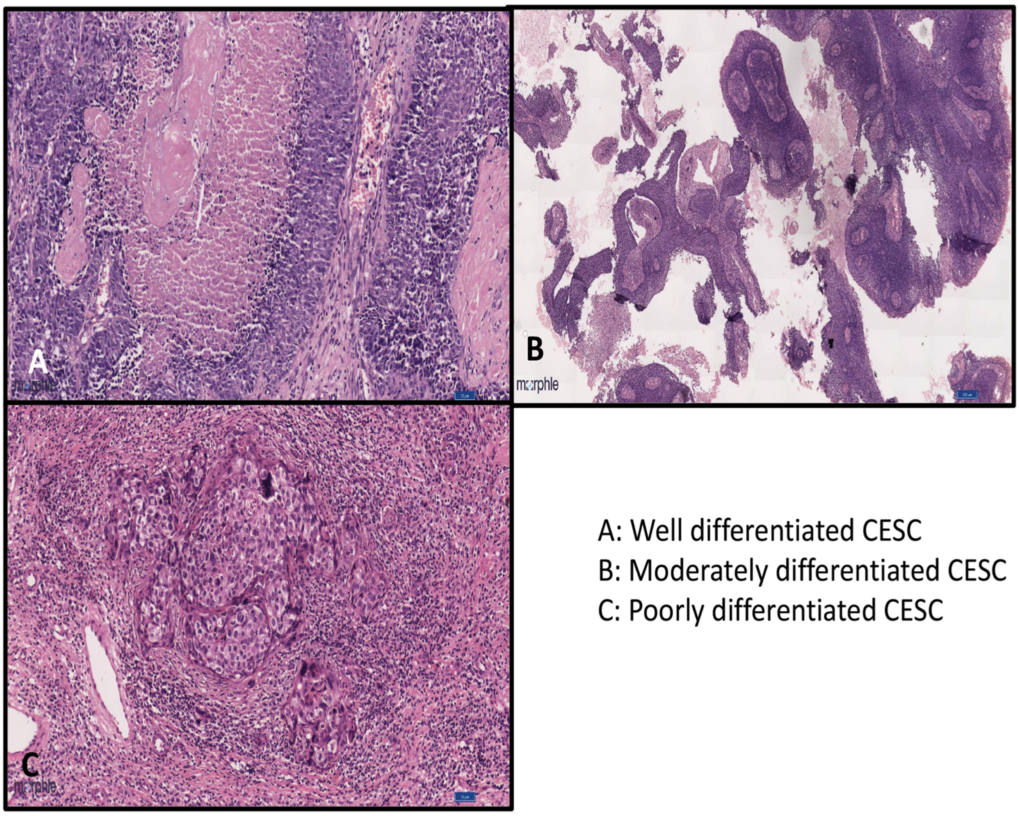

A: Well differentiated CESC
B: Moderately differentiated CESC
C: Poorly differentiated CESC

#### Supplementary File 1a: GO Output - miR-34a/b/c

| S.No | ID | Description | GeneRatio | BgRatio | p-value | p.adjust | q-value | geneID | Count |
|---|---|---|---|---|---|---|---|---|---|
| 1. | GO:0090287 | regulation of cellular response to growth factor stimulus | 19/353 | 196/11632 | 7.94E-06 | 0.02 | 0.02 | ENSG00000168743/ENSG00000166949/ENSG00000069869/ENSG00000171310/ENSG00000100614/ENSG00000137801/ENSG00000156535/ENSG00000105974/ENSG00000107485/ENSG00000136848/ENSG00000164056/ENSG00000174804/ENSG00000197879/ENSG00000097007/ENSG00000157613/ENSG00000102755/ENSG00000055163/ENSG00000187678/ENSG00000164442 | 19 |
| 2. | GO:0061041 | regulation of wound healing | 13/353 | 107/11632 | 2.05E-05 | 0.02 | 0.02 | ENSG00000166949/ENSG00000065675/ENSG00000065534/ENSG00000106366/ENSG00000100311/ENSG00000100345/ENSG00000137801/ENSG00000156535/ENSG00000178726/ENSG00000105974/ENSG00000162493/ENSG00000111276/ENSG00000174059 | 13 |
| 3. | GO:0030856 | regulation of epithelial cell differentiation | 13/353 | 108/11632 | 2.26E-05 | 0.02 | 0.02 | ENSG00000126821/ENSG00000110092/ENSG00000101384/ENSG00000106366/ENSG00000118263/ENSG00000156535/ENSG00000105974/ENSG00000137693/ENSG00000107485/ENSG00000065970/ENSG00000111276/ENSG00000080298/ENSG00000204301 | 13 |
| 4. | GO:0090288 | negative regulation of cellular response to growth factor stimulus | 13/353 | 111/11632 | 3.04E-05 | 0.02 | 0.02 | ENSG00000166949/ENSG00000069869/ENSG00000171310/ENSG00000100614/ENSG00000137801/ENSG00000156535/ENSG00000105974/ENSG00000107485/ENSG00000136848/ENSG00000164056/ENSG00000097007/ENSG00000157613/ENS | 13 |

| | | | | | | | | | |
|---|---|---|---|---|---|---|---|---|---|
| | | | | | | | | G00000187678 | |
| 5. | GO:0016311 | dephosphorylation | 27/353 | 382/11632 | 3.94E-05 | 0.02 | 0.02 | ENSG00000168743/ENSG00000058272/ENSG00000152104/ENSG00000126821/ENSG00000166949/ENSG00000074590/ENSG00000133789/ENSG00000108061/ENSG00000073711/ENSG00000100614/ENSG00000170525/ENSG00000132334/ENSG00000175215/ENSG00000172348/ENSG00000184007/ENSG00000144724/ENSG00000196950/ENSG00000173482/ENSG00000112245/ENSG00000112379/ENSG00000211456/ENSG00000149177/ENSG00000134324/ENSG00000184545/ENSG00000120875/ENSG00000112640/ENSG00000160014 | 27 |
| 6. | GO:0001667 | ameboidal-type cell migration | 24/353 | 319/11632 | 3.97E-05 | 0.02 | 0.02 | ENSG00000090776/ENSG00000071054/ENSG00000198909/ENSG00000171444/ENSG00000100311/ENSG00000100345/ENSG00000137801/ENSG00000049130/ENSG00000118257/ENSG00000140945/ENSG00000134954/ENSG00000166025/ENSG00000115310/ENSG00000131236/ENSG00000107485/ENSG00000136848/ENSG00000110880/ENSG00000117461/ENSG00000113273/ENSG00000144724/ENSG00000173482/ENSG00000097007/ENSG00000126016/ENSG00000179889 | 24 |
| 7. | GO:0040037 | negative regulation of fibroblast growth factor receptor signaling pathway | 5/353 | 14/11632 | 4.00E-05 | 0.02 | 0.02 | ENSG00000137801/ENSG00000107485/ENSG00000164056/ENSG00000157613/ENSG00000187678 | 5 |

| | | | | | | | | | |
|---|---|---|---|---|---|---|---|---|---|
| 8. | GO:0010631 | epithelial cell migration | 20/353 | 241/11632 | 4.52E-05 | 0.02 | 0.02 | ENSG00000071054/ENSG00000198909/ENSG00000171444/ENSG00000100311/ENSG00000100345/ENSG00000137801/ENSG00000118257/ENSG00000140945/ENSG00000134954/ENSG00000166025/ENSG00000115310/ENSG00000107485/ENSG00000136848/ENSG00000110880/ENSG00000117461/ENSG00000113273/ENSG00000144724/ENSG00000173482/ENSG00000097007/ENSG00000126016 | 20 |
| 9. | GO:0090132 | epithelium migration | 20/353 | 242/11632 | 4.79E-05 | 0.02 | 0.02 | ENSG00000071054/ENSG00000198909/ENSG00000171444/ENSG00000100311/ENSG00000100345/ENSG00000137801/ENSG00000118257/ENSG00000140945/ENSG00000134954/ENSG00000166025/ENSG00000115310/ENSG00000107485/ENSG00000136848/ENSG00000110880/ENSG00000117461/ENSG00000113273/ENSG00000144724/ENSG00000173482/ENSG00000097007/ENSG00000126016 | 20 |
| 10. | GO:0090130 | tissue migration | 20/353 | 245/11632 | 5.70E-05 | 0.02 | 0.02 | ENSG00000071054/ENSG00000198909/ENSG00000171444/ENSG00000100311/ENSG00000100345/ENSG00000137801/ENSG00000118257/ENSG00000140945/ENSG00000134954/ENSG00000166025/ENSG00000115310/ENSG00000107485/ENSG00000136848/ENSG00000110880/ENSG00000117461/ENSG00000113273/ENSG00000144724/ENSG00000173482/ENSG00000097007/ENSG00000126016 | 20 |
| 11. | GO:0007596 | blood coagulation | 19/353 | 230/11632 | 7.43E-05 | 0.02 | 0.02 | ENSG00000089902/ENSG00000088387/ENSG00000167601/ENSG00000065675/ENSG00000106366/ENS | 19 |

| | | | | | | | | | |
|---|---|---|---|---|---|---|---|---|---|
| | | | | | | | | G00000100311/ENSG00000100345/ENSG00000137801/ENSG00000178726/ENSG00000105974/ENSG00000035403/ENSG00000107485/ENSG00000162493/ENSG00000077044/ENSG00000131381/ENSG00000197249/ENSG00000137486/ENSG00000174059/ENSG00000138185 | |
| 12. | GO:0050817 | coagulation | 19/353 | 231/11632 | 7.88E-05 | 0.02 | 0.02 | ENSG00000089902/ENSG00000088387/ENSG00000167601/ENSG00000065675/ENSG00000106366/ENSG00000100311/ENSG00000100345/ENSG00000137801/ENSG00000178726/ENSG00000105974/ENSG00000035403/ENSG00000107485/ENSG00000162493/ENSG00000077044/ENSG00000131381/ENSG00000197249/ENSG00000137486/ENSG00000174059/ENSG00000138185 | 19 |
| 13. | GO:0007160 | cell-matrix adhesion | 16/353 | 175/11632 | 8.38E-05 | 0.02 | 0.02 | ENSG00000168743/ENSG00000166949/ENSG00000071054/ENSG00000101384/ENSG00000106366/ENSG00000137801/ENSG00000091409/ENSG00000140945/ENSG00000035403/ENSG00000026508/ENSG00000110880/ENSG00000064042/ENSG00000097007/ENSG00000149177/ENSG00000174059/ENSG00000132286 | 16 |
| 14. | GO:0007599 | hemostasis | 19/353 | 234/11632 | 9.36E-05 | 0.02 | 0.02 | ENSG00000089902/ENSG00000088387/ENSG00000167601/ENSG00000065675/ENSG00000106366/ENSG00000100311/ENSG00000100345/ENSG00000137801/ENSG00000178726/ENSG00000105974/ENSG00000035403/ENSG00000107485/ENSG00000162493/ENSG00000077044/ENSG00000131381/ENSG00000197249/ENSG00000137486/ENSG000 | 19 |

| | | | | | | | | | |
|---|---|---|---|---|---|---|---|---|---|
| | | | | | | | | 00174059/ENSG00000138185 | |
| 15. | GO:0042060 | wound healing | 27/353 | 402/11632 | 9.42E-05 | 0.02 | 0.02 | ENSG00000089902/ENSG00000088387/ENSG00000166949/ENSG00000167601/ENSG00000065675/ENSG00000065534/ENSG00000106366/ENSG00000100311/ENSG00000100345/ENSG00000137801/ENSG00000156535/ENSG00000178726/ENSG00000134954/ENSG00000105974/ENSG00000035403/ENSG00000137693/ENSG00000107485/ENSG00000162493/ENSG00000026508/ENSG00000077044/ENSG00000131381/ENSG00000111276/ENSG00000197249/ENSG00000137486/ENSG00000174059/ENSG00000204301/ENSG00000138185 | 27 |
| 16. | GO:0030335 | positive regulation of cell migration | 28/353 | 425/11632 | 9.73E-05 | 0.02 | 0.02 | ENSG00000131845/ENSG00000140443/ENSG00000166949/ENSG00000071054/ENSG00000198909/ENSG00000133789/ENSG00000065534/ENSG00000106366/ENSG00000100311/ENSG00000137801/ENSG00000049130/ENSG00000118257/ENSG00000091409/ENSG00000140945/ENSG00000134954/ENSG00000105974/ENSG00000166025/ENSG00000115310/ENSG00000107485/ENSG00000136848/ENSG00000162493/ENSG00000117461/ENSG00000197879/ENSG00000171105/ENSG00000112245/ENSG00000097007/ENSG00000128567/ENSG00000102755 | 28 |
| 17. | GO:0048193 | Golgi vesicle transport | 23/353 | 317/11632 | 1.00E-04 | 0.02 | 0.02 | ENSG00000197121/ENSG00000185963/ENSG00000198382/ENSG00000257923/ENSG00000197535/ENSG00000180447/ENSG00000131381/ENSG00000152700/ENSG0000013 | 23 |

| | | | | | | | | | |
|---|---|---|---|---|---|---|---|---|---|
| | | | | | | | | 4970/ENSG00000136152/ENSG00000168952/ENSG0000138078/ENSG00000220205/ENSG00000197249/ENSG00000107651/ENSG00000113719/ENSG00000101350/ENSG00000133103/ENSG00000113615/ENSG00000093183/ENSG00000108587/ENSG00000145817/ENSG00000148396 | |
| 18. | GO:0048514 | blood vessel morphogenesis | 30/353 | 473/11632 | 1.09E-04 | 0.02 | 0.02 | ENSG00000131845/ENSG00000112531/ENSG00000198909/ENSG00000101384/ENSG00000147649/ENSG00000064989/ENSG00000065534/ENSG00000106366/ENSG00000100345/ENSG00000137801/ENSG00000118257/ENSG00000140945/ENSG00000134954/ENSG00000105974/ENSG00000166025/ENSG00000120708/ENSG00000137693/ENSG00000136848/ENSG00000065970/ENSG00000117461/ENSG00000174804/ENSG00000181555/ENSG00000173482/ENSG00000097007/ENSG00000157613/ENSG00000174059/ENSG00000102755/ENSG00000204301/ENSG00000134817/ENSG00000126016 | 30 |
| 19. | GO:0031589 | cell-substrate adhesion | 21/353 | 279/11632 | 1.20E-04 | 0.02 | 0.02 | ENSG00000168743/ENSG00000166949/ENSG00000167601/ENSG00000071054/ENSG00000101384/ENSG00000106366/ENSG00000137801/ENSG00000091409/ENSG00000140945/ENSG00000164176/ENSG00000035403/ENSG00000169604/ENSG00000162493/ENSG00000026508/ENSG00000110880/ENSG00000064042/ENSG00000174804/ENSG00000097007/ENSG00000149177/ENSG00000174059/ENSG00000132286 | 21 |

| | | | | | | | | | |
|---|---|---|---|---|---|---|---|---|---|
| 20. | GO:0043407 | negative regulation of MAP kinase activity | 9/353 | 64/11632 | 1.24E-04 | 0.02 | 0.02 | ENSG00000117152/ENSG00000105974/ENSG00000136848/ENSG00000164056/ENSG00000149177/ENSG00000184545/ENSG00000120875/ENSG00000187678/ENSG00000138835 | 9 |
| 21. | GO:0006470 | protein dephosphorylation | 20/353 | 260/11632 | 1.29E-04 | 0.02 | 0.02 | ENSG00000058272/ENSG00000152104/ENSG00000074590/ENSG00000133789/ENSG00000108061/ENSG00000073711/ENSG00000100614/ENSG00000132334/ENSG00000175215/ENSG00000172348/ENSG00000184007/ENSG00000144724/ENSG00000196950/ENSG00000173482/ENSG00000112245/ENSG00000149177/ENSG00000184545/ENSG00000120875/ENSG00000112640/ENSG00000160014 | 20 |
| 22. | GO:0050678 | regulation of epithelial cell proliferation | 20/353 | 260/11632 | 1.29E-04 | 0.02 | 0.02 | ENSG00000131845/ENSG00000166949/ENSG00000110092/ENSG00000171444/ENSG00000100311/ENSG00000137801/ENSG00000118257/ENSG00000140945/ENSG00000156535/ENSG00000105974/ENSG00000115310/ENSG00000137693/ENSG00000107485/ENSG00000136848/ENSG00000173482/ENSG00000147862/ENSG00000111276/ENSG00000102755/ENSG00000134817/ENSG00000141564 | 20 |
| 23. | GO:0040017 | positive regulation of locomotion | 29/353 | 456/11632 | 1.35E-04 | 0.02 | 0.02 | ENSG00000131845/ENSG00000140443/ENSG00000166949/ENSG00000071054/ENSG00000198909/ENSG00000133789/ENSG00000065534/ENSG00000106366/ENSG00000100311/ENSG00000137801/ENSG00000049130/ENSG00000118257/ENSG00000091409/ENSG00000140945/ENSG00000134954/ENSG0000010 | 29 |

| | | | | | | | | | |
|---|---|---|---|---|---|---|---|---|---|
| | | | | | | | | 5974/ENSG00000166025/ENSG00000115310/ENSG00000107485/ENSG00000136848/ENSG00000162493/ENSG00000117461/ENSG00000083937/ENSG00000197879/ENSG00000171105/ENSG00000112245/ENSG00000097007/ENSG00000128567/ENSG00000102755 | |
| 24. | GO:2000147 | positive regulation of cell motility | 28/353 | 437/11632 | 1.56E-04 | 0.02 | 0.02 | ENSG00000131845/ENSG00000140443/ENSG00000166949/ENSG00000071054/ENSG00000198909/ENSG00000133789/ENSG00000065534/ENSG00000106366/ENSG00000100311/ENSG00000137801/ENSG00000049130/ENSG00000118257/ENSG00000091409/ENSG00000140945/ENSG00000134954/ENSG00000105974/ENSG00000166025/ENSG00000115310/ENSG00000107485/ENSG00000136848/ENSG00000162493/ENSG00000117461/ENSG00000197879/ENSG00000171105/ENSG00000112245/ENSG00000097007/ENSG00000128567/ENSG00000102755 | 28 |
| 25. | GO:1903034 | regulation of response to wounding | 13/353 | 130/11632 | 1.58E-04 | 0.02 | 0.02 | ENSG00000166949/ENSG00000065675/ENSG00000065534/ENSG00000106366/ENSG00000100311/ENSG00000100345/ENSG00000137801/ENSG00000156535/ENSG00000178726/ENSG00000105974/ENSG00000162493/ENSG00000111276/ENSG00000174059 | 13 |
| 26. | GO:0071900 | regulation of protein serine/threonine kinase activity | 27/353 | 418/11632 | 1.79E-04 | 0.03 | 0.02 | ENSG00000140443/ENSG00000198909/ENSG00000118971/ENSG00000198382/ENSG00000110092/ENSG00000100311/ENSG00000137801/ENSG00000049130/ENSG00000135932/ENSG00000117152/ENSG00000105974/ENSG00000136848/ENS | 27 |

| | | | | | | | | | |
|---|---|---|---|---|---|---|---|---|---|
| | | | | | | | | G00000164056/ENSG00000174804/ENSG00000171105/ENSG00000111276/ENSG00000097007/ENSG00000149177/ENSG00000113328/ENSG00000137486/ENSG00000184545/ENSG00000120875/ENSG00000102755/ENSG00000187678/ENSG00000160014/ENSG00000141564/ENSG00000138835 | |
| 27. | GO:0010632 | regulation of epithelial cell migration | 16/353 | 188/11632 | 1.94E-04 | 0.03 | 0.02 | ENSG00000071054/ENSG00000198909/ENSG00000171444/ENSG00000100311/ENSG00000137801/ENSG00000118257/ENSG00000134954/ENSG00000166025/ENSG00000115310/ENSG00000107485/ENSG00000136848/ENSG00000110880/ENSG00000113273/ENSG00000144724/ENSG00000173482/ENSG00000097007 | 16 |
| 28. | GO:0035850 | epithelial cell differentiation involved in kidney development | 6/353 | 29/11632 | 1.97E-04 | 0.03 | 0.02 | ENSG00000101384/ENSG00000100311/ENSG00000137693/ENSG00000107485/ENSG00000174059/ENSG00000128567 | 6 |
| 29. | GO:0061045 | negative regulation of wound healing | 8/353 | 54/11632 | 2.04E-04 | 0.03 | 0.02 | ENSG00000166949/ENSG00000106366/ENSG00000100311/ENSG00000137801/ENSG00000156535/ENSG00000178726/ENSG00000111276/ENSG00000174059 | 8 |
| 30. | GO:2000146 | negative regulation of cell motility | 17/353 | 210/11632 | 2.25E-04 | 0.03 | 0.02 | ENSG00000101384/ENSG00000171444/ENSG00000106366/ENSG00000137801/ENSG00000136141/ENSG00000035403/ENSG00000107485/ENSG00000136848/ENSG00000110880/ENSG00000064042/ENSG00000144724/ENSG00000276644/ENSG00000173482/ENSG00000149177/ENSG00000187098/ENSG00000044115/ENSG00000164442 | 17 |

| | | | | | | | | | |
|---|---|---|---|---|---|---|---|---|---|
| 31. | GO:0051272 | positive regulation of cellular component movement | 28/353 | 447/11632 | 2.27E-04 | 0.03 | 0.02 | ENSG00000131845/ENSG00000140443/ENSG00000166949/ENSG00000071054/ENSG00000198909/ENSG00000133789/ENSG00000065534/ENSG00000106366/ENSG00000100311/ENSG00000137801/ENSG00000049130/ENSG00000118257/ENSG00000091409/ENSG00000140945/ENSG00000134954/ENSG00000105974/ENSG00000166025/ENSG00000115310/ENSG00000107485/ENSG00000136848/ENSG00000162493/ENSG00000117461/ENSG00000197879/ENSG00000171105/ENSG00000112245/ENSG00000097007/ENSG00000128567/ENSG00000102755 | 28 |
| 32. | GO:0034332 | adherens junction organization | 12/353 | 118/11632 | 2.40E-04 | 0.03 | 0.02 | ENSG00000166949/ENSG00000071054/ENSG00000110400/ENSG00000137801/ENSG00000140945/ENSG00000035403/ENSG00000110880/ENSG00000064042/ENSG00000182985/ENSG00000097007/ENSG00000149177/ENSG00000044115 | 12 |
| 33. | GO:0001525 | angiogenesis | 26/353 | 403/11632 | 2.41E-04 | 0.03 | 0.02 | ENSG00000131845/ENSG00000198909/ENSG00000101384/ENSG00000147649/ENSG00000064989/ENSG00000106366/ENSG00000100345/ENSG00000137801/ENSG00000118257/ENSG00000140945/ENSG00000134954/ENSG00000105974/ENSG00000166025/ENSG00000120708/ENSG00000136848/ENSG00000065970/ENSG00000117461/ENSG00000181555/ENSG00000173482/ENSG00000097007/ENSG00000157613/ENSG00000174059/ENSG00000102755/ENSG00000204301/ENSG00000134817/ENSG00000126016 | 26 |

| | | | | | | | | | |
|---|---|---|---|---|---|---|---|---|---|
| 34. | GO:0050878 | regulation of body fluid levels | 23/353 | 339/11632 | 2.69E-04 | 0.03 | 0.03 | ENSG00000089902/ENSG00000088387/ENSG00000167601/ENSG00000065675/ENSG00000110092/ENSG00000106366/ENSG00000100311/ENSG00000100345/ENSG00000137801/ENSG00000178726/ENSG00000105974/ENSG00000035403/ENSG00000107485/ENSG00000162493/ENSG00000077044/ENSG00000131381/ENSG00000123191/ENSG00000197249/ENSG00000137486/ENSG00000174059/ENSG00000134817/ENSG00000057663/ENSG00000138185 | 23 |
| 35. | GO:0010810 | regulation of cell-substrate adhesion | 15/353 | 175/11632 | 2.83E-04 | 0.03 | 0.03 | ENSG00000168743/ENSG00000166949/ENSG00000071054/ENSG00000101384/ENSG00000106366/ENSG00000137801/ENSG00000091409/ENSG00000140945/ENSG00000164176/ENSG00000162493/ENSG00000110880/ENSG00000064042/ENSG00000174804/ENSG00000097007/ENSG00000149177 | 15 |
| 36. | GO:0043409 | negative regulation of MAPK cascade | 13/353 | 138/11632 | 2.86E-04 | 0.03 | 0.03 | ENSG00000140443/ENSG00000198053/ENSG00000117152/ENSG00000105974/ENSG00000136848/ENSG00000164056/ENSG00000097007/ENSG00000149177/ENSG00000137486/ENSG00000184545/ENSG00000120875/ENSG00000187678/ENSG00000138835 | 13 |
| 37. | GO:0071901 | negative regulation of protein serine/threonine kinase activity | 12/353 | 121/11632 | 3.03E-04 | 0.03 | 0.03 | ENSG00000117152/ENSG00000105974/ENSG00000136848/ENSG00000164056/ENSG00000111276/ENSG00000097007/ENSG00000149177/ENSG00000184545/ENSG00000120875/ENSG00000187678/ENSG00000141564/ENSG00000138835 | 12 |

| | | | | | | | | | |
|---|---|---|---|---|---|---|---|---|---|
| 38. | GO:0051271 | negative regulation of cellular component movement | 18/353 | 238/11632 | 3.43E-04 | 0.03 | 0.03 | ENSG00000101384/ENSG00000171444/ENSG00000106366/ENSG00000137801/ENSG00000136141/ENSG00000035403/ENSG00000107485/ENSG00000136848/ENSG00000110880/ENSG00000064042/ENSG00000144724/ENSG00000276644/ENSG00000173482/ENSG00000111276/ENSG00000149177/ENSG00000187098/ENSG00000044115/ENSG00000164442 | 18 |
| 39. | GO:0030857 | negative regulation of epithelial cell differentiation | 6/353 | 32/11632 | 3.48E-04 | 0.03 | 0.03 | ENSG00000110092/ENSG00000101384/ENSG00000105974/ENSG00000137693/ENSG00000065970/ENSG00000204301 | 6 |
| 40. | GO:0050673 | epithelial cell proliferation | 21/353 | 303/11632 | 3.72E-04 | 0.03 | 0.03 | ENSG00000131845/ENSG00000166949/ENSG00000110092/ENSG00000171444/ENSG00000100311/ENSG00000137801/ENSG00000118257/ENSG00000140945/ENSG00000156535/ENSG00000105974/ENSG00000115310/ENSG00000137693/ENSG00000107485/ENSG00000136848/ENSG00000173482/ENSG00000147862/ENSG00000111276/ENSG00000174059/ENSG00000102755/ENSG00000134817/ENSG00000141564 | 21 |
| 41. | GO:0030193 | regulation of blood coagulation | 8/353 | 59/11632 | 3.81E-04 | 0.03 | 0.03 | ENSG00000065675/ENSG00000106366/ENSG00000100311/ENSG00000137801/ENSG00000178726/ENSG00000105974/ENSG00000162493/ENSG00000174059 | 8 |
| 42. | GO:1900046 | regulation of hemostasis | 8/353 | 60/11632 | 4.29E-04 | 0.04 | 0.03 | ENSG00000065675/ENSG00000106366/ENSG00000100311/ENSG00000137801/ENSG00000178726/ENSG00000105974/ENSG00000162493/ENSG00000174059 | 8 |
| 43. | GO:0040036 | regulation of fibroblast | 5/353 | 22/11632 | 4.30E-04 | 0.04 | 0.03 | ENSG00000137801/ENSG00000107485/ENSG00000164056/ENSG000 | 5 |

| | | | | | | | | | |
|---|---|---|---|---|---|---|---|---|---|
| | | growth factor receptor signaling pathway | | | | | | 00157613/ENSG00000187678 | |
| 44. | GO:0050818 | regulation of coagulation | 8/353 | 61/11632 | 4.80E-04 | 0.04 | 0.04 | ENSG00000065675/ENSG00000106366/ENSG00000100311/ENSG00000137801/ENSG00000178726/ENSG00000105974/ENSG00000162493/ENSG00000174059 | 8 |
| 45. | GO:0035335 | peptidyl-tyrosine dephosphorylation | 9/353 | 77/11632 | 5.16E-04 | 0.04 | 0.04 | ENSG00000152104/ENSG00000132334/ENSG00000184007/ENSG00000144724/ENSG00000173482/ENSG00000112245/ENSG00000149177/ENSG00000184545/ENSG00000120875 | 9 |
| 46. | GO:0040013 | negative regulation of locomotion | 18/353 | 247/11632 | 5.36E-04 | 0.04 | 0.04 | ENSG00000101384/ENSG00000171444/ENSG00000106366/ENSG00000137801/ENSG00000136141/ENSG00000035403/ENSG00000107485/ENSG00000136848/ENSG00000110880/ENSG00000113369/ENSG00000064042/ENSG00000144724/ENSG00000276644/ENSG00000173482/ENSG00000149177/ENSG00000187098/ENSG00000044115/ENSG00000164442 | 18 |
| 47. | GO:0000188 | inactivation of MAPK activity | 5/353 | 23/11632 | 5.36E-04 | 0.04 | 0.04 | ENSG00000117152/ENSG00000105974/ENSG00000184545/ENSG00000120875/ENSG00000138835 | 5 |
| 48. | GO:1902532 | negative regulation of intracellular signal transduction | 26/353 | 425/11632 | 5.44E-04 | 0.04 | 0.04 | ENSG00000140443/ENSG00000198053/ENSG00000100614/ENSG00000137801/ENSG00000143322/ENSG00000117152/ENSG00000105974/ENSG00000136848/ENSG00000026508/ENSG00000113369/ENSG00000144791/ENSG00000164056/ENSG00000162409/ENSG00000172572/ENSG00000159459/ENSG00000097007/ENSG00000149177/ENSG000 | 26 |

| | | | | | | | | | |
|---|---|---|---|---|---|---|---|---|---|
| | | | | | | | | 00157613/ENSG00000137486/ENSG00000140577/ENSG00000184545/ENSG00000120875/ENSG00000187678/ENSG00000134817/ENSG00000160014/ENSG00000138835 | |
| 49. | GO:0043951 | negative regulation of cAMP-mediated signaling | 4/353 | 14/11632 | 6.56E-04 | 0.05 | 0.04 | ENSG00000113369/ENSG00000172572/ENSG00000140577/ENSG00000134817 | 4 |
| 50. | GO:0001570 | vasculogenesis | 8/353 | 64/11632 | 6.67E-04 | 0.05 | 0.04 | ENSG00000112531/ENSG00000105974/ENSG00000137693/ENSG00000174804/ENSG00000181555/ENSG00000174059/ENSG00000134817/ENSG00000126016 | 8 |
| 51. | GO:0061005 | cell differentiation involved in kidney development | 6/353 | 36/11632 | 6.75E-04 | 0.05 | 0.04 | ENSG00000101384/ENSG00000100311/ENSG00000137693/ENSG00000107485/ENSG00000174059/ENSG00000128567 | 6 |
| 52. | GO:1901888 | regulation of cell junction assembly | 9/353 | 80/11632 | 6.84E-04 | 0.05 | 0.04 | ENSG00000166949/ENSG00000071054/ENSG00000137801/ENSG00000105974/ENSG00000110880/ENSG00000064042/ENSG00000197879/ENSG00000097007/ENSG00000149177 | 9 |
| 53. | GO:0009611 | response to wounding | 28/353 | 481/11632 | 7.34E-04 | 0.05 | 0.05 | ENSG00000089902/ENSG00000088387/ENSG00000166949/ENSG00000167601/ENSG00000065675/ENSG00000065534/ENSG00000106366/ENSG00000100311/ENSG00000100345/ENSG00000137801/ENSG00000156535/ENSG00000178726/ENSG00000134954/ENSG00000105974/ENSG00000035403/ENSG00000137693/ENSG00000107485/ENSG00000162493/ENSG00000026508/ENSG00000077044/ENSG00000131381/ENSG00000111276/ENSG000001972 | 28 |

| | | | | | | | | | |
|---|---|---|---|---|---|---|---|---|---|
| | | | | | | | | 49/ENSG00000137486/ENSG00000174059/ENSG00000204301/ENSG00000138185/ENSG00000044115 | |
| 54. | GO:1903510 | mucopolysaccharide metabolic process | 9/353 | 81/11632 | 7.50E-04 | 0.05 | 0.05 | ENSG00000122863/ENSG00000116704/ENSG00000171310/ENSG00000100311/ENSG00000070614/ENSG00000026508/ENSG00000113273/ENSG00000135677/ENSG00000170266 | 9 |
| 55. | GO:0006888 | ER to Golgi vesicle-mediated transport | 14/353 | 173/11632 | 7.97E-04 | 0.05 | 0.05 | ENSG00000197121/ENSG00000180447/ENSG00000152700/ENSG00000134970/ENSG00000136152/ENSG00000197249/ENSG00000107651/ENSG00000113719/ENSG00000133103/ENSG00000113615/ENSG00000093183/ENSG00000108587/ENSG00000145817/ENSG00000148396 | 14 |
| 56. | GO:1903035 | negative regulation of response to wounding | 8/353 | 66/11632 | 8.22E-04 | 0.05 | 0.05 | ENSG00000166949/ENSG00000106366/ENSG00000100311/ENSG00000137801/ENSG00000156535/ENSG00000178726/ENSG00000111276/ENSG00000174059 | 8 |
| 57. | GO:0001952 | regulation of cell-matrix adhesion | 10/353 | 99/11632 | 8.26E-04 | 0.05 | 0.05 | ENSG00000166949/ENSG00000071054/ENSG00000101384/ENSG00000106366/ENSG00000137801/ENSG00000140945/ENSG00000110880/ENSG00000064042/ENSG00000097007/ENSG00000149177 | 10 |

#### Supplementary File 1b : Reactome Output miR-34a/b/c

| S.No. | ID | Description | Gene Ratio | BgRatio | p-value | p.adjust | q-value | geneID | Count |
|---|---|---|---|---|---|---|---|---|---|
| 1. | R-HSA-380972 | Energy dependent regulation of mTOR by LKB1-AMPK | 6/229 | 29/10654 | 2.90E-05 | 0.01 | 0.01 | PRKAB2/PPM1A/CAB39/CAB39L/PRKAA2/RPTOR | 6 |
| 2. | R-HSA-69231 | Cyclin D associated events in G1 | 7/229 | 44/10654 | 3.76E-05 | 0.01 | 0.01 | CCND2/CCND1/SKP1/CDKN1B/ABL1/E2F5/E2F3 | 7 |
| 3. | R-HSA-69236 | G1 Phase | 7/229 | 44/10654 | 3.76E-05 | 0.01 | 0.01 | CCND2/CCND1/SKP1/CDKN1B/ABL1/E2F5/E2F3 | 7 |
| 4. | R-HSA-1445148 | Translocation of SLC2A4 (GLUT4) to the plasma membrane | 8/229 | 72/10654 | 1.48E-04 | 0.02 | 0.02 | PRKAB2/MYO5A/MYH9/PRKAA2/MYO1C/VAMP2/KIF3B/CALM3 | 8 |
| 5. | R-HSA-76002 | Platelet activation, signaling and aggregation | 16/229 | 262/10654 | 1.67E-04 | 0.02 | 0.02 | ENDOD1/PRKCQ/GNG12/SERPINE1/PDGFB/THBS1/CD109/VCL/CAP1/PDPN/DGKD/PIK3R3/SERPINA1/ARRB1/BRPF3/CALM3 | 16 |
| 6. | R-HSA-948021 | Transport to the Golgi and subsequent modification | 13/229 | 185/10654 | 1.76E-04 | 0.02 | 0.02 | FUT8/MAN2A2/SAR1B/TMED7/COG3/MGAT4A/SERPINA1/SEC23IP/COG6/SEC24A/SEC22C/GOSR1/SEC16A | 13 |
| 7. | R-HSA-165159 | mTOR signalling | 6/229 | 40/10654 | 1.92E-04 | 0.02 | 0.02 | PRKAB2/PPM1A/CAB39/CAB39L/PRKAA2/RPTOR | 6 |
| 8. | R-HSA-114608 | Platelet degranulation | 10/229 | 129/10654 | 4.56E-04 | 0.04 | 0.04 | ENDOD1/SERPINE1/PDGFB/THBS1/CD109/VCL/CAP1/SERPINA1/BRPF3/CALM3 | 10 |
| 9. | R-HSA-9006934 | Signaling by Receptor Tyrosine Kinases | 22/229 | 473/10654 | 5.39E-04 | 0.04 | 0.04 | IGF1R/AXL/NEDD4/COL5A2/PDGFB/THBS1/KITLG/NRP2/RBFOX2/CAV1/YAP1/PIK3R3/SPRY1/INSR/PTPRJ/FGFRL1/DUSP4/FLT1/CYFIP2/PPP2R5D/CALM3/CTNNA1 | 22 |
| 10. | R-HSA-76005 | Response to elevated platelet cytosolic Ca2+ | 10/229 | 134/10654 | 6.16E-04 | 0.04 | 0.04 | ENDOD1/SERPINE1/PDGFB/THBS1/CD109/VCL/CAP1/SERPINA1/BRPF3/CALM3 | 10 |

#### Supplementary File 2

| EMT_gene | condition | corr_vs_mir34b | pvalue |
|---|---|---|---|
| ACVR1 | G1 | 0.206015038 | 0.383533 |
| ACVR1 | G2 | -0.047968226 | 0.579198 |
| FSCN1 | GX | 0.703478261 | 1.26E-04 |
| ACVR1 | GX | -0.260869565 | 0.218238 |
| ACVR1 | Stage I | 0.077489006 | 0.325519 |
| ACVR1 | Stage II | 0.042589224 | 0.724348 |
| ACVR1 | Stage III | 0.187195462 | 0.212864 |
| ACVR1 | Stage IV | -0.162055336 | 0.471204 |
| ACVR1 | overall | 0.061483333 | 0.281294 |
| ADAM17 | G1 | 0.003007519 | 0.98996 |
| ADAM17 | G2 | 0.040857317 | 0.636732 |
| ADAM17 | G3 | 0.026951426 | 0.770133 |
| ADAM17 | GX | -0.016521739 | 0.938923 |
| ADAM17 | Stage I | 0.016540262 | 0.83401 |
| ADAM17 | Stage II | 0.185866749 | 0.120686 |
| ADAM17 | Stage III | 0.076420157 | 0.613712 |
| ADAM17 | Stage IV | -0.191417278 | 0.393465 |
| ADAM17 | overall | 0.031824003 | 0.57733 |
| AGER | G1 | 0.17593985 | 0.458099 |
| AGER | G2 | -0.071989313 | 0.404921 |
| AGER | G3 | 0.047002648 | 0.610205 |
| AGER | GX | 0.03826087 | 0.859118 |
| AGER | Stage I | 0.094788529 | 0.228753 |
| CAV1 | GX | 0.632173913 | 9.19E-04 |
| AGER | Stage III | -0.125023131 | 0.407746 |
| AGER | Stage IV | 0.357425184 | 0.102452 |
| AGER | overall | -0.014328865 | 0.801914 |
| AKT1 | G1 | -0.010526316 | 0.964869 |
| AKT1 | G2 | -0.106980905 | 0.215106 |
| AKT1 | G3 | 0.003649781 | 0.968442 |
| AKT1 | GX | -0.108695652 | 0.613156 |
| AKT1 | Stage I | 0.002178027 | 0.977987 |
| AKT1 | Stage II | -0.204830691 | 0.086617 |
| AKT1 | Stage III | -0.196940728 | 0.189571 |
| AKT1 | Stage IV | 0.197063806 | 0.379389 |
| AKT1 | overall | -0.083702357 | 0.142113 |
| ANPEP | G1 | -0.25112782 | 0.285522 |
| ANPEP | G2 | -0.106553916 | 0.216953 |
| ANPEP | G3 | -0.052062335 | 0.572259 |
| ANPEP | GX | -0.266086957 | 0.208836 |
| ANPEP | Stage I | -0.051305569 | 0.515425 |

| | | | |
|---|---|---|---|
| ANPEP | Stage II | -0.136822073 | 0.255212 |
| ANPEP | Stage III | -0.22037871 | 0.141099 |
| ANPEP | Stage IV | 0.119141728 | 0.597436 |
| ANPEP | overall | -0.090669622 | 0.11169 |
| ANXA1 | G1 | 0.108270677 | 0.649567 |
| ANXA1 | G2 | 0.150526579 | 0.080256 |
| ANXA1 | G3 | 0.158617876 | 0.083566 |
| ANXA1 | GX | 0.286086957 | 0.17534 |
| MYC | GX | 0.623478261 | 0.001134 |
| ANXA1 | Stage II | 0.09590959 | 0.426241 |
| ANXA1 | Stage III | 0.180040709 | 0.231188 |
| ANXA1 | Stage IV | -0.075098814 | 0.739774 |
| HRAS | GX | 0.62173913 | 0.001181 |
| ATM | G1 | -0.057142857 | 0.81088 |
| FOSL1 | GX | 0.603478261 | 0.001796 |
| ATM | G3 | -0.130638538 | 0.154969 |
| ATM | GX | -0.205217391 | 0.336061 |
| ATM | Stage I | -0.080739421 | 0.305574 |
| CD44 | GX | 0.596521739 | 0.002093 |
| ATM | Stage III | -0.083451552 | 0.581371 |
| ATM | Stage IV | -0.306606437 | 0.165179 |
| MMP14 | GX | 0.592173913 | 0.002299 |
| AURKA | G1 | -0.10075188 | 0.672556 |
| AURKA | G2 | -1.93E-04 | 0.998219 |
| AURKA | G3 | -0.06241785 | 0.498243 |
| AURKA | GX | 0.340869565 | 0.103099 |
| AURKA | Stage I | -0.049393561 | 0.531221 |
| AURKA | Stage II | 0.122485936 | 0.308867 |
| AURKA | Stage III | -0.105285883 | 0.486197 |
| AURKA | Stage IV | -0.326933936 | 0.137515 |
| AURKA | overall | -0.019817018 | 0.728611 |
| AXIN1 | G1 | 0.060150376 | 0.801113 |
| AXIN1 | G2 | -0.082737975 | 0.338255 |
| AXIN1 | G3 | -0.149529887 | 0.103091 |
| AXIN1 | GX | -0.251304348 | 0.236199 |
| AXIN1 | Stage I | -0.082063972 | 0.297682 |
| AXIN1 | Stage II | -0.20645713 | 0.084092 |
| AXIN1 | Stage III | -0.098007772 | 0.516991 |
| AXIN1 | Stage IV | -0.023150762 | 0.918549 |
| THBD | GX | 0.588695652 | 0.002476 |
| AXIN2 | G1 | -0.05112782 | 0.830495 |
| AXIN2 | G2 | -0.013976122 | 0.871706 |
| AXIN2 | G3 | -0.076298129 | 0.407521 |
| AXIN2 | GX | -0.213913043 | 0.315527 |

| | | | |
|---|---|---|---|
| AXIN2 | Stage I | 0.014173804 | 0.857486 |
| HBEGF | GX | 0.579130435 | 0.003023 |
| AXIN2 | Stage III | -0.160056746 | 0.287985 |
| AXIN2 | Stage IV | 0.025409373 | 0.910632 |
| AXIN2 | overall | -0.075886835 | 0.18336 |
| AXL | G1 | 0.045112782 | 0.850206 |
| AXL | G2 | 0.045334733 | 0.600224 |
| PTPRZ1 | GX | 0.579130435 | 0.003023 |
| AXL | GX | 0.404347826 | 0.050024 |
| ID1 | GX | 0.552173913 | 0.005147 |
| YBX1 | GX | 0.549565217 | 0.005407 |
| AXL | Stage III | -0.029914267 | 0.843555 |
| AXL | Stage IV | 0.055900621 | 0.804842 |
| JAG2 | GX | 0.535652174 | 0.006983 |
| BCL2 | G1 | 0.078195489 | 0.743149 |
| BCL2 | G2 | -0.089610343 | 0.299517 |
| BCL2 | G3 | -0.013994878 | 0.879416 |
| SNAI2 | GX | 0.528695652 | 0.007905 |
| BCL2 | Stage I | -0.069347728 | 0.379066 |
| BCL2 | Stage II | -0.117941968 | 0.327293 |
| BCL2 | Stage III | -0.014124468 | 0.925772 |
| BCL2 | Stage IV | 0.106719368 | 0.636432 |
| BCL2 | overall | -0.061409102 | 0.281876 |
| BCL2L1 | G1 | -0.138345865 | 0.560787 |
| BCL2L1 | G2 | -0.003704543 | 0.965858 |
| BCL2L1 | G3 | 0.032038894 | 0.728303 |
| BCL2L1 | GX | 0.052173913 | 0.808688 |
| BCL2L1 | Stage I | 0.053516849 | 0.497464 |
| BCL2L1 | Stage II | -0.093360944 | 0.438692 |
| BCL2L1 | Stage III | 0.200641462 | 0.18122 |
| BCL2L1 | Stage IV | -0.269339356 | 0.225463 |
| BCL2L1 | overall | 0.020361237 | 0.721463 |
| BIRC2 | G1 | 0.252631579 | 0.28255 |
| BIRC2 | G2 | -0.002399723 | 0.97788 |
| BIRC2 | G3 | 0.17233494 | 0.059811 |
| BIRC2 | GX | 0.010434783 | 0.961404 |
| FGFR2 | G1 | 0.52481203 | 0.017509 |
| BIRC2 | Stage II | 0.179226855 | 0.134777 |
| BIRC2 | Stage III | -0.11046691 | 0.464874 |
| BIRC2 | Stage IV | -0.186900056 | 0.404933 |
| BIRC2 | overall | 0.091286648 | 0.109263 |
| BMI1 | G1 | 0.421052632 | 0.064488 |
| BMI1 | G2 | -0.042217001 | 0.625547 |
| BMI1 | G3 | -0.090709728 | 0.324473 |

| | | | |
|---|---|---|---|
| EGFR | GX | 0.52173913 | 0.008926 |
| BMI1 | Stage I | -0.101544293 | 0.197124 |
| BMI1 | Stage II | -0.042924572 | 0.722265 |
| BMI1 | Stage III | 0.072966139 | 0.629874 |
| BMI1 | Stage IV | -0.015245624 | 0.946313 |
| BMI1 | overall | -0.068933195 | 0.226947 |
| BMP2 | G1 | 0.17593985 | 0.458099 |
| BMP2 | G2 | -0.044857651 | 0.604069 |
| BMP2 | G3 | 0.131812302 | 0.151262 |
| BMP2 | GX | 0.188695652 | 0.377216 |
| BMP2 | Stage I | 0.069519532 | 0.377883 |
| PKP3 | GX | 0.520869565 | 0.009061 |
| BMP2 | Stage III | -0.156109296 | 0.300195 |
| BMP2 | Stage IV | 0.043478261 | 0.84765 |
| BMP2 | overall | 0.063942284 | 0.262461 |
| BMP4 | G1 | 0.311278195 | 0.181576 |
| BMP4 | G2 | -0.036136589 | 0.67619 |
| BMP4 | G3 | -0.037859096 | 0.681416 |
| BMP4 | GX | -0.199130435 | 0.350899 |
| BMP4 | Stage I | -0.018053243 | 0.819076 |
| BMP4 | Stage II | -0.146480101 | 0.222866 |
| BMP4 | Stage III | -0.089126011 | 0.555848 |
| BMP4 | Stage IV | 0.17108978 | 0.44649 |
| BMP4 | overall | -0.0429929 | 0.451431 |
| BMP7 | G1 | 0.272180451 | 0.245669 |
| EDN1 | G1 | 0.509774436 | 0.021669 |
| EIF5A2 | GX | 0.508695652 | 0.011136 |
| BMP7 | GX | 0.126956522 | 0.554416 |
| MMP13 | GX | 0.503478261 | 0.012137 |
| BMP7 | Stage II | 0.162425909 | 0.175948 |
| HS3ST3B1 | GX | 0.497391304 | 0.013399 |
| BMP7 | Stage IV | -0.410502541 | 0.057741 |
| ITGA5 | GX | 0.493913043 | 0.014166 |
| BOP1 | G1 | 0.160902256 | 0.497974 |
| BOP1 | G2 | 0.053025297 | 0.53981 |
| BOP1 | G3 | 0.082858011 | 0.368274 |
| IL1B | GX | 0.477391304 | 0.018322 |
| BOP1 | Stage I | 0.09368566 | 0.234238 |
| BOP1 | Stage II | 0.049916582 | 0.679316 |
| BOP1 | Stage III | 0.039782891 | 0.792937 |
| BOP1 | Stage IV | 0.014116318 | 0.950285 |
| BOP1 | overall | 0.078193461 | 0.170361 |
| BRAF | G1 | 0.255639098 | 0.276664 |
| BRAF | G2 | -0.062760159 | 0.467923 |

| | | | |
|---|---|---|---|
| BRAF | G3 | 0.058316621 | 0.526943 |
| KRT19 | G1 | 0.473684211 | 0.03488 |
| BRAF | Stage I | -0.067571499 | 0.391425 |
| BRAF | Stage II | -0.161369562 | 0.178813 |
| FGFR2 | Stage IV | 0.469226426 | 0.02759 |
| BRAF | Stage IV | 0.242236025 | 0.277415 |
| BRAF | overall | -0.024483757 | 0.668138 |
| CAMK1D | G1 | 0.046616541 | 0.84527 |
| CAMK1D | G2 | 0.023441432 | 0.786479 |
| CAMK1D | G3 | -0.07248166 | 0.431445 |
| CAMK1D | GX | -0.38173913 | 0.065663 |
| CAMK1D | Stage I | -0.026551983 | 0.736536 |
| CAMK1D | Stage II | -0.165343438 | 0.168208 |
| CAMK1D | Stage III | -0.036082157 | 0.811832 |
| CAMK1D | Stage IV | -0.019762846 | 0.930438 |
| CAMK1D | overall | -0.054298868 | 0.341442 |
| CAV1 | G1 | 0.069172932 | 0.771985 |
| CDH13 | GX | 0.467826087 | 0.021148 |
| ITGB4 | GX | 0.467826087 | 0.021148 |
| IL18 | GX | 0.454782609 | 0.025562 |
| BOP1 | GX | 0.449565217 | 0.027522 |
| PIK3CA | G1 | 0.448120301 | 0.047532 |
| CAV1 | Stage III | 0.126503424 | 0.402172 |
| CAV1 | Stage IV | -0.059288538 | 0.793255 |
| ITGA6 | GX | 0.446956522 | 0.028546 |
| CBR1 | G1 | -0.030075188 | 0.899837 |
| WNT3A | Stage IV | 0.4432524 | 0.038816 |
| CBR1 | G3 | -0.001083474 | 0.990629 |
| CBR1 | GX | 0.174782609 | 0.414007 |
| CBR1 | Stage I | 0.092474721 | 0.240367 |
| CBR1 | Stage II | 0.227382859 | 0.056519 |
| CBR1 | Stage III | 0.189292544 | 0.207691 |
| CBR1 | Stage IV | -0.319028797 | 0.147851 |
| CBR1 | overall | 0.108283936 | 0.057258 |
| CD274 | G1 | -0.02556391 | 0.914804 |
| CD274 | G2 | 0.037515356 | 0.664568 |
| CD274 | G3 | 0.020478361 | 0.82431 |
| CD274 | GX | 0.283478261 | 0.179483 |
| CD274 | Stage I | 0.104054844 | 0.186218 |
| CD274 | Stage II | 0.131054083 | 0.275983 |
| CD274 | Stage III | -0.002282119 | 0.987991 |
| CD274 | Stage IV | -0.20609825 | 0.357472 |
| CD274 | overall | 0.048019808 | 0.40025 |
| CD44 | G1 | -0.082706767 | 0.728851 |

| | | | |
|---|---|---|---|
| CD44 | G2 | 0.152406283 | 0.07651 |
| YWHAZ | Stage III | 0.435884787 | 0.002462 |
| FOXC1 | GX | 0.424347826 | 0.03876 |
| VSNL1 | GX | 0.424347826 | 0.03876 |
| PTPRZ1 | Stage II | 0.424098123 | 2.28E-04 |
| CD44 | Stage III | -0.01683834 | 0.911562 |
| CD44 | Stage IV | -0.39582157 | 0.068228 |
| CD44 | Stage II | 0.421683616 | 0.00025 |
| CDH1 | G1 | 0.055639098 | 0.815774 |
| CDH1 | G2 | 0.121706046 | 0.158104 |
| BMP7 | Stage III | 0.416517612 | 0.003985 |
| FSCN1 | Stage II | 0.412796888 | 3.47E-04 |
| CDH1 | Stage I | 0.105684208 | 0.17938 |
| CDH1 | Stage II | 0.022937818 | 0.84941 |
| GSN | Stage III | 0.405785484 | 0.005143 |
| CDH1 | Stage IV | 0.14850367 | 0.509538 |
| CDH1 | overall | 0.098810131 | 0.082895 |
| CDH13 | G1 | -0.103759398 | 0.663326 |
| CDH13 | G2 | 0.102768269 | 0.233824 |
| RAC1 | Stage II | 0.402183117 | 5.08E-04 |
| SNAI2 | Stage II | 0.399349424 | 5.61E-04 |
| MMP13 | Stage II | 0.394117992 | 6.72E-04 |
| CDH13 | Stage II | 0.120155266 | 0.318232 |
| CDH13 | Stage III | 0.194720288 | 0.194712 |
| CDH13 | Stage IV | 0.103331451 | 0.647236 |
| CLDN1 | Stage II | 0.386723564 | 8.64E-04 |
| CDH2 | G1 | -0.009022556 | 0.969885 |
| CDH2 | G2 | -0.100287442 | 0.245366 |
| CDH2 | G3 | 0.024464991 | 0.790828 |
| CDH2 | GX | -0.086086957 | 0.68918 |
| CDH2 | Stage I | -0.005893975 | 0.940477 |
| CDH2 | Stage II | -0.08703963 | 0.470441 |
| CDH2 | Stage III | -0.104669093 | 0.488769 |
| CDH2 | Stage IV | 0.180124224 | 0.422475 |
| CDH2 | overall | -0.045600718 | 0.424433 |
| CDKN1B | G1 | -0.231578947 | 0.325899 |
| CDKN1B | G2 | -0.131469532 | 0.127094 |
| CDKN1B | G3 | -0.141261449 | 0.123811 |
| CDKN1B | GX | -0.386086957 | 0.062397 |
| HRAS | Stage II | 0.37859137 | 0.001132 |
| CDKN1B | Stage II | -0.155551271 | 0.195205 |
| CDKN1B | Stage III | -0.03867267 | 0.798593 |
| CDKN1B | Stage IV | 0.252399774 | 0.25712 |
| FOSL1 | Stage II | 0.374298913 | 0.001301 |

| | | | |
|---|---|---|---|
| CDKN2A | G1 | 0.084210526 | 0.724104 |
| CDKN2A | G2 | 0.04651074 | 0.590792 |
| CDKN2A | G3 | 0.097870383 | 0.287578 |
| CDKN2A | GX | 0.146956522 | 0.493193 |
| CDKN2A | Stage I | 0.116369289 | 0.13906 |
| CDKN2A | Stage II | 0.16125219 | 0.179133 |
| CDKN2A | Stage III | -0.094923827 | 0.530327 |
| CDKN2A | Stage IV | -0.102202146 | 0.650853 |
| CDKN2A | overall | 0.085335015 | 0.13447 |
| CDX2 | G1 | 0.077926102 | 0.744005 |
| CDX2 | G2 | -0.081406334 | 0.346113 |
| WNT3A | Stage II | 0.368631528 | 0.00156 |
| CDX2 | GX | -0.2495527 | 0.239591 |
| EGFR | Stage II | 0.364054025 | 0.001803 |
| CDX2 | Stage II | -0.006391657 | 0.95781 |
| PKP3 | Stage II | 0.348611239 | 0.002888 |
| CDX2 | Stage IV | -0.203336209 | 0.364093 |
| ID1 | Stage II | 0.345023013 | 0.003212 |
| CLDN1 | G1 | 0.431578947 | 0.057425 |
| CLDN1 | G3 | 0.343037723 | 1.25E-04 |
| CLDN1 | Stage I | 0.342762215 | 7.50E-06 |
| CLDN1 | GX | 0.383478261 | 0.064341 |
| LIMA1 | Stage II | 0.341015602 | 0.003612 |
| MMP14 | Stage I | 0.338317488 | 1.00E-05 |
| CLDN1 | Stage III | 0.231974343 | 0.120836 |
| CLDN1 | Stage IV | -0.341614907 | 0.119702 |
| PTPN14 | G3 | 0.336981378 | 1.68E-04 |
| CLDN4 | G1 | 0.222556391 | 0.345621 |
| CLDN4 | G2 | -0.057822358 | 0.503716 |
| CLDN4 | G3 | 0.104496245 | 0.256028 |
| IL18 | Stage II | 0.336421332 | 0.004124 |
| CLDN4 | Stage I | -0.029674931 | 0.706894 |
| CLDN4 | Stage II | -0.124028538 | 0.302767 |
| CLDN4 | Stage III | 0.192746563 | 0.199363 |
| CLDN4 | Stage IV | -0.210615471 | 0.346796 |
| CLDN4 | overall | -0.008481357 | 0.881958 |
| CLU | G1 | 0.264661654 | 0.259469 |
| CLU | G2 | 0.090294956 | 0.295825 |
| CLU | G3 | 0.120692799 | 0.189152 |
| CLU | GX | -0.114782609 | 0.593293 |
| CLU | Stage I | 0.122055437 | 0.120634 |
| CLU | Stage II | -0.100436791 | 0.404628 |
| CLU | Stage III | 0.155492507 | 0.302133 |
| CLU | Stage IV | 0.208356861 | 0.352111 |

| | | | |
|---|---|---|---|
| CLU | overall | 0.090100592 | 0.113965 |
| CMTM8 | G1 | 0.169924812 | 0.473849 |
| ITGA6 | Stage I | 0.329103822 | 1.79E-05 |
| CMTM8 | G3 | -0.151460694 | 0.098668 |
| CMTM8 | GX | -0.347826087 | 0.095813 |
| CMTM8 | Stage I | -0.151320256 | 0.053833 |
| PDPN | Stage II | 0.326344118 | 0.005478 |
| CMTM8 | Stage III | -0.066058102 | 0.66271 |
| CMTM8 | Stage IV | 0.008469791 | 0.970159 |
| CAV1 | Stage I | 0.324160309 | 2.43E-05 |
| COL8A1 | G1 | 0.058646617 | 0.805993 |
| COL8A1 | G2 | -0.152663908 | 0.076008 |
| COL8A1 | G3 | -0.012991275 | 0.888005 |
| COL8A1 | GX | -0.166956522 | 0.435538 |
| COL8A1 | Stage I | -0.036494429 | 0.643724 |
| COL8A1 | Stage II | -0.178472321 | 0.136454 |
| COL8A1 | Stage III | -0.04262012 | 0.778529 |
| COL8A1 | Stage IV | -0.051383399 | 0.820353 |
| COL8A1 | overall | -0.082330218 | 0.14879 |
| COL8A2 | G1 | -0.006015038 | 0.979921 |
| COL8A2 | G2 | -0.030638217 | 0.723275 |
| COL8A2 | G3 | 0.01941225 | 0.833321 |
| COL8A2 | GX | 0.13826087 | 0.519391 |
| COL8A2 | Stage I | 0.087234709 | 0.268173 |
| COL8A2 | Stage II | 0.127784438 | 0.288245 |
| EIF5A2 | Stage II | 0.322973868 | 0.006011 |
| COL8A2 | Stage IV | -0.005081875 | 0.982093 |
| COL8A2 | overall | 0.025417227 | 0.65628 |
| CSK | G1 | 0.015037594 | 0.949828 |
| CSK | G2 | 0.133282445 | 0.12189 |
| CSK | G3 | 0.015005426 | 0.870782 |
| CSK | GX | 0.035652174 | 0.868639 |
| CSK | Stage I | 0.096800295 | 0.218981 |
| CSK | Stage II | 0.029829224 | 0.804955 |
| CSK | Stage III | -0.045210634 | 0.765438 |
| CSK | Stage IV | 0.047995483 | 0.832029 |
| CSK | overall | 0.047090202 | 0.409444 |
| CSNK2B | G1 | 0.106766917 | 0.654142 |
| CSNK2B | G2 | 0.011154181 | 0.897451 |
| CSNK2B | G3 | -0.060063376 | 0.514618 |
| CSNK2B | GX | 0.393913043 | 0.056835 |
| CSNK2B | Stage I | 0.074257988 | 0.346158 |
| CSNK2B | Stage II | -0.004543968 | 0.97 |
| CSNK2B | Stage III | -0.285264913 | 0.054649 |

| | | | |
|---|---|---|---|
| CSNK2B | Stage IV | 0.104460757 | 0.643627 |
| CSNK2B | overall | -0.005936584 | 0.917222 |
| CTBP1 | G1 | 0.091729323 | 0.700516 |
| CTBP1 | G2 | -0.045499326 | 0.5989 |
| CTBP1 | G3 | -0.068567956 | 0.456793 |
| CTBP1 | GX | -0.371304348 | 0.074035 |
| CTBP1 | Stage I | -0.09994264 | 0.204319 |
| CTBP1 | Stage II | -0.154478156 | 0.198342 |
| CTBP1 | Stage III | -0.020415716 | 0.892873 |
| CTBP1 | Stage IV | 0.317899492 | 0.149371 |
| CTBP1 | overall | -0.064721192 | 0.25668 |
| CTGF | G1 | 0.02406015 | 0.9198 |
| CTGF | G2 | -0.118309221 | 0.170126 |
| CTGF | G3 | 0.02787863 | 0.762456 |
| CTGF | GX | -0.303478261 | 0.149409 |
| CTGF | Stage I | 0.015476187 | 0.844549 |
| RUNX3 | Stage II | 0.322621752 | 0.00607 |
| CTGF | Stage III | -0.108123112 | 0.474457 |
| CTGF | Stage IV | 0.279503106 | 0.207759 |
| CTGF | overall | -0.060118921 | 0.292128 |
| CTNNB1 | G1 | -0.108270677 | 0.649567 |
| CTNNB1 | G2 | -0.089073626 | 0.302432 |
| CTNNB1 | G3 | -0.170421496 | 0.062748 |
| CTNNB1 | GX | -0.26 | 0.219832 |
| CTNNB1 | Stage I | -0.13975953 | 0.075184 |
| CTNNB1 | Stage II | -0.21953571 | 0.065839 |
| CTNNB1 | Stage III | -0.16597792 | 0.270286 |
| CTNNB1 | Stage IV | -0.128176172 | 0.569723 |
| PRDX1 | Stage III | 0.32190218 | 0.029139 |
| CTNNBIP1 | G1 | -0.054135338 | 0.820675 |
| CTNNBIP1 | G2 | 0.129458631 | 0.133064 |
| CTNNBIP1 | G3 | 0.039963537 | 0.664746 |
| CTNNBIP1 | GX | -0.342608696 | 0.10124 |
| CTNNBIP1 | Stage I | 0.025208035 | 0.749417 |
| CTNNBIP1 | Stage II | 0.146329195 | 0.223348 |
| CTNNBIP1 | Stage III | 0.109850121 | 0.467385 |
| WNT3A | G3 | 0.319753996 | 3.71E-04 |
| CTNNBIP1 | overall | 0.056229259 | 0.324529 |
| CTNND1 | G1 | 0.103759398 | 0.663326 |
| RAC1 | Stage III | 0.318818235 | 0.030809 |
| CTNND1 | G3 | 0.114525329 | 0.212936 |
| CTNND1 | GX | 0.246956522 | 0.244676 |
| SDC1 | Stage II | 0.318228691 | 0.00684 |
| CTNND1 | Stage II | 0.231977129 | 0.05158 |

| | | | |
|---|---|---|---|
| CTNND1 | Stage III | 0.22321594 | 0.135923 |
| CTNND1 | Stage IV | -0.331451158 | 0.131845 |
| TP73 | Stage II | 0.317256181 | 0.007021 |
| CTSZ | G1 | -0.221052632 | 0.348974 |
| CTSZ | G2 | -0.087594671 | 0.310563 |
| CTSZ | G3 | -0.152186483 | 0.097045 |
| CTSZ | GX | -0.029565217 | 0.890923 |
| CTSZ | Stage I | -0.116862532 | 0.137381 |
| CTSZ | Stage II | -0.091281785 | 0.448999 |
| CTSZ | Stage III | -0.027817184 | 0.854399 |
| CTSZ | Stage IV | -0.216261999 | 0.333717 |
| CTSZ | overall | -0.103615772 | 0.068924 |
| CXCL12 | G1 | -0.227067669 | 0.335674 |
| CXCL12 | G2 | -0.107956538 | 0.210928 |
| CXCL12 | G3 | -0.101634762 | 0.269349 |
| CXCL12 | GX | 0.143478261 | 0.503593 |
| CXCL12 | Stage I | -0.026402348 | 0.737967 |
| MMP13 | Stage I | 0.314443188 | 4.35E-05 |
| CXCL12 | Stage III | -0.078393882 | 0.604557 |
| CXCL12 | Stage IV | -0.016374929 | 0.942342 |
| CXCL12 | overall | -0.077660427 | 0.173302 |
| CXCL16 | G1 | -0.012030075 | 0.959853 |
| CXCL16 | G2 | -0.096575743 | 0.263363 |
| CXCL16 | G3 | -0.030816513 | 0.738286 |
| CXCL16 | GX | -0.086086957 | 0.68918 |
| CXCL16 | Stage I | -0.097470883 | 0.215791 |
| CXCL16 | Stage II | 0.092924991 | 0.440842 |
| CXCL16 | Stage III | -0.01178067 | 0.938063 |
| CXCL16 | Stage IV | -0.108977979 | 0.629268 |
| CXCL16 | overall | -0.038947457 | 0.495164 |
| CXCL5 | G1 | -0.105263158 | 0.658728 |
| CXCL5 | G2 | -0.094137853 | 0.275662 |
| CXCL5 | G3 | -0.005601424 | 0.951583 |
| CXCL5 | GX | -0.035659927 | 0.868611 |
| CXCL5 | Stage I | 0.093186135 | 0.236753 |
| CXCL5 | Stage II | -0.154394319 | 0.198589 |
| CXCL5 | Stage III | -0.226176527 | 0.130674 |
| CXCL5 | Stage IV | -0.067193676 | 0.766386 |
| CXCL5 | overall | -0.042724901 | 0.454259 |
| CXCR4 | G1 | -0.216541353 | 0.359146 |
| CXCR4 | G2 | -0.072924394 | 0.398831 |
| TMPRSS4 | Stage III | 0.312773702 | 0.034313 |
| CXCR4 | GX | -0.393043478 | 0.057433 |
| CXCR4 | Stage I | -0.108998357 | 0.166047 |

| | | | |
|---|---|---|---|
| CXCR4 | Stage II | -0.159491612 | 0.18399 |
| CXCR4 | Stage III | -0.199284526 | 0.184251 |
| CXCR4 | Stage IV | -0.097684924 | 0.665396 |
| CDH1 | Stage III | 0.312156913 | 0.034688 |
| CYR61 | G1 | 0.052631579 | 0.825581 |
| CYR61 | G2 | -0.058693033 | 0.497303 |
| CYR61 | G3 | 0.034528802 | 0.708109 |
| CYR61 | GX | -0.113043478 | 0.59894 |
| CYR61 | Stage I | 0.052807466 | 0.50319 |
| CYR61 | Stage II | -0.119836685 | 0.319526 |
| CYR61 | Stage III | -0.182877939 | 0.223797 |
| CYR61 | Stage IV | 0.417278374 | 0.05334 |
| CYR61 | overall | -0.025300289 | 0.657761 |
| DAB2 | G1 | -0.198496241 | 0.401504 |
| BMP7 | G2 | 0.311310425 | 2.25E-04 |
| DAB2 | G3 | -0.05758736 | 0.532133 |
| DAB2 | GX | 0.188695652 | 0.377216 |
| DAB2 | Stage I | -0.094281431 | 0.231264 |
| DAB2 | Stage II | -0.072351378 | 0.548767 |
| DAB2 | Stage III | -0.046197496 | 0.760468 |
| DAB2 | Stage IV | 0.015245624 | 0.946313 |
| DAB2 | overall | -0.093375668 | 0.101354 |
| DAPK1 | G1 | -0.042105263 | 0.860095 |
| VSNL1 | Stage II | 0.309476102 | 0.008634 |
| DAPK1 | G3 | -0.021197205 | 0.818248 |
| DAPK1 | GX | -0.324347826 | 0.122033 |
| DAPK1 | Stage I | -0.150627499 | 0.054953 |
| DAPK1 | Stage II | -0.181624594 | 0.129552 |
| DAPK1 | Stage III | -0.135508544 | 0.369227 |
| DAPK1 | Stage IV | 0.299830604 | 0.175203 |
| PTPN14 | Stage II | 0.309274893 | 0.00868 |
| DDR2 | G1 | -0.097744361 | 0.681832 |
| DDR2 | G2 | -0.065243372 | 0.450463 |
| DDR2 | G3 | 0.052551982 | 0.568646 |
| DDR2 | GX | -0.030434783 | 0.887734 |
| DDR2 | Stage I | 0.051904111 | 0.510531 |
| DDR2 | Stage II | -0.161554004 | 0.17831 |
| DDR2 | Stage III | -0.035588725 | 0.814359 |
| DDR2 | Stage IV | 0.162055336 | 0.471204 |
| DDR2 | overall | -0.015314805 | 0.788596 |
| DDX5 | G1 | 0.013533835 | 0.95484 |
| DDX5 | G2 | -0.066906003 | 0.43898 |
| DDX5 | G3 | -0.134253592 | 0.143766 |
| DDX5 | GX | -0.353913043 | 0.089758 |

| | | | |
|---|---|---|---|
| DDX5 | Stage I | -0.09303724 | 0.237506 |
| SP1 | Stage III | 0.309072968 | 0.036616 |
| DDX5 | Stage III | -0.20915315 | 0.163019 |
| DDX5 | Stage IV | 0.277244495 | 0.211611 |
| THBD | Stage I | 0.308861468 | 6.03E-05 |
| DLX4 | G1 | 0.237593985 | 0.313131 |
| DLX4 | G2 | -0.052054435 | 0.547268 |
| DLX4 | G3 | -0.116973564 | 0.203252 |
| DLX4 | GX | 0.147826087 | 0.49061 |
| DLX4 | Stage I | -0.134541686 | 0.086844 |
| DLX4 | Stage II | 0.046546333 | 0.699901 |
| DLX4 | Stage III | -0.02917412 | 0.847379 |
| DLX4 | Stage IV | 0.001693958 | 0.994031 |
| DLX4 | overall | -0.045579364 | 0.42465 |
| DNAJB6 | G1 | 0.207518797 | 0.379993 |
| DNAJB6 | G2 | 0.015445535 | 0.858352 |
| DNAJB6 | G3 | 0.008831011 | 0.923736 |
| DNAJB6 | GX | -0.043478261 | 0.840133 |
| DNAJB6 | Stage I | 0.035106144 | 0.656397 |
| DNAJB6 | Stage II | 0.064755741 | 0.591599 |
| DNAJB6 | Stage III | 0.160796893 | 0.285733 |
| DNAJB6 | Stage IV | -0.001693958 | 0.994031 |
| DNAJB6 | overall | 0.042492202 | 0.456723 |
| ECT2 | G1 | 0.305263158 | 0.190609 |
| ECT2 | G2 | 0.013406009 | 0.876897 |
| ECT2 | G3 | -0.005889656 | 0.949095 |
| ECT2 | GX | -0.217391304 | 0.307533 |
| ECT2 | Stage I | 0.00679733 | 0.931375 |
| ECT2 | Stage II | 0.095825753 | 0.426647 |
| ECT2 | Stage III | 0.001171899 | 0.993833 |
| ECT2 | Stage IV | -0.037831733 | 0.867254 |
| ECT2 | overall | 0.019038923 | 0.738872 |
| TGFB1 | Stage II | 0.303456602 | 0.010095 |
| EDN1 | G2 | -0.060451081 | 0.484485 |
| EDN1 | G3 | 0.099697009 | 0.278632 |
| EDN1 | GX | -0.255652174 | 0.227918 |
| EDN1 | Stage I | 0.061613791 | 0.434614 |
| EDN1 | Stage II | -0.05601992 | 0.642635 |
| EDN1 | Stage III | 0.029544193 | 0.845466 |
| EDN1 | Stage IV | 0.162055336 | 0.471204 |
| EDN1 | overall | 0.008376622 | 0.883406 |
| EDNRA | G1 | -0.127819549 | 0.591246 |
| EDNRA | G2 | -0.129692401 | 0.132359 |
| EDNRA | G3 | 0.019773408 | 0.830266 |

| | | | |
|---|---|---|---|
| EDNRA | GX | -0.053043478 | 0.805559 |
| EDNRA | Stage I | 0.034443869 | 0.662478 |
| EDNRA | Stage II | -0.118948013 | 0.323154 |
| EDNRA | Stage III | -0.183988159 | 0.220949 |
| EDNRA | Stage IV | -0.079616036 | 0.724693 |
| EDNRA | overall | -0.056442188 | 0.322697 |
| EGFR | G1 | 0.042105263 | 0.860095 |
| EGFR | G2 | 0.156151378 | 0.069468 |
| SNAI2 | G3 | 0.301792769 | 8.09E-04 |
| BRAF | Stage III | 0.301054711 | 0.042043 |
| GSK3B | Stage III | 0.300931354 | 0.042131 |
| SMAD3 | Stage I | 0.300240802 | 9.86E-05 |
| EGFR | Stage III | 0.025473386 | 0.866551 |
| EGFR | Stage IV | 0.005081875 | 0.982093 |
| SNAI2 | Stage I | 0.298763845 | 1.07E-04 |
| EGR1 | G1 | 0.212030075 | 0.369486 |
| EGR1 | G2 | 0.077144186 | 0.372031 |
| EGR1 | G3 | 0.017453662 | 0.849929 |
| EGR1 | GX | -0.137391304 | 0.522047 |
| EGR1 | Stage I | 0.065343594 | 0.407265 |
| EGR1 | Stage II | -0.046328356 | 0.70124 |
| EGR1 | Stage III | 0.140812929 | 0.350604 |
| EGR1 | Stage IV | 0.228684359 | 0.305996 |
| EGR1 | overall | 0.055864413 | 0.327683 |
| EIF5A2 | G1 | -0.25112782 | 0.285522 |
| SMAD3 | Stage III | 0.29834084 | 0.044022 |
| NDRG1 | Stage II | 0.296598731 | 0.012018 |
| NOTCH1 | G3 | 0.296080219 | 0.001026 |
| CLDN1 | overall | 0.294908776 | 1.29E-07 |
| VDR | Stage II | 0.294100386 | 0.012793 |
| EIF5A2 | Stage III | 0.00339234 | 0.982149 |
| EIF5A2 | Stage IV | 0.229813665 | 0.303548 |
| LYPD3 | Stage III | 0.292789739 | 0.048307 |
| ELF5 | G1 | 0.431578947 | 0.057425 |
| ELF5 | G2 | 0.034206226 | 0.692589 |
| ELF5 | G3 | -0.130652356 | 0.154925 |
| WNT3A | G2 | 0.289798791 | 6.21E-04 |
| AXL | Stage II | 0.286286773 | 0.015505 |
| ELF5 | Stage II | -0.129196667 | 0.282905 |
| ELF5 | Stage III | 0.225822406 | 0.131294 |
| ELF5 | Stage IV | 0.089805143 | 0.691043 |
| ELF5 | overall | -0.065018487 | 0.254497 |
| ENG | G1 | -0.190977444 | 0.419927 |
| FGFR2 | G3 | 0.285752486 | 0.001558 |

| | | | |
|---|---|---|---|
| ENG | G3 | -0.097950254 | 0.287183 |
| ENG | GX | 0.086086957 | 0.68918 |
| ENG | Stage I | -0.048019131 | 0.542723 |
| ENG | Stage II | -0.232010664 | 0.051545 |
| ENG | Stage III | -0.18731882 | 0.212558 |
| ENG | Stage IV | 0.008469791 | 0.970159 |
| ENG | overall | -0.110244426 | 0.052871 |
| EPAS1 | G1 | -0.058646617 | 0.805993 |
| EPAS1 | G2 | 0.076874635 | 0.373709 |
| EPAS1 | G3 | 0.121581803 | 0.18589 |
| EPAS1 | GX | 0.36 | 0.083995 |
| EPAS1 | Stage I | 0.153656232 | 0.050195 |
| EPAS1 | Stage II | 0.121530194 | 0.312685 |
| EPAS1 | Stage III | 0.135385186 | 0.369667 |
| EPAS1 | Stage IV | -0.389045737 | 0.073529 |
| EPAS1 | overall | 0.089949691 | 0.114574 |
| EPB41L5 | G1 | 0.010526316 | 0.964869 |
| EPB41L5 | G2 | -0.150660162 | 0.079985 |
| EPB41L5 | G3 | 0.01693276 | 0.854358 |
| EPB41L5 | GX | -0.226956522 | 0.2862 |
| EPB41L5 | Stage I | -0.097714734 | 0.214639 |
| EPB41L5 | Stage II | -0.168344805 | 0.16051 |
| EPB41L5 | Stage III | 0.163880838 | 0.27647 |
| EPB41L5 | Stage IV | 0.095426313 | 0.672712 |
| EPB41L5 | overall | -0.079100696 | 0.165441 |
| ERBB2 | G1 | 0.064661654 | 0.786515 |
| ERBB2 | G2 | -0.037186169 | 0.667336 |
| FSCN1 | Stage I | 0.283880657 | 2.40E-04 |
| WNT3A | overall | 0.28362053 | 3.98E-07 |
| SPRR2A | G2 | 0.281661717 | 8.94E-04 |
| ERBB2 | Stage II | -0.203874949 | 0.088129 |
| ERBB2 | Stage III | 0.022019367 | 0.884513 |
| ERBB2 | Stage IV | -0.238848108 | 0.284397 |
| CAV1 | Stage II | 0.280468481 | 0.017832 |
| ERF | G1 | 0.084210526 | 0.724104 |
| ERF | G2 | -0.110497 | 0.200321 |
| ERF | G3 | 0.012147415 | 0.895237 |
| ERF | GX | -0.08 | 0.710196 |
| ERF | Stage I | 0.01572558 | 0.842077 |
| ERF | Stage II | -0.091466226 | 0.44808 |
| ERF | Stage III | -0.171158948 | 0.255402 |
| ERF | Stage IV | 0.33822699 | 0.123656 |
| ERF | overall | -0.038420728 | 0.50102 |
| ESR1 | G1 | -0.069172932 | 0.771985 |

| | | | |
|---|---|---|---|
| ESR1 | G2 | -0.056911132 | 0.510474 |
| ESR1 | G3 | -0.014185875 | 0.877783 |
| ESR1 | GX | -0.05826087 | 0.786844 |
| ESR1 | Stage I | -0.051532794 | 0.513564 |
| ESR1 | Stage II | -0.001995322 | 0.986824 |
| ESR1 | Stage III | -0.135878617 | 0.367909 |
| ESR1 | Stage IV | 0.267080745 | 0.229528 |
| ESR1 | overall | -0.035383188 | 0.535486 |
| ETV4 | G1 | 0.007518797 | 0.974903 |
| ETV4 | G2 | 0.154107081 | 0.073244 |
| ETV4 | G3 | -0.053906325 | 0.55871 |
| ETV4 | GX | -0.108695652 | 0.613156 |
| ETV4 | Stage I | 0.005791447 | 0.94151 |
| ETV4 | Stage II | 0.003437319 | 0.977304 |
| ETV4 | Stage III | 0.119348672 | 0.429519 |
| ETV4 | Stage IV | 0.184641446 | 0.410735 |
| ETV4 | overall | 0.0503545 | 0.37771 |
| EZH2 | G1 | 0.338345865 | 0.144526 |
| EZH2 | G2 | 0.079503357 | 0.357541 |
| EZH2 | G3 | 0.014137257 | 0.878199 |
| EZH2 | GX | -0.363478261 | 0.080829 |
| EZH2 | Stage I | 0.078882833 | 0.316866 |
| EZH2 | Stage II | -0.13120499 | 0.275426 |
| EZH2 | Stage III | 0.046814285 | 0.757366 |
| EZH2 | Stage IV | 0.209486166 | 0.349447 |
| EZH2 | overall | 0.026150174 | 0.647029 |
| FBLN5 | G1 | -0.054135338 | 0.820675 |
| PTHLH | Stage II | 0.278053974 | 0.018883 |
| FBLN5 | G3 | -0.020443634 | 0.824604 |
| FBLN5 | GX | -0.143478261 | 0.503593 |
| FBLN5 | Stage I | -0.036549849 | 0.64322 |
| PTPRZ1 | G3 | 0.276900237 | 0.002201 |
| FBLN5 | Stage III | -0.226299885 | 0.130459 |
| FBLN5 | Stage IV | 0.214003388 | 0.338913 |
| FBLN5 | overall | -0.085777752 | 0.132453 |
| FGFR1 | G1 | -0.117293233 | 0.622379 |
| FGFR1 | G2 | -0.144102668 | 0.09418 |
| FGFR1 | G3 | 0.015668707 | 0.865123 |
| FGFR1 | GX | -0.339130435 | 0.104983 |
| FGFR1 | Stage I | -0.026380179 | 0.738179 |
| WNT3A | Stage I | 0.276841054 | 3.47E-04 |
| FGFR1 | Stage III | -0.099734781 | 0.509597 |
| FGFR1 | Stage IV | 0.319028797 | 0.147851 |
| FGFR1 | overall | -0.085996782 | 0.131464 |

| | | | |
|---|---|---|---|
| EIF5A2 | G3 | 0.276522117 | 0.002234 |
| FGFR2 | G2 | -0.019042735 | 0.825836 |
| MSN | Stage II | 0.274113633 | 0.02071 |
| FGFR2 | GX | -0.366956522 | 0.077754 |
| FGFR2 | Stage I | 0.147036248 | 0.061071 |
| FGFR2 | Stage II | -0.16928378 | 0.158156 |
| FGFR2 | Stage III | 0.229013756 | 0.125787 |
| CBR1 | G2 | 0.273678184 | 0.001264 |
| FOSL1 | G2 | 0.272886228 | 0.001308 |
| FHL2 | G1 | 0.091729323 | 0.700516 |
| FHL2 | G2 | -0.038965686 | 0.652429 |
| FHL2 | G3 | -0.034504493 | 0.708305 |
| FHL2 | GX | 0.093913043 | 0.662485 |
| FHL2 | Stage I | 0.091128002 | 0.247313 |
| FHL2 | Stage II | 0.076492928 | 0.526061 |
| RAC1 | G2 | 0.272850447 | 0.00131 |
| FHL2 | Stage IV | 0.002823264 | 0.990051 |
| FHL2 | overall | -0.019404989 | 0.734039 |
| FLT1 | G1 | -0.120300752 | 0.613418 |
| SMAD3 | G2 | 0.272304188 | 0.001341 |
| SMAD3 | G3 | 0.271302687 | 0.002724 |
| FLT1 | GX | -0.072173913 | 0.737519 |
| YWHAZ | Stage II | 0.270223593 | 0.022661 |
| FLT1 | Stage II | -0.160028169 | 0.1825 |
| FLT1 | Stage III | -0.218898417 | 0.143858 |
| FLT1 | Stage IV | -0.228684359 | 0.305996 |
| ANXA1 | Stage I | 0.269828778 | 4.95E-04 |
| FN1 | G1 | -0.040601504 | 0.865047 |
| FN1 | G2 | -0.067015732 | 0.438228 |
| FN1 | G3 | -0.036779095 | 0.690031 |
| FN1 | GX | -0.153043478 | 0.475255 |
| FN1 | Stage I | 0.036037209 | 0.647887 |
| FN1 | Stage II | -0.10011821 | 0.406128 |
| FN1 | Stage III | -0.232591132 | 0.119823 |
| FN1 | Stage IV | -0.106719368 | 0.636432 |
| FN1 | overall | -0.052016239 | 0.362151 |
| FOSL1 | G1 | 0.085714286 | 0.719366 |
| IL18 | Stage I | 0.267556536 | 5.55E-04 |
| FOSL1 | G3 | 0.16049312 | 0.079932 |
| FSCN1 | G2 | 0.266426535 | 0.001717 |
| JAG2 | Stage II | 0.265796997 | 0.025068 |
| IL18 | G3 | 0.265215089 | 0.003417 |
| FOSL1 | Stage III | 0.091593167 | 0.544917 |
| FOSL1 | Stage IV | 0.023150762 | 0.918549 |

| | | | |
|---|---|---|---|
| EIF5A2 | Stage I | 0.265131887 | 6.25E-04 |
| FOXA1 | G1 | 0.380451128 | 0.097968 |
| FOXA1 | G2 | 0.014655964 | 0.865523 |
| FOXA1 | G3 | -0.118880063 | 0.195933 |
| FOXA1 | GX | -0.38 | 0.067006 |
| FOXA1 | Stage I | -6.60E-04 | 0.993334 |
| FOXA1 | Stage II | -0.113381232 | 0.346474 |
| FOXA1 | Stage III | -0.072596065 | 0.631616 |
| FOXA1 | Stage IV | -0.151891587 | 0.499815 |
| FOXA1 | overall | -0.065178572 | 0.253327 |
| FOXC1 | G1 | 0.389473684 | 0.089619 |
| FOXC1 | G2 | -0.055224646 | 0.523104 |
| FOXC1 | G3 | 0.042397882 | 0.645666 |
| THBD | G3 | 0.263843383 | 0.003594 |
| FOXC1 | Stage I | 0.058651563 | 0.457064 |
| FOXC1 | Stage II | 0.150621652 | 0.209915 |
| FOXC1 | Stage III | -0.202491829 | 0.177144 |
| FOXC1 | Stage IV | 0.090909091 | 0.687429 |
| FOXC1 | overall | 0.037044317 | 0.516491 |
| FOXM1 | G1 | 0.034586466 | 0.884902 |
| FOXM1 | G2 | -0.100354234 | 0.24505 |
| FOXM1 | G3 | -0.03429266 | 0.710016 |
| FOXM1 | GX | -0.247826087 | 0.242965 |
| FOXM1 | Stage I | -0.083942728 | 0.286721 |
| FOXM1 | Stage II | 0.075654558 | 0.530619 |
| FOXM1 | Stage III | -0.113427497 | 0.452919 |
| FOXM1 | Stage IV | -0.156408809 | 0.486995 |
| FOXM1 | overall | -0.054651716 | 0.338309 |
| FOXQ1 | G1 | 0.345864662 | 0.135252 |
| FOXQ1 | G2 | 0.007137149 | 0.934277 |
| FOXQ1 | G3 | 0.089508183 | 0.330949 |
| FOXQ1 | GX | 0.22 | 0.30162 |
| FOXQ1 | Stage I | 0.032795107 | 0.67771 |
| FOXQ1 | Stage II | 0.10808273 | 0.369619 |
| FOXQ1 | Stage III | 0.152901993 | 0.310357 |
| FOXQ1 | Stage IV | -0.131564088 | 0.559479 |
| FOXQ1 | overall | 0.069356612 | 0.224099 |
| FSCN1 | G1 | -0.066165414 | 0.781664 |
| CAV1 | G3 | 0.262044538 | 0.003838 |
| FSCN1 | G3 | 0.169299822 | 0.064524 |
| FOSL1 | Stage I | 0.261382687 | 7.51E-04 |
| GATA3 | Stage II | 0.261035052 | 0.027896 |
| LYPD3 | Stage II | 0.260632634 | 0.028146 |
| FSCN1 | Stage III | -0.103312158 | 0.494452 |

| | | | |
|---|---|---|---|
| FSCN1 | Stage IV | -0.047995483 | 0.832029 |
| JAG2 | Stage I | 0.260102473 | 7.99E-04 |
| GAB2 | G1 | 0.178947368 | 0.450325 |
| GAB2 | G2 | -0.153909092 | 0.073618 |
| HSPB1 | Stage II | 0.259375079 | 0.028942 |
| GAB2 | GX | -0.233043478 | 0.27312 |
| ITGA6 | Stage II | 0.259140335 | 0.029093 |
| CDH13 | G3 | 0.258571863 | 0.004351 |
| GAB2 | Stage III | 6.17E-05 | 0.999675 |
| GAB2 | Stage IV | 0.024280068 | 0.91459 |
| PTPRZ1 | overall | 0.25851735 | 4.14E-06 |
| GATA3 | G1 | 0.270676692 | 0.248391 |
| GATA3 | G2 | -0.132991424 | 0.122714 |
| GATA3 | G3 | 0.090845162 | 0.323748 |
| GATA3 | GX | 0.162608696 | 0.447754 |
| GATA3 | Stage I | -0.051000756 | 0.517927 |
| IL18 | overall | 0.258140636 | 4.28E-06 |
| GATA3 | Stage III | -0.224202802 | 0.134156 |
| GATA3 | Stage IV | -0.162055336 | 0.471204 |
| GATA3 | overall | -3.58E-04 | 0.995006 |
| GIPC2 | G1 | 0.237593985 | 0.313131 |
| GIPC2 | G2 | -0.023272068 | 0.787985 |
| GIPC2 | G3 | 0.11538308 | 0.209506 |
| GIPC2 | GX | -0.190434783 | 0.372753 |
| GIPC2 | Stage I | 0.037863316 | 0.631328 |
| GIPC2 | Stage II | -0.045858869 | 0.704127 |
| GIPC2 | Stage III | 0.024609881 | 0.871036 |
| GIPC2 | Stage IV | -0.079616036 | 0.724693 |
| GIPC2 | overall | 0.018399526 | 0.747339 |
| GLRX | G1 | -0.368421053 | 0.109964 |
| GLRX | G2 | -0.11789893 | 0.171623 |
| PDPN | Stage I | 0.25710145 | 9.23E-04 |
| GLRX | GX | 0.012173913 | 0.954977 |
| GLRX | Stage I | -0.132222336 | 0.092471 |
| GLRX | Stage II | -0.132680522 | 0.270015 |
| GLRX | Stage III | -0.1379757 | 0.360492 |
| GLRX | Stage IV | -0.15866742 | 0.480647 |
| SNAI2 | overall | 0.256638339 | 4.88E-06 |
| GMNN | G1 | 0.222556391 | 0.345621 |
| GMNN | G2 | 0.002945982 | 0.972846 |
| GMNN | G3 | -3.06E-04 | 0.997357 |
| GMNN | GX | 0.167826087 | 0.433117 |
| GMNN | Stage I | -0.045874356 | 0.560915 |
| GMNN | Stage II | 0.203053346 | 0.089445 |

| | | | |
|---|---|---|---|
| GMNN | Stage III | -0.094800469 | 0.530864 |
| GMNN | Stage IV | -0.121400339 | 0.590455 |
| GMNN | overall | 0.006017729 | 0.916095 |
| GREM1 | G1 | 0 | 1 |
| GREM1 | G2 | 0.015161671 | 0.860929 |
| GREM1 | G3 | -0.001264053 | 0.989068 |
| GREM1 | GX | 0.157391304 | 0.462649 |
| GREM1 | Stage I | 0.075144717 | 0.340413 |
| GREM1 | Stage II | 0.008148962 | 0.946226 |
| GREM1 | Stage III | 0.010547092 | 0.944538 |
| GREM1 | Stage IV | -0.114624506 | 0.6115 |
| GREM1 | overall | 0.032751575 | 0.566279 |
| GSK3A | G1 | -0.091729323 | 0.700516 |
| GSK3A | G2 | -0.050418043 | 0.559951 |
| GSK3A | G3 | -0.111452012 | 0.225552 |
| GSK3A | GX | -0.227826087 | 0.284307 |
| GSK3A | Stage I | -0.084084051 | 0.285907 |
| GSK3A | Stage II | -0.053135925 | 0.659868 |
| GSK3A | Stage III | -0.125146488 | 0.407279 |
| GSK3A | Stage IV | 0.166572558 | 0.458761 |
| GSK3A | overall | -0.072032356 | 0.206689 |
| GSK3B | G1 | 0.203007519 | 0.390666 |
| GSK3B | G2 | 0.146330642 | 0.089149 |
| GSK3B | G3 | 0.112472978 | 0.221304 |
| GSK3B | GX | -0.16173913 | 0.450219 |
| GSK3B | Stage I | 0.077987791 | 0.322405 |
| GSK3B | Stage II | 0.117690457 | 0.328333 |
| SPRR2A | Stage I | 0.255108121 | 0.001014 |
| GSK3B | Stage IV | -0.079616036 | 0.724693 |
| GSK3B | overall | 0.104258423 | 0.067211 |
| GSN | G1 | -0.018045113 | 0.939809 |
| GSN | G2 | 0.09931658 | 0.249989 |
| GSN | G3 | 0.020526978 | 0.8239 |
| GSN | GX | -0.312173913 | 0.137518 |
| GSN | Stage I | 0.052840719 | 0.502921 |
| GSN | Stage II | -0.227500231 | 0.056388 |
| TGFB1 | Stage I | 0.254108741 | 0.001063 |
| GSN | Stage IV | -0.270468662 | 0.223449 |
| GSN | overall | 0.029709968 | 0.602888 |
| HAS2 | G1 | -0.252631579 | 0.28255 |
| HAS2 | G2 | 0.031153465 | 0.718815 |
| HAS2 | G3 | 0.032056257 | 0.728161 |
| HAS2 | GX | 0.24 | 0.258646 |
| IGF1R | G3 | 0.253654556 | 0.005183 |

| | | | |
|---|---|---|---|
| HAS2 | Stage II | 0.096563519 | 0.423079 |
| ITGA6 | G3 | 0.249497765 | 0.005994 |
| HAS2 | Stage IV | -0.241106719 | 0.27973 |
| HAS2 | overall | 0.046229742 | 0.418064 |
| HBEGF | G1 | -0.07518797 | 0.752727 |
| TNC | Stage I | 0.247987541 | 0.001414 |
| HBEGF | G3 | 0.09150497 | 0.320232 |
| JAG1 | Stage I | 0.247671644 | 0.001435 |
| SMAD3 | overall | 0.247017904 | 1.12E-05 |
| HBEGF | Stage II | 0.24648094 | 0.038255 |
| HBEGF | Stage III | 0.159193241 | 0.290628 |
| HBEGF | Stage IV | 0.024280068 | 0.91459 |
| HSPB1 | G2 | 0.245838054 | 0.003916 |
| HDAC6 | G1 | 0.198496241 | 0.401504 |
| HDAC6 | G2 | -0.123759885 | 0.151151 |
| HDAC6 | G3 | -0.015873595 | 0.863377 |
| TMPRSS4 | G3 | 0.245000652 | 0.006997 |
| HDAC6 | Stage I | -0.007639722 | 0.922895 |
| IGF1R | Stage I | 0.244537613 | 0.001656 |
| HDAC6 | Stage III | -0.120212176 | 0.426164 |
| HDAC6 | Stage IV | -0.199322417 | 0.37384 |
| HDAC6 | overall | -0.087383564 | 0.125332 |
| HDGF | G1 | 0.22406015 | 0.342286 |
| HDGF | G2 | -0.012122658 | 0.888602 |
| HDGF | G3 | 0.168226766 | 0.066261 |
| HDGF | GX | -0.048695652 | 0.821234 |
| HDGF | Stage I | 0.033036187 | 0.675475 |
| HDGF | Stage II | 0.14299248 | 0.234198 |
| HDGF | Stage III | 0.005982853 | 0.968523 |
| HDGF | Stage IV | 0.114624506 | 0.6115 |
| HDGF | overall | 0.062927059 | 0.270128 |
| HIF1A | G1 | -0.314285714 | 0.17717 |
| HIF1A | G2 | -0.045246473 | 0.600934 |
| HIF1A | G3 | -0.062150454 | 0.500089 |
| HIF1A | GX | 0.140869565 | 0.511462 |
| HIF1A | Stage I | -0.015345949 | 0.845841 |
| HIF1A | Stage II | 0.024832535 | 0.837134 |
| HIF1A | Stage III | -0.018318633 | 0.903823 |
| HIF1A | Stage IV | -0.241106719 | 0.27973 |
| HIF1A | overall | -0.063853004 | 0.263129 |
| HMGB1 | G1 | 0.203007519 | 0.390666 |
| HMGB1 | G2 | -0.102985342 | 0.232832 |
| HMGB1 | G3 | -0.115938708 | 0.207306 |
| HMGB1 | GX | -0.177391304 | 0.406963 |

| | | | |
|---|---|---|---|
| HMGB1 | Stage I | -0.041909016 | 0.595298 |
| HMGB1 | Stage II | -0.152466067 | 0.204322 |
| HMGB1 | Stage III | -0.042003331 | 0.781655 |
| HMGB1 | Stage IV | -0.333709768 | 0.129073 |
| HMGB1 | overall | -0.092275231 | 0.105462 |
| HMGB3 | G1 | 0.332330827 | 0.152259 |
| HMGB3 | G2 | 0.087277411 | 0.312325 |
| HMGB3 | G3 | -0.031017928 | 0.736638 |
| HMGB3 | GX | 0.13826087 | 0.519391 |
| HMGB3 | Stage I | 0.084416574 | 0.284 |
| HMGB3 | Stage II | 0.020942496 | 0.862375 |
| HMGB3 | Stage III | 0.070375625 | 0.642109 |
| HMGB3 | Stage IV | 0.12365895 | 0.583509 |
| HMGB3 | overall | 0.054518711 | 0.339488 |
| HMOX1 | G1 | -0.069172932 | 0.771985 |
| HMOX1 | G2 | -0.007077514 | 0.934824 |
| HMOX1 | G3 | 0.015762469 | 0.864324 |
| HMOX1 | GX | 0.050434783 | 0.814956 |
| HMOX1 | Stage I | 0.002635247 | 0.973367 |
| HMOX1 | Stage II | -0.053186227 | 0.659566 |
| HMOX1 | Stage III | 0.109726763 | 0.467889 |
| HMOX1 | Stage IV | 0.164313947 | 0.464961 |
| HMOX1 | overall | 0.022676608 | 0.691328 |
| HNF4A | G1 | -0.213533835 | 0.366021 |
| HNF4A | G2 | -0.046866726 | 0.587951 |
| HNF4A | G3 | -0.016872775 | 0.854868 |
| HNF4A | GX | 0.145098384 | 0.498735 |
| HNF4A | Stage I | -0.114851645 | 0.144324 |
| HNF4A | Stage II | 0.116772783 | 0.332145 |
| HNF4A | Stage III | -0.011412004 | 0.939998 |
| HNF4A | Stage IV | -0.115884755 | 0.607563 |
| HNF4A | overall | -0.032774901 | 0.566002 |
| HOXA10 | G1 | -0.142857143 | 0.54795 |
| HOXA10 | G2 | -0.074596567 | 0.388076 |
| HOXA10 | G3 | 0.042102704 | 0.647968 |
| HOXA10 | GX | 0.120869565 | 0.57371 |
| HOXA10 | Stage I | -0.004982307 | 0.94967 |
| HOXA10 | Stage II | -0.119467802 | 0.321029 |
| HOXA10 | Stage III | 0.082464689 | 0.585864 |
| HOXA10 | Stage IV | 0.090909091 | 0.687429 |
| HOXA10 | overall | -0.00587659 | 0.918056 |
| HOXB7 | G1 | -0.003007519 | 0.98996 |
| HOXB7 | G2 | 0.049859857 | 0.564309 |
| HOXB7 | G3 | 0.041262317 | 0.654539 |

| | | | |
|---|---|---|---|
| HOXB7 | GX | -0.233043478 | 0.27312 |
| HOXB7 | Stage I | -0.058515782 | 0.458108 |
| HOXB7 | Stage II | 0.173576237 | 0.14772 |
| HOXB7 | Stage III | 0.13908592 | 0.356603 |
| HOXB7 | Stage IV | 0.059288538 | 0.793255 |
| HOXB7 | overall | 0.036650185 | 0.520967 |
| HOXB9 | G1 | -0.013533835 | 0.95484 |
| HOXB9 | G2 | -0.076636094 | 0.375198 |
| HOXB9 | G3 | -0.050405869 | 0.58456 |
| HOXB9 | GX | -0.216521739 | 0.30952 |
| HOXB9 | Stage I | -0.117264331 | 0.136025 |
| HOXB9 | Stage II | 0.083786752 | 0.487249 |
| HOXB9 | Stage III | -0.192129774 | 0.200833 |
| HOXB9 | Stage IV | 0.058159232 | 0.797113 |
| HOXB9 | overall | -0.073131776 | 0.199826 |
| HPGD | G1 | -0.105263158 | 0.658728 |
| HPGD | G2 | 0.046701573 | 0.589268 |
| HPGD | G3 | -0.059525112 | 0.518401 |
| HPGD | GX | 0.043478261 | 0.840133 |
| HPGD | Stage I | 0.01739928 | 0.825524 |
| HPGD | Stage II | -0.137040049 | 0.254449 |
| HPGD | Stage III | -0.089496084 | 0.554201 |
| HPGD | Stage IV | 0.077357425 | 0.732221 |
| HPGD | overall | -0.020772045 | 0.716083 |
| HPSE | G1 | 0.040601504 | 0.865047 |
| HPSE | G2 | 0.091788223 | 0.287877 |
| HPSE | G3 | 0.071964232 | 0.434749 |
| HPSE | GX | 0.16 | 0.45517 |
| HPSE | Stage I | 0.096969328 | 0.218174 |
| HPSE | Stage II | 0.131121153 | 0.275735 |
| HPSE | Stage III | 0.235181646 | 0.11564 |
| HPSE | Stage IV | -0.04121965 | 0.855482 |
| HPSE | overall | 0.085809071 | 0.132311 |
| HRAS | G1 | 0.276691729 | 0.237619 |
| HRAS | G2 | 0.148541918 | 0.084369 |
| HRAS | G3 | 0.116598516 | 0.204714 |
| PIK3CA | Stage II | 0.244301176 | 0.040053 |
| FOSL1 | overall | 0.243826617 | 1.46E-05 |
| PRDX1 | G2 | 0.243047124 | 0.004357 |
| HRAS | Stage III | -0.036945661 | 0.807413 |
| HRAS | Stage IV | -0.014116318 | 0.950285 |
| FSCN1 | overall | 0.242594192 | 1.62E-05 |
| HS3ST3B1 | G1 | 0.021052632 | 0.929799 |
| HS3ST3B1 | G2 | 0.026031988 | 0.763543 |

| | | | |
|---|---|---|---|
| HS3ST3B1 | G3 | 0.054937709 | 0.5512 |
| THBD | Stage II | 0.241400414 | 0.042553 |
| EIF5A2 | overall | 0.238883904 | 2.20E-05 |
| HS3ST3B1 | Stage II | 0.008886728 | 0.941366 |
| HS3ST3B1 | Stage III | -0.171282306 | 0.255054 |
| HS3ST3B1 | Stage IV | -0.08865048 | 0.694829 |
| HS3ST3B1 | overall | 0.06504089 | 0.254333 |
| HS6ST2 | G1 | 0.263157895 | 0.262287 |
| HS6ST2 | G2 | 0.077406581 | 0.370402 |
| HS6ST2 | G3 | 0.067373356 | 0.464691 |
| HS6ST2 | GX | 0.228695652 | 0.282423 |
| BMP2 | Stage II | 0.238399048 | 0.045274 |
| HS6ST2 | Stage II | 0.093176502 | 0.439601 |
| HS6ST2 | Stage III | -0.028804046 | 0.849292 |
| HS6ST2 | Stage IV | -0.256916996 | 0.248414 |
| HS6ST2 | overall | 0.070099125 | 0.219166 |
| HSP90AA1 | G1 | 0.121804511 | 0.608957 |
| HSP90AA1 | G2 | -0.042541417 | 0.622891 |
| HSP90AA1 | G3 | 0.067658115 | 0.462801 |
| HSP90AA1 | GX | -0.010434783 | 0.961404 |
| HSP90AA1 | Stage I | -0.03220765 | 0.68317 |
| HSP90AA1 | Stage II | -0.032344336 | 0.788876 |
| HSP90AA1 | Stage III | -0.001788688 | 0.990587 |
| HSP90AA1 | Stage IV | 0.273856578 | 0.217476 |
| HSP90AA1 | overall | -0.016415648 | 0.773798 |
| HSPA4 | G1 | -0.323308271 | 0.16439 |
| HSPA4 | G2 | 0.002242286 | 0.979331 |
| HSPA4 | G3 | -0.052656162 | 0.567879 |
| HSPA4 | GX | -0.016521739 | 0.938923 |
| HSPA4 | Stage I | -0.088841905 | 0.259421 |
| HSPA4 | Stage II | 0.046848146 | 0.698049 |
| HSPA4 | Stage III | -0.033861716 | 0.82322 |
| HSPA4 | Stage IV | 0.016374929 | 0.942342 |
| HSPA4 | overall | -0.052123415 | 0.361162 |
| HSPB1 | G1 | 0.079699248 | 0.738374 |
| CAV1 | overall | 0.238218679 | 2.32E-05 |
| VSNL1 | Stage I | 0.235102265 | 0.00252 |
| HSPB1 | GX | 0.132173913 | 0.538118 |
| BMP7 | Stage I | 0.234528662 | 0.002584 |
| ITGB4 | Stage II | 0.233955684 | 0.049565 |
| HSPB1 | Stage III | 0.228396967 | 0.126838 |
| HSPB1 | Stage IV | 0.054771316 | 0.808714 |
| IL1B | Stage I | 0.232342322 | 0.002841 |
| ID1 | G1 | 0.154887218 | 0.514383 |

| | | | |
|---|---|---|---|
| ID1 | G2 | 0.105151295 | 0.2231 |
| ID1 | G3 | 0.107048661 | 0.244532 |
| PTPRZ1 | Stage I | 0.231472655 | 0.002949 |
| ID1 | Stage I | 0.144231968 | 0.066228 |
| SDC1 | Stage I | 0.230183691 | 0.003117 |
| ID1 | Stage III | 0.172762599 | 0.250909 |
| ID1 | Stage IV | -0.34839074 | 0.11207 |
| BMP7 | overall | 0.226342867 | 5.95E-05 |
| ID2 | G1 | 0.031578947 | 0.894854 |
| EIF5A2 | G2 | 0.225607385 | 0.008269 |
| CDH13 | Stage I | 0.224694286 | 0.003932 |
| IL18 | G2 | 0.223911358 | 0.00878 |
| CLDN1 | G2 | 0.223713369 | 0.008842 |
| MET | G3 | 0.221514955 | 0.015038 |
| ID2 | Stage III | -0.215074325 | 0.151167 |
| ID2 | Stage IV | -0.1959345 | 0.382181 |
| TGFB1 | G2 | 0.220829407 | 0.00978 |
| IDH1 | G1 | 0.106766917 | 0.654142 |
| IDH1 | G2 | -0.13122622 | 0.127805 |
| IDH1 | G3 | 0.039341928 | 0.669654 |
| IDH1 | GX | -0.288695652 | 0.171264 |
| IDH1 | Stage I | -0.08087243 | 0.304776 |
| IDH1 | Stage II | -0.07246875 | 0.548117 |
| IDH1 | Stage III | 0.146857461 | 0.330097 |
| IDH1 | Stage IV | -0.042348955 | 0.851564 |
| IDH1 | overall | -0.062334031 | 0.274678 |
| IDH2 | G1 | 0.156390977 | 0.510257 |
| IDH2 | G2 | -0.128704841 | 0.135357 |
| IDH2 | G3 | -0.093269089 | 0.310951 |
| IDH2 | GX | -0.402608696 | 0.051113 |
| IDH2 | Stage I | -0.106185764 | 0.177313 |
| IDH2 | Stage II | -0.196463753 | 0.10057 |
| IDH2 | Stage III | -0.005736138 | 0.96982 |
| IDH2 | Stage IV | -0.387916431 | 0.074441 |
| PHLDA1 | Stage I | 0.219667643 | 0.004841 |
| IGF1R | G1 | 0.285714286 | 0.222035 |
| IGF1R | G2 | 0.153090896 | 0.075181 |
| VSNL1 | G3 | 0.219108391 | 0.0162 |
| IGF1R | GX | 0.31826087 | 0.129609 |
| BIRC2 | Stage I | 0.218307069 | 0.005117 |
| IGF1R | Stage II | 0.221816078 | 0.063011 |
| IGF1R | Stage III | 0.216184545 | 0.149017 |
| IGF1R | Stage IV | 0.219649915 | 0.326013 |
| PTPN14 | Stage I | 0.217725153 | 0.00524 |

| | | | |
|---|---|---|---|
| IGFBP3 | G1 | 0.285714286 | 0.222035 |
| IGFBP3 | G2 | -0.030719321 | 0.722572 |
| IGFBP3 | G3 | 0.132670053 | 0.148596 |
| IGFBP3 | GX | -0.133043478 | 0.535424 |
| IGFBP3 | Stage I | 0.100369378 | 0.202383 |
| IGFBP3 | Stage II | 0.049648304 | 0.680946 |
| IGFBP3 | Stage III | -0.050638377 | 0.738222 |
| IGFBP3 | Stage IV | -0.016374929 | 0.942342 |
| IGFBP3 | overall | 0.057491783 | 0.313766 |
| IL18 | G1 | 0.163909774 | 0.489867 |
| SDC1 | G2 | 0.217382489 | 0.011017 |
| THBD | overall | 0.215030391 | 1.39E-04 |
| CD44 | Stage I | 0.214474738 | 0.005974 |
| TP73 | G2 | 0.213475186 | 0.012582 |
| LIMA1 | Stage I | 0.212255145 | 0.006526 |
| IL18 | Stage III | 0.203108618 | 0.175801 |
| IL18 | Stage IV | 0.18351214 | 0.413653 |
| PTPN14 | overall | 0.211346541 | 1.82E-04 |
| IL1B | G1 | -0.168421053 | 0.477829 |
| IL1B | G2 | 0.057495557 | 0.506134 |
| IL1B | G3 | 0.119237748 | 0.194581 |
| JAG1 | G3 | 0.210603812 | 0.020951 |
| HBEGF | G2 | 0.210443329 | 0.013928 |
| IL1B | Stage II | 0.204679784 | 0.086855 |
| IL1B | Stage III | -0.197187444 | 0.189006 |
| IL1B | Stage IV | -0.314511575 | 0.153997 |
| IL1B | overall | 0.107873738 | 0.058213 |
| IL6R | G1 | -0.190977444 | 0.419927 |
| IL6R | G2 | 0.120048185 | 0.16389 |
| PDPN | G3 | 0.210398924 | 0.021079 |
| IL6R | GX | 0.275652174 | 0.192317 |
| IL6R | Stage I | -0.013336954 | 0.865819 |
| IL6R | Stage II | 0.167539969 | 0.162548 |
| IL6R | Stage III | -0.084068341 | 0.578572 |
| PDPN | overall | 0.210212954 | 1.98E-04 |
| IL6R | overall | -0.014903793 | 0.794141 |
| ILK | G1 | -0.118796992 | 0.617892 |
| ILK | G2 | -0.144973343 | 0.092188 |
| ILK | G3 | -0.082073187 | 0.372843 |
| ILK | GX | -0.31826087 | 0.129609 |
| ILK | Stage I | -0.060211651 | 0.445161 |
| ILK | Stage II | -0.189706486 | 0.113069 |
| ILK | Stage III | -0.138962562 | 0.357034 |
| ILK | Stage IV | -0.225296443 | 0.313412 |

| | | | |
|---|---|---|---|
| MMP13 | overall | 0.210013875 | 2.01E-04 |
| ITGA5 | G1 | 0.009022556 | 0.969885 |
| ITGA5 | G2 | 0.044831411 | 0.604281 |
| ITGA5 | G3 | 0.030479663 | 0.741045 |
| VSNL1 | overall | 0.20877838 | 2.19E-04 |
| ITGA5 | Stage I | 0.118854901 | 0.130758 |
| ITGA5 | Stage II | 0.211956841 | 0.07598 |
| ITGA5 | Stage III | -0.103312158 | 0.494452 |
| ITGA5 | Stage IV | -0.112365895 | 0.618583 |
| ITGA5 | overall | 0.078322398 | 0.169655 |
| ITGA6 | G1 | -0.243609023 | 0.300668 |
| BMP7 | G3 | 0.208416027 | 0.022352 |
| JAG2 | G3 | 0.207707601 | 0.022822 |
| LCN2 | G2 | 0.207036962 | 0.015588 |
| IGF1R | overall | 0.205395721 | 2.78E-04 |
| SDC1 | overall | 0.204819166 | 2.90E-04 |
| ITGA6 | Stage III | 0.087028928 | 0.565219 |
| SPRR2A | overall | 0.204330505 | 3.00E-04 |
| EGFR | G3 | 0.203651518 | 0.025683 |
| ITGB1 | G1 | 0.022556391 | 0.924798 |
| ITGB1 | G2 | 0.018167289 | 0.833725 |
| ITGB1 | G3 | -0.091463298 | 0.320453 |
| ITGB1 | GX | -0.027826087 | 0.897305 |
| ITGB1 | Stage I | 0.015675701 | 0.842571 |
| ITGB1 | Stage II | 0.064353323 | 0.59391 |
| ITGB1 | Stage III | -0.144637021 | 0.337541 |
| ITGB1 | Stage IV | -0.029926595 | 0.894823 |
| ITGB1 | overall | -0.024749969 | 0.664748 |
| ITGB4 | G1 | -0.02556391 | 0.914804 |
| ITGB4 | G2 | 0.15758501 | 0.066915 |
| ITGB4 | G3 | 0.08194817 | 0.373574 |
| CDH1 | G3 | 0.202818076 | 0.026307 |
| EGFR | Stage I | 0.202379204 | 0.009576 |
| NOTCH1 | Stage I | 0.20099369 | 0.010092 |
| ITGB4 | Stage III | -0.044470487 | 0.769172 |
| ITGB4 | Stage IV | -0.128176172 | 0.569723 |
| LYPD3 | G2 | 0.200469926 | 0.019279 |
| JAG1 | G1 | 0.127819549 | 0.591246 |
| ITGA6 | overall | 0.200459312 | 3.92E-04 |
| HSPB1 | overall | 0.200053588 | 4.03E-04 |
| JAG1 | GX | 0.223478261 | 0.293847 |
| MET | Stage I | 0.19821435 | 0.011201 |
| JAG1 | Stage II | 0.177734555 | 0.138109 |
| JAG1 | Stage III | 0.191389627 | 0.202606 |

| | | | |
|---|---|---|---|
| JAG1 | Stage IV | -0.184641446 | 0.410735 |
| CD44 | overall | 0.195633944 | 5.43E-04 |
| JAG2 | G1 | 0.090225564 | 0.705213 |
| PRKCE | Stage I | 0.19562344 | 0.01233 |
| KRT19 | G3 | 0.194223207 | 0.033535 |
| MUC4 | Stage I | 0.194085519 | 0.013046 |
| JAG2 | G2 | 0.193952984 | 0.023666 |
| MMP13 | G3 | 0.192498624 | 0.035171 |
| JAG2 | Stage III | -0.039166102 | 0.796078 |
| JAG2 | Stage IV | 0.383399209 | 0.078178 |
| TGFA | Stage I | 0.192217847 | 0.013965 |
| JAK2 | G1 | 0.210526316 | 0.37297 |
| JAK2 | G2 | -0.086413893 | 0.317155 |
| JAK2 | G3 | 0.068463776 | 0.457478 |
| JAK2 | GX | 0.047826087 | 0.824378 |
| JAK2 | Stage I | 0.062661239 | 0.426829 |
| JAK2 | Stage II | 0.06774034 | 0.57459 |
| JAK2 | Stage III | 0.002775551 | 0.985394 |
| JAK2 | Stage IV | -0.080745342 | 0.720938 |
| JAK2 | overall | 0.023863274 | 0.676066 |
| KHDRBS1 | G1 | 0.168421053 | 0.477829 |
| KHDRBS1 | G2 | -0.058545138 | 0.498389 |
| KHDRBS1 | G3 | -0.081694665 | 0.375059 |
| ACVR1 | G3 | 0.191976386 | 0.035679 |
| KHDRBS1 | Stage I | -0.047717089 | 0.545267 |
| KHDRBS1 | Stage II | -0.062458605 | 0.604842 |
| KHDRBS1 | Stage III | -0.166224636 | 0.269564 |
| KHDRBS1 | Stage IV | 0.083003953 | 0.713446 |
| KHDRBS1 | overall | -0.074411993 | 0.192046 |
| KIT | G1 | 0.094736842 | 0.691153 |
| KIT | G2 | -0.040055819 | 0.643363 |
| KIT | G3 | -0.005354864 | 0.953712 |
| KIT | GX | -0.057391304 | 0.789955 |
| KIT | Stage I | 0.085477878 | 0.277967 |
| ITGB4 | Stage I | 0.190774142 | 0.014714 |
| KIT | Stage III | -0.162277186 | 0.281262 |
| KIT | Stage IV | 0.033314512 | 0.882991 |
| KIT | overall | -0.025434717 | 0.656058 |
| KLF4 | G1 | -0.139849624 | 0.556494 |
| KLF4 | G2 | 0.118853094 | 0.168157 |
| KLF4 | G3 | 0.0663003 | 0.471849 |
| KLF4 | GX | -0.066086957 | 0.758987 |
| KLF4 | Stage I | 0.115718098 | 0.1413 |
| KLF4 | Stage II | -0.107194058 | 0.37359 |

| | | | |
|---|---|---|---|
| KLF4 | Stage III | 0.20298526 | 0.176069 |
| KLF4 | Stage IV | -0.086391869 | 0.702256 |
| KLF4 | overall | 0.075167717 | 0.187558 |
| KLF6 | G1 | 0.187969925 | 0.427421 |
| KLF6 | G2 | 0.049986284 | 0.563321 |
| KLF6 | G3 | 0.0185163 | 0.84091 |
| KLF6 | GX | -0.05826087 | 0.786844 |
| SDC1 | G3 | 0.190503972 | 0.037147 |
| KLF6 | Stage II | 0.096077264 | 0.425429 |
| KLF6 | Stage III | -0.161537039 | 0.283491 |
| KLF6 | Stage IV | -0.093167702 | 0.680057 |
| KLF6 | overall | 0.053124608 | 0.351999 |
| KLF8 | G1 | 0.058646617 | 0.805993 |
| KLF8 | G2 | 0.090657539 | 0.293882 |
| KLF8 | G3 | 0.147470591 | 0.107977 |
| KLF8 | GX | -0.266086957 | 0.208836 |
| KLF8 | Stage I | 0.145140865 | 0.064519 |
| KLF8 | Stage II | 0.106506594 | 0.376681 |
| KLF8 | Stage III | 0.089372726 | 0.55475 |
| KLF8 | Stage IV | -0.333709768 | 0.129073 |
| KLF8 | overall | 0.077407638 | 0.17471 |
| KLK6 | G1 | 0.118796992 | 0.617892 |
| KLK6 | G2 | 0.143858687 | 0.094745 |
| KLK6 | G3 | -7.05E-04 | 0.993903 |
| KLK6 | GX | 0.01826087 | 0.932507 |
| KLK6 | Stage I | 0.113235336 | 0.150094 |
| KLK6 | Stage II | 0.020944252 | 0.862364 |
| KLK6 | Stage III | -0.001912105 | 0.989938 |
| KLK6 | Stage IV | 0.002823264 | 0.990051 |
| KLK6 | overall | 0.06172282 | 0.27942 |
| KRAS | G1 | 0.237593985 | 0.313131 |
| KRAS | G2 | -0.109910189 | 0.202736 |
| KRAS | G3 | -0.160805661 | 0.079339 |
| KRAS | GX | -0.302608696 | 0.150636 |
| VSNL1 | G2 | 0.190226972 | 0.026539 |
| KRAS | Stage II | 0.017253666 | 0.886439 |
| KRAS | Stage III | 0.037685808 | 0.80363 |
| KRAS | Stage IV | 0.052512705 | 0.816469 |
| SNAI2 | G2 | 0.190179264 | 0.026578 |
| RAC1 | overall | 0.189931846 | 7.91E-04 |
| KRT19 | G2 | -0.016931646 | 0.844887 |
| HS3ST3B1 | Stage I | 0.189917894 | 0.015174 |
| KRT19 | GX | -0.297391304 | 0.158154 |
| PDPN | G2 | 0.189790442 | 0.026895 |

| | | | |
|---|---|---|---|
| KRT19 | Stage II | 0.079544597 | 0.509637 |
| KRT19 | Stage III | -0.058779992 | 0.697994 |
| MYC | Stage I | 0.18931104 | 0.015508 |
| KRT19 | overall | 0.09574656 | 0.092936 |
| L1CAM | G1 | 0.233082707 | 0.322678 |
| L1CAM | G2 | -0.085402478 | 0.322874 |
| L1CAM | G3 | -0.036615879 | 0.691337 |
| L1CAM | GX | 0.273043478 | 0.196731 |
| L1CAM | Stage I | 0.043663076 | 0.579971 |
| L1CAM | Stage II | 0.043746175 | 0.717171 |
| L1CAM | Stage III | -0.27255906 | 0.066865 |
| L1CAM | Stage IV | -0.086391869 | 0.702256 |
| L1CAM | overall | -0.022064463 | 0.69925 |
| LAMA5 | G1 | 0.329323308 | 0.156231 |
| LAMA5 | G2 | 0.004990279 | 0.95402 |
| LAMA5 | G3 | 0.104131615 | 0.2577 |
| LAMA5 | GX | -0.056521739 | 0.79307 |
| LAMA5 | Stage I | 0.101741036 | 0.196253 |
| LAMA5 | Stage II | -0.030919106 | 0.797977 |
| LAMA5 | Stage III | -0.076420157 | 0.613712 |
| LAMA5 | Stage IV | 0.21852061 | 0.328569 |
| LAMA5 | overall | 0.051839714 | 0.363785 |
| LATS1 | G1 | 0.15037594 | 0.526855 |
| LATS1 | G2 | -0.154519757 | 0.072468 |
| LATS1 | G3 | -0.145272388 | 0.113391 |
| JAG2 | overall | 0.188784023 | 8.53E-04 |
| PKP3 | G2 | 0.188609663 | 0.027876 |
| LATS1 | Stage II | -0.175571559 | 0.143048 |
| LATS1 | Stage III | -0.116018011 | 0.442597 |
| LATS1 | Stage IV | -0.129305477 | 0.566299 |
| JAG1 | overall | 0.18748225 | 9.27E-04 |
| LCN2 | G1 | -0.326315789 | 0.160275 |
| CTNND1 | Stage I | 0.186315559 | 0.017252 |
| LCN2 | G3 | 0.107944611 | 0.240583 |
| LCN2 | GX | 0.193043478 | 0.366116 |
| CD44 | G3 | 0.185933933 | 0.042027 |
| LCN2 | Stage II | 0.043176083 | 0.720705 |
| LCN2 | Stage III | 0.26861161 | 0.071073 |
| LCN2 | Stage IV | -0.184641446 | 0.410735 |
| EGFR | overall | 0.185828239 | 0.001031 |
| LEF1 | G1 | 0.087218045 | 0.714638 |
| LEF1 | G2 | -0.004141073 | 0.961838 |
| LEF1 | G3 | 0.01273777 | 0.890177 |
| LEF1 | GX | -0.124347826 | 0.562649 |

| | | | |
|---|---|---|---|
| LEF1 | Stage I | -0.022132195 | 0.779152 |
| LEF1 | Stage II | -0.032092824 | 0.79048 |
| LEF1 | Stage III | 0.006723 | 0.964631 |
| LEF1 | Stage IV | 0.419536985 | 0.051932 |
| LEF1 | overall | 0.007245882 | 0.899054 |
| LEFTY1 | G1 | -0.079699248 | 0.738374 |
| LEFTY1 | G2 | -0.043004301 | 0.619109 |
| LEFTY1 | G3 | 0.069124064 | 0.453141 |
| LEFTY1 | GX | -0.279626012 | 0.185724 |
| LEFTY1 | Stage I | -0.014604163 | 0.853207 |
| JAG1 | G2 | 0.185475233 | 0.030632 |
| LEFTY1 | Stage III | 0.046754048 | 0.757669 |
| LEFTY1 | Stage IV | 0.124788255 | 0.580049 |
| LEFTY1 | overall | -0.043049448 | 0.450835 |
| LETMD1 | G1 | -0.030075188 | 0.899837 |
| LETMD1 | G2 | -0.119225219 | 0.16682 |
| LETMD1 | G3 | 0.027052134 | 0.769298 |
| LETMD1 | GX | 0.336521739 | 0.107856 |
| LETMD1 | Stage I | 0.048407075 | 0.539464 |
| LETMD1 | Stage II | -0.20464625 | 0.086907 |
| LETMD1 | Stage III | -0.083328194 | 0.581932 |
| LETMD1 | Stage IV | 0.254658385 | 0.252743 |
| LETMD1 | overall | -0.012862768 | 0.821824 |
| LIMA1 | G1 | 0.009022556 | 0.969885 |
| LIMA1 | G2 | 0.037532054 | 0.664428 |
| LIMA1 | G3 | 0.114525329 | 0.212936 |
| LIMA1 | GX | 0.32173913 | 0.125239 |
| HSPB1 | G3 | 0.185062291 | 0.043016 |
| LCN2 | Stage I | 0.184264999 | 0.018541 |
| LIMA1 | Stage III | -0.242706472 | 0.104116 |
| LIMA1 | Stage IV | -0.323546019 | 0.141879 |
| LIMA1 | overall | 0.107401715 | 0.059328 |
| LIMS1 | G1 | 0.198496241 | 0.401504 |
| LIMS1 | G2 | -0.042016626 | 0.62719 |
| LIMS1 | G3 | -0.030767895 | 0.738684 |
| LIMS1 | GX | -0.213043478 | 0.317545 |
| LIMS1 | Stage I | -0.03622841 | 0.646145 |
| LIMS1 | Stage II | 0.137090351 | 0.254273 |
| LIMS1 | Stage III | -0.139702709 | 0.354453 |
| LIMS1 | Stage IV | -0.181253529 | 0.419523 |
| LIMS1 | overall | -0.034322001 | 0.547802 |
| LOX | G1 | 0.082706767 | 0.728851 |
| LOX | G2 | -0.09908281 | 0.251111 |
| LOX | G3 | 0.14691149 | 0.109335 |

| | | | |
|---|---|---|---|
| LOX | GX | -0.391304348 | 0.058644 |
| LOX | Stage I | 0.020325485 | 0.796771 |
| LOX | Stage II | 0.092405201 | 0.443414 |
| LOX | Stage III | -0.188922471 | 0.208598 |
| LOX | Stage IV | -0.115753811 | 0.607971 |
| LOX | overall | -0.019961817 | 0.726707 |
| LOXL2 | G1 | -0.010526316 | 0.964869 |
| LOXL2 | G2 | -0.119241916 | 0.16676 |
| LOXL2 | G3 | -0.051555324 | 0.576011 |
| LOXL2 | GX | 0.010434783 | 0.961404 |
| LOXL2 | Stage I | 0.019189364 | 0.807904 |
| LOXL2 | Stage II | -0.083602311 | 0.488211 |
| CTNND1 | G2 | 0.184110778 | 0.031902 |
| LOXL2 | Stage IV | 0.1134952 | 0.615037 |
| LOXL2 | overall | -0.064695974 | 0.256866 |
| LOXL3 | G1 | 0.112781955 | 0.635917 |
| PTHLH | G3 | 0.184082997 | 0.044151 |
| LOXL3 | G3 | -0.003347658 | 0.971053 |
| LOXL3 | GX | -0.242608696 | 0.253348 |
| LOXL3 | Stage I | -0.005616872 | 0.94327 |
| MMP3 | Stage I | 0.18272985 | 0.01956 |
| LOXL3 | Stage III | -0.126256709 | 0.403098 |
| LOXL3 | Stage IV | 5.65E-04 | 0.99801 |
| LOXL3 | overall | -0.099693572 | 0.080168 |
| LRP6 | G1 | 0.07518797 | 0.752727 |
| LRP6 | G2 | -0.159335902 | 0.063902 |
| LRP6 | G3 | -0.082576725 | 0.369907 |
| AXL | Stage I | 0.181926252 | 0.020113 |
| MARVELD3 | G3 | 0.181742415 | 0.046964 |
| LRP6 | Stage II | -0.195910428 | 0.101551 |
| LRP6 | Stage III | 0.141676434 | 0.347628 |
| LRP6 | Stage IV | 0.180124224 | 0.422475 |
| HAS2 | Stage I | 0.180917598 | 0.020826 |
| LYPD3 | G1 | -0.010526316 | 0.964869 |
| PARD3 | G2 | 0.180439631 | 0.035542 |
| LYPD3 | G3 | 0.113671051 | 0.216391 |
| LYPD3 | GX | 0.374782609 | 0.071159 |
| AXL | G3 | 0.180294309 | 0.048777 |
| ITGA6 | G2 | 0.179700154 | 0.036316 |
| KRT19 | Stage I | 0.17903607 | 0.022214 |
| LYPD3 | Stage IV | -0.224167137 | 0.315909 |
| HSPB1 | Stage I | 0.177991393 | 0.023018 |
| MAGED1 | G1 | -0.138345865 | 0.560787 |
| MAGED1 | G2 | -0.132678936 | 0.123604 |

| | | | |
|---|---|---|---|
| MAGED1 | G3 | 0.027639016 | 0.764438 |
| MAGED1 | GX | -0.382608696 | 0.065 |
| MAGED1 | Stage I | 0.013062622 | 0.868554 |
| MAGED1 | Stage II | -0.149649142 | 0.212907 |
| CDH13 | overall | 0.176652371 | 0.001826 |
| MAGED1 | Stage IV | 0.157538114 | 0.483816 |
| MAGED1 | overall | -0.069237031 | 0.2249 |
| MAP2K1 | G1 | -0.230075188 | 0.329138 |
| MAP2K1 | G2 | 0.133704662 | 0.120702 |
| MAP2K1 | G3 | 0.10215219 | 0.266906 |
| MAP2K1 | GX | -8.70E-04 | 0.996782 |
| MAP2K1 | Stage I | 0.121722914 | 0.121657 |
| MAP2K1 | Stage II | 0.086503073 | 0.473191 |
| MAP2K1 | Stage III | 0.251464876 | 0.091836 |
| MAP2K1 | Stage IV | -0.363071711 | 0.096757 |
| MAP2K1 | overall | 0.074773382 | 0.18989 |
| MAP3K3 | G1 | -0.082706767 | 0.728851 |
| MAP3K3 | G2 | -0.0501962 | 0.561681 |
| MAP3K3 | G3 | -0.115105266 | 0.210613 |
| MAP3K3 | GX | -0.280869565 | 0.183694 |
| MAP3K3 | Stage I | -0.049961621 | 0.526502 |
| MAP3K3 | Stage II | -0.108703125 | 0.366861 |
| MAP3K3 | Stage III | -0.123789553 | 0.412424 |
| MAP3K3 | Stage IV | 0.010728402 | 0.962207 |
| MAP3K3 | overall | -0.077670393 | 0.173247 |
| MAP3K4 | G1 | 0.419548872 | 0.065548 |
| MAP3K4 | G2 | -0.077335019 | 0.370846 |
| MAP3K4 | G3 | -0.079868039 | 0.385866 |
| MAP3K4 | GX | -0.136521739 | 0.52471 |
| MAP3K4 | Stage I | -0.034316401 | 0.66365 |
| MAP3K4 | Stage II | -0.18346901 | 0.125639 |
| MAP3K4 | Stage III | 0.014741257 | 0.922541 |
| MAP3K4 | Stage IV | -0.102202146 | 0.650853 |
| MAP3K4 | overall | -0.075556561 | 0.185279 |
| MAP3K7 | G1 | 0.273684211 | 0.242966 |
| MAP3K7 | G2 | -0.14129981 | 0.100827 |
| MMP14 | overall | 0.175374596 | 0.001973 |
| MAP3K7 | GX | -0.251304348 | 0.236199 |
| HBEGF | Stage I | 0.172186091 | 0.02796 |
| MAP3K7 | Stage II | -0.032746753 | 0.786311 |
| MAP3K7 | Stage III | -0.131684452 | 0.383014 |
| MAP3K7 | Stage IV | 0.107848673 | 0.632846 |
| CAV1 | G2 | 0.1718116 | 0.045494 |
| MAPK1 | G1 | 0.067669173 | 0.776821 |

| | | | |
|---|---|---|---|
| MAPK1 | G2 | -2.46E-04 | 0.997735 |
| MAPK1 | G3 | 0.032841082 | 0.721776 |
| SEMA7A | Stage I | 0.171216232 | 0.028868 |
| MAPK1 | Stage I | -0.00520676 | 0.947406 |
| MAPK1 | Stage II | 0.014403206 | 0.905105 |
| MAPK1 | Stage III | -0.021155863 | 0.889013 |
| MAPK1 | Stage IV | -0.334839074 | 0.127703 |
| MAPK1 | overall | -0.028380943 | 0.61921 |
| MAPK14 | G1 | 0.030075188 | 0.899837 |
| MAPK14 | G2 | 0.001493267 | 0.986234 |
| MAPK14 | G3 | -0.080611191 | 0.381447 |
| MAPK14 | GX | 0.035652174 | 0.868639 |
| MAPK14 | Stage I | -0.009859315 | 0.900595 |
| MAPK14 | Stage II | -0.002934297 | 0.980624 |
| MAPK14 | Stage III | -0.046814285 | 0.757366 |
| MAPK14 | Stage IV | -0.192546584 | 0.390627 |
| MAPK14 | overall | -0.03731785 | 0.513397 |
| MAPK3 | G1 | -0.036090226 | 0.879931 |
| MAPK3 | G2 | -0.027048173 | 0.754601 |
| MAPK3 | G3 | -0.102520294 | 0.265177 |
| MAPK3 | GX | -0.220869565 | 0.299665 |
| MAPK3 | Stage I | -0.069876994 | 0.37543 |
| PKP3 | Stage I | 0.170570582 | 0.029487 |
| MAPK3 | Stage III | 0.212977242 | 0.15529 |
| MAPK3 | Stage IV | -0.084133258 | 0.709709 |
| MAPK3 | overall | -0.058804945 | 0.302823 |
| MAPK7 | G1 | -0.178947368 | 0.450325 |
| MAPK7 | G2 | -0.021463927 | 0.804112 |
| MAPK7 | G3 | -0.023430134 | 0.799486 |
| MAPK7 | GX | -0.288695652 | 0.171264 |
| MAPK7 | Stage I | 0.00428955 | 0.956661 |
| MAPK7 | Stage II | -0.077867856 | 0.518628 |
| MAPK7 | Stage III | -0.278850307 | 0.06057 |
| MAPK7 | Stage IV | -0.020892151 | 0.926474 |
| MAPK7 | overall | -0.057945095 | 0.30996 |
| MAPK8 | G1 | -0.007518797 | 0.974903 |
| MAPK8 | G2 | 0.007430555 | 0.931581 |
| MAPK8 | G3 | -0.097752312 | 0.288163 |
| MAPK8 | GX | -0.204347826 | 0.338157 |
| MAPK8 | Stage I | -0.033493406 | 0.671242 |
| MAPK8 | Stage II | -0.019601103 | 0.871112 |
| MAPK8 | Stage III | 0.022389441 | 0.882586 |
| MAPK8 | Stage IV | 0.059288538 | 0.793255 |
| MAPK8 | overall | -0.023790061 | 0.677004 |

| | | | |
|---|---|---|---|
| MARVELD3 | G1 | -0.144360902 | 0.543701 |
| MARVELD3 | G2 | 0.032890044 | 0.703854 |
| HRAS | overall | 0.169988535 | 0.002719 |
| MARVELD3 | GX | -0.233913043 | 0.271284 |
| ANXA1 | overall | 0.169915118 | 0.002731 |
| MARVELD3 | Stage II | 0.072183704 | 0.549696 |
| MARVELD3 | Stage III | -0.21088016 | 0.159494 |
| MARVELD3 | Stage IV | -0.21852061 | 0.328569 |
| MARVELD3 | overall | 0.047856502 | 0.401856 |
| MCAM | G1 | -0.093233083 | 0.695829 |
| MCAM | G2 | -0.064744821 | 0.453939 |
| MCAM | G3 | -0.016654946 | 0.856722 |
| MCAM | GX | 0.066086957 | 0.758987 |
| MCAM | Stage I | 0.06660164 | 0.398274 |
| MCAM | Stage II | -0.136587329 | 0.256036 |
| YAP1 | Stage I | 0.169373498 | 0.030663 |
| MCAM | Stage IV | 0.017504235 | 0.938373 |
| MCAM | overall | -0.045623699 | 0.424199 |
| MDK | G1 | 0.028571429 | 0.904822 |
| MDK | G2 | -5.77E-04 | 0.994678 |
| MDK | G3 | 0.076617615 | 0.405554 |
| MDK | GX | -0.286086957 | 0.17534 |
| MDK | Stage I | 0.04442788 | 0.573346 |
| MDK | Stage II | 0.100973348 | 0.40211 |
| MDK | Stage III | -0.276383151 | 0.06298 |
| MDK | Stage IV | 0.184641446 | 0.410735 |
| MDK | overall | 0.021291249 | 0.709304 |
| MET | G1 | -0.013533835 | 0.95484 |
| MET | G2 | 0.013451332 | 0.876485 |
| TGFB1 | overall | 0.167751054 | 0.003098 |
| MET | GX | 0.29826087 | 0.156883 |
| TMPRSS4 | Stage I | 0.16678813 | 0.033341 |
| MET | Stage II | 0.102549485 | 0.394766 |
| MET | Stage III | 0.008820083 | 0.953609 |
| MET | Stage IV | -0.121400339 | 0.590455 |
| MET | overall | 0.099191247 | 0.08171 |
| MGAT3 | G1 | 0.027067669 | 0.909812 |
| MGAT3 | G2 | -0.085326145 | 0.323308 |
| MGAT3 | G3 | 0.050572557 | 0.583316 |
| MGAT3 | GX | -0.273043478 | 0.196731 |
| MGAT3 | Stage I | -0.107945366 | 0.170201 |
| MGAT3 | Stage II | -0.10018528 | 0.405812 |
| MGAT3 | Stage III | 0.20409548 | 0.173666 |
| MGAT3 | Stage IV | 0.259175607 | 0.244133 |

| | | | |
|---|---|---|---|
| MGAT3 | overall | -0.019520504 | 0.732516 |
| MMP13 | G1 | 0.181954887 | 0.442621 |
| MMP13 | G2 | 0.161002303 | 0.061138 |
| LYPD3 | Stage I | 0.166322597 | 0.033844 |
| TMPRSS4 | overall | 0.166239809 | 0.003381 |
| NOTCH1 | overall | 0.166197101 | 0.003389 |
| LYPD3 | overall | 0.165425311 | 0.003543 |
| MMP13 | Stage III | -0.20076482 | 0.180946 |
| MMP13 | Stage IV | -0.378881988 | 0.082054 |
| HS6ST2 | Stage I | 0.16503407 | 0.035269 |
| MMP14 | G1 | 0.166917293 | 0.481825 |
| MMP14 | G2 | 0.100473504 | 0.244487 |
| MMP14 | G3 | 0.157381604 | 0.086034 |
| HBEGF | overall | 0.163682427 | 0.003913 |
| PKP3 | overall | 0.163474989 | 0.003959 |
| MMP14 | Stage II | 0.189874161 | 0.112745 |
| MMP14 | Stage III | -0.194473572 | 0.195289 |
| MMP14 | Stage IV | -0.130434783 | 0.562884 |
| TGFB2 | Stage I | 0.161595225 | 0.039322 |
| MMP2 | G1 | -0.093233083 | 0.695829 |
| MMP2 | G2 | 0.008453896 | 0.922186 |
| MMP2 | G3 | 0.006532101 | 0.943551 |
| MMP2 | GX | -0.116521739 | 0.587669 |
| MMP2 | Stage I | 0.080379187 | 0.307744 |
| MMP2 | Stage II | -0.10258302 | 0.39461 |
| MMP2 | Stage III | -0.06334423 | 0.675787 |
| MMP2 | Stage IV | -0.118012422 | 0.600939 |
| MMP2 | overall | -0.002089224 | 0.970823 |
| MMP3 | G1 | 0.126315789 | 0.595654 |
| MMP3 | G2 | -0.057258277 | 0.507894 |
| MMP3 | G3 | 0.165886183 | 0.070181 |
| MMP3 | GX | 0.212173913 | 0.319571 |
| KLF6 | Stage I | 0.161545346 | 0.039384 |
| MMP3 | Stage II | 0.076267208 | 0.527286 |
| MMP3 | Stage III | -0.112687351 | 0.455891 |
| MMP3 | Stage IV | -0.2783738 | 0.209679 |
| MMP3 | overall | 0.075870392 | 0.183455 |
| MMP7 | G1 | 0.076691729 | 0.747933 |
| MMP7 | G2 | -0.050954761 | 0.555776 |
| MMP7 | G3 | 0.122345792 | 0.18312 |
| MMP7 | GX | -0.155652174 | 0.467671 |
| MMP7 | Stage I | 0.064750594 | 0.411544 |
| MMP7 | Stage II | -0.027850669 | 0.817662 |
| MMP7 | Stage III | -0.025720101 | 0.86527 |

| | | | |
|---|---|---|---|
| MMP7 | Stage IV | 0.055900621 | 0.804842 |
| MMP7 | overall | 0.028230652 | 0.621068 |
| MMP9 | G1 | -0.309774436 | 0.183806 |
| MMP9 | G2 | 0.112355235 | 0.192811 |
| MMP9 | G3 | 0.098853149 | 0.282742 |
| MMP9 | GX | 0.307826087 | 0.143375 |
| HRAS | Stage I | 0.160669702 | 0.040478 |
| MMP9 | Stage II | 0.166584227 | 0.164993 |
| MMP9 | Stage III | -0.078023809 | 0.606269 |
| MMP9 | Stage IV | -0.06606437 | 0.77021 |
| MMP9 | overall | 0.103110194 | 0.070297 |
| MSN | G1 | -0.183458647 | 0.438795 |
| MSN | G2 | 0.090209081 | 0.296286 |
| MSN | G3 | -0.015904849 | 0.86311 |
| MSN | GX | 0.289565217 | 0.16992 |
| MSN | Stage I | 0.111184697 | 0.157664 |
| MARVELD3 | Stage I | 0.158915642 | 0.042746 |
| MSN | Stage III | -0.178807131 | 0.234453 |
| MSN | Stage IV | -0.233201581 | 0.296276 |
| MSN | overall | 0.065832001 | 0.248589 |
| MST1R | G1 | 0.016541353 | 0.944817 |
| MST1R | G2 | 0.064818768 | 0.453422 |
| MST1R | G3 | -0.146734384 | 0.109767 |
| MST1R | GX | -0.153913043 | 0.47272 |
| MST1R | Stage I | -0.047564683 | 0.546553 |
| MST1R | Stage II | 0.055583967 | 0.645228 |
| MST1R | Stage III | -0.198791095 | 0.185362 |
| MST1R | Stage IV | -0.203839639 | 0.362881 |
| MST1R | overall | -0.049822483 | 0.382776 |
| MSX2 | G1 | -0.118796992 | 0.617892 |
| CTNND1 | overall | 0.157814175 | 0.005431 |
| MSX2 | G3 | -0.019155272 | 0.835496 |
| MSX2 | GX | -0.400869565 | 0.052219 |
| MSX2 | Stage I | -0.073889442 | 0.348563 |
| MSX2 | Stage II | -0.224029377 | 0.060361 |
| MSX2 | Stage III | -0.136495406 | 0.365718 |
| MSX2 | Stage IV | -0.063805759 | 0.777872 |
| NFKB1 | Stage I | 0.157150497 | 0.045136 |
| MTA1 | G1 | 0.030075188 | 0.899837 |
| MTA1 | G2 | -0.108879692 | 0.207028 |
| MTA1 | G3 | -0.013706646 | 0.881882 |
| MTA1 | GX | -0.04 | 0.85278 |
| MTA1 | Stage I | 0.022328938 | 0.77724 |
| TXNIP | Stage I | 0.15632196 | 0.046295 |

| | | | |
|---|---|---|---|
| MTA1 | Stage III | -0.266144454 | 0.073808 |
| MTA1 | Stage IV | 0.32580463 | 0.138959 |
| MTA1 | overall | -0.088548876 | 0.120354 |
| MTA3 | G1 | 0.178947368 | 0.450325 |
| MTA3 | G2 | -0.021545031 | 0.803387 |
| MTA3 | G3 | -0.00686895 | 0.940645 |
| MTA3 | GX | -0.392173913 | 0.058036 |
| MTA3 | Stage I | -0.070364695 | 0.372098 |
| MTA3 | Stage II | -0.017136294 | 0.887206 |
| MTA3 | Stage III | 0.092950102 | 0.53895 |
| MTA3 | Stage IV | 0.092038396 | 0.683739 |
| MTA3 | overall | -0.013266255 | 0.816333 |
| MTDH | G1 | -0.293233083 | 0.209571 |
| MTDH | G2 | -0.08799542 | 0.308346 |
| MTDH | G3 | -0.090782654 | 0.324083 |
| MTDH | GX | 0.025217391 | 0.90689 |
| VDR | Stage I | 0.156305334 | 0.046319 |
| MTDH | Stage II | -0.042706595 | 0.723619 |
| MTDH | Stage III | 0.021772652 | 0.885799 |
| MTDH | Stage IV | -0.210615471 | 0.346796 |
| MTDH | overall | -0.09523427 | 0.094706 |
| MUC1 | G1 | -0.132330827 | 0.578107 |
| MUC1 | G2 | -0.072079959 | 0.404328 |
| MUC1 | G3 | -0.029753874 | 0.747 |
| MUC1 | GX | -0.19826087 | 0.353049 |
| MUC1 | Stage I | -0.034618443 | 0.660873 |
| MUC1 | Stage II | -0.027112903 | 0.822412 |
| MUC1 | Stage III | -0.178437058 | 0.235439 |
| MUC1 | Stage IV | -0.037831733 | 0.867254 |
| MUC1 | overall | -0.067427035 | 0.237285 |
| MUC16 | G1 | -0.04962406 | 0.835414 |
| MUC16 | G2 | 0.001602996 | 0.985223 |
| MUC16 | G3 | 0.060907236 | 0.508718 |
| MUC16 | GX | -0.069565217 | 0.746697 |
| MUC16 | Stage I | 0.011763011 | 0.88153 |
| MUC16 | Stage II | 0.05550013 | 0.645727 |
| MUC16 | Stage III | -8.02E-04 | 0.99578 |
| MUC16 | Stage IV | 0.274985884 | 0.215509 |
| MUC16 | overall | 0.031011742 | 0.58709 |
| MUC2 | G1 | -0.248965794 | 0.289828 |
| MUC2 | G2 | -0.003523336 | 0.967527 |
| MUC2 | G3 | -0.069011443 | 0.453879 |
| MUC2 | GX | -8.70E-04 | 0.996782 |
| MUC2 | Stage I | -0.012697339 | 0.872198 |

| | | | |
|---|---|---|---|
| MUC2 | Stage II | 0.076411654 | 0.526502 |
| MUC2 | Stage III | -0.277774351 | 0.061612 |
| MUC2 | Stage IV | -0.12365895 | 0.583509 |
| MUC2 | overall | -0.049679522 | 0.384145 |
| MUC4 | G1 | 0.222556391 | 0.345621 |
| MUC4 | G2 | 0.049821691 | 0.564608 |
| MUC4 | G3 | 0.171265356 | 0.061439 |
| MUC4 | GX | 0.022608696 | 0.916487 |
| TP73 | Stage I | 0.156197264 | 0.046472 |
| MUC4 | Stage II | -0.001961787 | 0.987045 |
| MUC4 | Stage III | 0.09813113 | 0.516461 |
| MUC4 | Stage IV | 0.036702428 | 0.871184 |
| MUC4 | overall | 0.110679232 | 0.051937 |
| MYC | G1 | -0.126315789 | 0.595654 |
| MYC | G2 | 0.084510335 | 0.327973 |
| MYC | G3 | 0.13958762 | 0.128371 |
| MMP9 | Stage I | 0.154346218 | 0.04916 |
| ID1 | overall | 0.152699411 | 0.007164 |
| MYC | Stage II | 0.174280468 | 0.146058 |
| MYC | Stage III | -0.069018689 | 0.648556 |
| MYC | Stage IV | -0.069452287 | 0.758755 |
| PTHLH | overall | 0.149536799 | 0.008469 |
| MYCN | G1 | 0.314285714 | 0.17717 |
| AXL | overall | 0.145254632 | 0.010571 |
| MYCN | G3 | 0.00103833 | 0.99102 |
| TP73 | overall | 0.143990686 | 0.011274 |
| MYCN | Stage I | -0.086972907 | 0.269618 |
| MYCN | Stage II | -0.176996789 | 0.139779 |
| MYCN | Stage III | -0.191142911 | 0.2032 |
| MYCN | Stage IV | 0.040090344 | 0.859403 |
| MYCN | overall | -0.094150519 | 0.098539 |
| NCSTN | G1 | 0.052631579 | 0.825581 |
| NCSTN | G2 | 0.039144592 | 0.650937 |
| NCSTN | G3 | 0.005809784 | 0.949785 |
| NCSTN | GX | -0.194782609 | 0.36173 |
| NCSTN | Stage I | 0.021935452 | 0.781065 |
| NCSTN | Stage II | 0.051308277 | 0.670882 |
| NCSTN | Stage III | 0.001788688 | 0.990587 |
| NCSTN | Stage IV | 0.172219085 | 0.44345 |
| NCSTN | overall | 0.012306347 | 0.829411 |
| NDRG1 | G1 | 0.126315789 | 0.595654 |
| NDRG1 | G2 | 0.164509858 | 0.055639 |
| NDRG1 | G3 | 0.153863785 | 0.093375 |
| NDRG1 | GX | -0.037391304 | 0.862289 |

| | | | |
|---|---|---|---|
| NDRG1 | Stage I | 0.088528779 | 0.261111 |
| RUNX3 | overall | 0.143854631 | 0.011352 |
| NDRG1 | Stage III | 0.045087276 | 0.76606 |
| NDRG1 | Stage IV | -0.192546584 | 0.390627 |
| LCN2 | overall | 0.141812793 | 0.012583 |
| NF1 | G1 | -0.085714286 | 0.719366 |
| NF1 | G2 | -0.063158522 | 0.465097 |
| NF1 | G3 | -0.068491557 | 0.457295 |
| TGFA | overall | 0.137971533 | 0.01522 |
| NF1 | Stage I | -0.112952613 | 0.151121 |
| NF1 | Stage II | -0.015140972 | 0.900268 |
| NF1 | Stage III | 0.020539074 | 0.89223 |
| NF1 | Stage IV | -0.225296443 | 0.313412 |
| NF1 | overall | -0.079791952 | 0.161764 |
| NFIC | G1 | -0.192481203 | 0.416206 |
| NFIC | G2 | 0.055584843 | 0.520393 |
| ITGB4 | overall | 0.136918888 | 0.016022 |
| NFIC | GX | -0.370434783 | 0.074768 |
| NFIC | Stage I | -0.006428783 | 0.935087 |
| NFIC | Stage II | -0.211386749 | 0.076791 |
| NDRG1 | overall | 0.13422057 | 0.01825 |
| NFIC | Stage IV | 0.328063241 | 0.136081 |
| NFIC | overall | -0.097724337 | 0.086349 |
| NFKB1 | G1 | 0.036090226 | 0.879931 |
| NFKB1 | G2 | 0.091339766 | 0.290249 |
| NFKB1 | G3 | 0.060264792 | 0.513207 |
| NFKB1 | GX | -0.103478261 | 0.630395 |
| YWHAZ | overall | 0.128037502 | 0.024395 |
| NFKB1 | Stage II | 0.182965987 | 0.126697 |
| NFKB1 | Stage III | -0.046074138 | 0.761089 |
| NFKB1 | Stage IV | -0.319028797 | 0.147851 |
| NFKB1 | overall | 0.080586317 | 0.157616 |
| NOTCH1 | G1 | 0.064661654 | 0.786515 |
| NOTCH1 | G2 | 0.121231349 | 0.159745 |
| MYC | overall | 0.122238396 | 0.031705 |
| NOTCH1 | GX | 0.10173913 | 0.636184 |
| VDR | overall | 0.118522209 | 0.037312 |
| NOTCH1 | Stage II | 0.213147327 | 0.07431 |
| NOTCH1 | Stage III | 0.101461791 | 0.502256 |
| NOTCH1 | Stage IV | 0.06606437 | 0.77021 |
| PTK2 | overall | 0.116262561 | 0.041116 |
| NOTCH2 | G1 | 0.042105263 | 0.860095 |
| NOTCH2 | G2 | 0.013487113 | 0.876159 |
| NOTCH2 | G3 | -0.004320007 | 0.96265 |

| | | | |
|---|---|---|---|
| NOTCH2 | GX | -0.343478261 | 0.10032 |
| NOTCH2 | Stage I | 0.008235493 | 0.916902 |
| NOTCH2 | Stage II | -0.091030274 | 0.450255 |
| NOTCH2 | Stage III | 0.138099057 | 0.360059 |
| NOTCH2 | Stage IV | -0.294184077 | 0.18387 |
| NOTCH2 | overall | -0.023341019 | 0.682767 |
| NRP2 | G1 | 0.079699248 | 0.738374 |
| NRP2 | G2 | 0.002490369 | 0.977045 |
| NRP2 | G3 | 0.067783132 | 0.461973 |
| NRP2 | GX | 0.057391304 | 0.789955 |
| NRP2 | Stage I | 0.14739371 | 0.060438 |
| NRP2 | Stage II | -0.077951693 | 0.518177 |
| NRP2 | Stage III | 0.028310615 | 0.851845 |
| NRP2 | Stage IV | 0.064935065 | 0.774038 |
| NRP2 | overall | 0.058805555 | 0.302818 |
| NUMB | G1 | -0.067669173 | 0.776821 |
| NUMB | G2 | 0.137125341 | 0.111405 |
| NUMB | G3 | -0.007431523 | 0.935794 |
| NUMB | GX | 0.217391304 | 0.307533 |
| NUMB | Stage I | 0.098213519 | 0.212297 |
| NUMB | Stage II | -0.008601682 | 0.943243 |
| NUMB | Stage III | 0.259236417 | 0.081908 |
| NUMB | Stage IV | 0.005081875 | 0.982093 |
| NUMB | overall | 0.063607129 | 0.264975 |
| OCLN | G1 | -0.05112782 | 0.830495 |
| OCLN | G2 | -0.114809823 | 0.183206 |
| OCLN | G3 | 0.039807267 | 0.665979 |
| FGFR2 | overall | 0.114688882 | 0.043954 |
| OCLN | Stage I | -0.115701472 | 0.141358 |
| OCLN | Stage II | -0.174146329 | 0.146373 |
| OCLN | Stage III | 0.211250233 | 0.158746 |
| OCLN | Stage IV | -0.120271033 | 0.593941 |
| OCLN | overall | -0.093620526 | 0.100458 |
| PAG1 | G1 | 0.070676692 | 0.767158 |
| PAG1 | G2 | -0.134689837 | 0.117965 |
| PAG1 | G3 | 0.023964926 | 0.795009 |
| PAG1 | GX | 0.100869565 | 0.639086 |
| PAG1 | Stage I | -0.110514109 | 0.160201 |
| PAG1 | Stage II | 0.072368145 | 0.548674 |
| PAG1 | Stage III | 0.001048541 | 0.994482 |
| PAG1 | Stage IV | -0.003952569 | 0.986072 |
| PAG1 | overall | -0.036622527 | 0.521281 |
| PAK1 | G1 | 0.019548872 | 0.934803 |
| PAK1 | G2 | 0.084128669 | 0.33017 |

| | | | |
|---|---|---|---|
| PAK1 | G3 | 0.03727916 | 0.686037 |
| PAK1 | GX | -0.236521739 | 0.26582 |
| PAK1 | Stage I | 0.016839533 | 0.831051 |
| PAK1 | Stage II | 0.077465438 | 0.520798 |
| PAK1 | Stage III | 0.099981497 | 0.508545 |
| PAK1 | Stage IV | 0.032185206 | 0.886933 |
| PAK1 | overall | 0.045271055 | 0.427793 |
| PARD3 | G1 | -0.02556391 | 0.914804 |
| ROCK2 | overall | -0.111719471 | 0.049758 |
| PARD3 | G3 | 0.072453879 | 0.431622 |
| PARD3 | GX | -0.322608696 | 0.124164 |
| PARD3 | Stage I | 0.126015235 | 0.108963 |
| PARD3 | Stage II | 0.056640314 | 0.638951 |
| PARD3 | Stage III | 0.222722508 | 0.136813 |
| PARD3 | Stage IV | -0.079616036 | 0.724693 |
| PARD3 | overall | 0.100880238 | 0.07662 |
| PARP1 | G1 | 0.072180451 | 0.762339 |
| PARP1 | G2 | 0.055222261 | 0.523122 |
| PARP1 | G3 | 0.014442853 | 0.875587 |
| TCF3 | overall | -0.112348292 | 0.048478 |
| PARP1 | Stage I | 0.013151295 | 0.867669 |
| PARP1 | Stage II | 0.032797056 | 0.785991 |
| PARP1 | Stage III | -0.18115093 | 0.228276 |
| PARP1 | Stage IV | 0.049124788 | 0.828133 |
| PARP1 | overall | -0.007768544 | 0.891816 |
| PCMT1 | G1 | 0.019548872 | 0.934803 |
| PCMT1 | G2 | -0.062979617 | 0.466365 |
| AXIN1 | overall | -0.113152215 | 0.046883 |
| PCMT1 | GX | -0.067826087 | 0.752835 |
| PCMT1 | Stage I | -0.142910188 | 0.068779 |
| PCMT1 | Stage II | 0.001743811 | 0.988485 |
| PCMT1 | Stage III | -0.135261828 | 0.370108 |
| PCMT1 | Stage IV | -0.258046302 | 0.246268 |
| IDH2 | overall | -0.115828569 | 0.041883 |
| PDE4A | G1 | -0.081203008 | 0.733608 |
| PDE4A | G2 | -0.095099173 | 0.270767 |
| PDE4A | G3 | -0.09090767 | 0.323414 |
| PDE4A | GX | 0.22173913 | 0.297718 |
| PDE4A | Stage I | -0.080085458 | 0.309521 |
| PDE4A | Stage II | 0.099849932 | 0.407393 |
| PDE4A | Stage III | -0.181521003 | 0.227311 |
| PDE4A | Stage IV | -0.209486166 | 0.349447 |
| PDE4A | overall | -0.049209728 | 0.388663 |
| PDGFB | G1 | 0.138345865 | 0.560787 |

| | | | |
|---|---|---|---|
| PDGFB | G2 | -0.075908543 | 0.379762 |
| PDGFB | G3 | -0.018596171 | 0.840233 |
| PDGFB | GX | -0.226956522 | 0.2862 |
| PDGFB | Stage I | 0.008590184 | 0.913337 |
| PDGFB | Stage II | -0.221598102 | 0.063277 |
| PDGFB | Stage III | -0.087028928 | 0.565219 |
| PDGFB | Stage IV | 0.137210615 | 0.542591 |
| PDGFB | overall | -0.05960175 | 0.296307 |
| PDPN | G1 | 0.012030075 | 0.959853 |
| DDX5 | overall | -0.11620094 | 0.041224 |
| MSX2 | overall | -0.117059773 | 0.039738 |
| PDPN | GX | 0.375652174 | 0.070454 |
| RAF1 | overall | -0.117332289 | 0.039276 |
| ILK | overall | -0.118228339 | 0.037789 |
| PDPN | Stage III | 0.070498983 | 0.641524 |
| PDPN | Stage IV | 0.049124788 | 0.828133 |
| VEGFA | overall | -0.122197112 | 0.031763 |
| PHLDA1 | G1 | -0.084210526 | 0.724104 |
| PHLDA1 | G2 | 0.14356118 | 0.095436 |
| PHLDA1 | G3 | 0.029604549 | 0.748227 |
| PHLDA1 | GX | 0.4 | 0.05278 |
| KRAS | overall | -0.13024224 | 0.022025 |
| PHLDA1 | Stage II | 0.103119577 | 0.392129 |
| PHLDA1 | Stage III | -0.133411461 | 0.376751 |
| PHLDA1 | Stage IV | -0.062676454 | 0.78171 |
| PHLDA1 | overall | 0.092307364 | 0.10534 |
| CXCR4 | overall | -0.136018568 | 0.016737 |
| PIK3CA | G2 | 0.146476152 | 0.088828 |
| PIK3CA | G3 | 0.009237314 | 0.920239 |
| PIK3CA | GX | 0.023478261 | 0.913287 |
| PIK3CA | Stage I | 0.082172042 | 0.297044 |
| PCMT1 | overall | -0.137919064 | 0.015259 |
| PIK3CA | Stage III | 0.175969902 | 0.242083 |
| PIK3CA | Stage IV | -0.199322417 | 0.37384 |
| PIK3CA | overall | 0.10819852 | 0.057456 |
| PIN1 | G1 | -0.207518797 | 0.379993 |
| PIN1 | G2 | 0.053046766 | 0.539646 |
| PIN1 | G3 | 0.091564006 | 0.319918 |
| PIN1 | GX | -0.130434783 | 0.543526 |
| PIN1 | Stage I | 0.023933362 | 0.761697 |
| PIN1 | Stage II | -0.017085992 | 0.887535 |
| PIN1 | Stage III | -0.010177019 | 0.946481 |
| PIN1 | Stage IV | -0.009599097 | 0.966183 |
| PIN1 | overall | 0.033174789 | 0.56127 |

| | | | |
|---|---|---|---|
| PKP3 | G1 | 0.189473684 | 0.423665 |
| PLXND1 | overall | -0.138005293 | 0.015195 |
| PKP3 | G3 | 0.069981335 | 0.447545 |
| MAP3K7 | overall | -0.138309942 | 0.014969 |
| DAPK1 | overall | -0.139321912 | 0.014242 |
| ATM | overall | -0.142708232 | 0.012029 |
| PKP3 | Stage III | 0.033738358 | 0.823854 |
| PKP3 | Stage IV | -0.295313382 | 0.182113 |
| RGS3 | overall | -0.142967733 | 0.011873 |
| SEMA4C | overall | -0.145034789 | 0.01069 |
| PLAUR | G2 | 0.006900994 | 0.936447 |
| PLAUR | G3 | -0.03274732 | 0.722538 |
| PLAUR | GX | 0.20173913 | 0.344493 |
| PLAUR | Stage I | 0.050402215 | 0.522858 |
| PLAUR | Stage II | 0.066180971 | 0.583449 |
| PLAUR | Stage III | -0.124653057 | 0.409146 |
| PLAUR | Stage IV | -0.239977414 | 0.282058 |
| PLAUR | overall | -0.009609656 | 0.866393 |
| PLXND1 | G1 | -0.106766917 | 0.654142 |
| PLXND1 | G2 | -0.154598476 | 0.072321 |
| PLXND1 | G3 | -0.127909016 | 0.163852 |
| PLXND1 | GX | -0.142608696 | 0.506209 |
| PLXND1 | Stage I | -0.090640301 | 0.249863 |
| PLXND1 | Stage II | -0.213801256 | 0.073404 |
| PLXND1 | Stage III | -0.254425463 | 0.08795 |
| PLXND1 | Stage IV | -0.193675889 | 0.3878 |
| LRP6 | overall | -0.146650973 | 0.00984 |
| POSTN | G1 | -0.027067669 | 0.909812 |
| POSTN | G2 | -0.019920566 | 0.817944 |
| POSTN | G3 | -0.128853584 | 0.160736 |
| POSTN | GX | -0.048695652 | 0.821234 |
| POSTN | Stage I | 0.009615465 | 0.903042 |
| POSTN | Stage II | -0.149330561 | 0.213894 |
| POSTN | Stage III | -0.106026029 | 0.48312 |
| POSTN | Stage IV | -0.139469226 | 0.535903 |
| POSTN | overall | -0.061338329 | 0.282432 |
| POU5F1 | G1 | 0.156390977 | 0.510257 |
| POU5F1 | G2 | 0.051925623 | 0.548261 |
| POU5F1 | G3 | 0.173508704 | 0.058066 |
| POU5F1 | GX | -0.052173913 | 0.808688 |
| POU5F1 | Stage I | 0.113304533 | 0.149844 |
| POU5F1 | Stage II | 0.125520838 | 0.296941 |
| POU5F1 | Stage III | 0.023376303 | 0.877449 |
| POU5F1 | Stage IV | 0.355166573 | 0.104797 |

| | | | |
|---|---|---|---|
| POU5F1 | overall | 0.108534895 | 0.05668 |
| PPARG | G1 | 0.109774436 | 0.645005 |
| PPARG | G2 | -0.146693224 | 0.088351 |
| PPARG | G3 | -0.072589313 | 0.430759 |
| PPARG | GX | 0.247826087 | 0.242965 |
| PPARG | Stage I | -0.042260937 | 0.592208 |
| CDKN1B | overall | -0.150556498 | 0.008027 |
| PPARG | Stage III | -0.181027572 | 0.228598 |
| PPARG | Stage IV | -0.016374929 | 0.942342 |
| PPARG | overall | -0.074650343 | 0.190622 |
| PRDX1 | G1 | -0.022556391 | 0.924798 |
| CTNNB1 | overall | -0.151474105 | 0.007647 |
| PRDX1 | G3 | 0.004431133 | 0.96169 |
| PRDX1 | GX | 8.70E-04 | 0.996782 |
| PRDX1 | Stage I | 0.069522303 | 0.377864 |
| PRDX1 | Stage II | 0.14111453 | 0.240463 |
| GLRX | overall | -0.151659782 | 0.007572 |
| PRDX1 | Stage IV | -0.025409373 | 0.910632 |
| PRDX1 | overall | 0.106596368 | 0.061271 |
| PRKCA | G1 | -0.027067669 | 0.909812 |
| PRKCA | G2 | -0.122598189 | 0.155055 |
| PRKCA | G3 | -0.020631159 | 0.823021 |
| PRKCA | GX | -0.166086957 | 0.437967 |
| PRKCA | Stage I | -0.07523339 | 0.339842 |
| PRKCA | Stage II | -0.160514424 | 0.181157 |
| PRKCA | Stage III | 0.067538396 | 0.655618 |
| PRKCA | Stage IV | -0.214003388 | 0.338913 |
| PRKCA | overall | -0.06809592 | 0.232653 |
| PRKCE | G1 | 0.12481203 | 0.600074 |
| PRKCE | G2 | 0.062209129 | 0.471847 |
| PRKCE | G3 | 0.125266311 | 0.172808 |
| PRKCE | GX | 0.006086957 | 0.97748 |
| ERBB2 | overall | -0.153108185 | 0.007009 |
| PRKCE | Stage II | -0.044701917 | 0.71126 |
| PRKCE | Stage III | 0.086412139 | 0.567989 |
| PRKCE | Stage IV | -0.376623377 | 0.084045 |
| PRKCE | overall | 0.097701152 | 0.086424 |
| PROM1 | G1 | -0.054155701 | 0.820608 |
| CMTM8 | overall | -0.157227452 | 0.005608 |
| PROM1 | G3 | -0.100928088 | 0.27271 |
| TRPS1 | overall | -0.157912403 | 0.005401 |
| PROM1 | Stage I | -0.122242674 | 0.120062 |
| CDX2 | overall | -0.159448308 | 0.004962 |
| PROM1 | Stage III | -0.221118857 | 0.139735 |

| | | | |
|---|---|---|---|
| PROM1 | Stage IV | 0.106719368 | 0.636432 |
| SON | overall | -0.160578792 | 0.00466 |
| PRSS8 | G1 | -0.165413534 | 0.485838 |
| PRSS8 | G2 | -0.016421168 | 0.849507 |
| PRSS8 | G3 | 0.029052394 | 0.75277 |
| PRSS8 | GX | -0.28173913 | 0.182283 |
| PRSS8 | Stage I | -0.057199544 | 0.468299 |
| PRSS8 | Stage II | -0.040610669 | 0.736677 |
| PRSS8 | Stage III | 0.073829643 | 0.625817 |
| PRSS8 | Stage IV | 0.138339921 | 0.539242 |
| PRSS8 | overall | -0.020383607 | 0.72117 |
| PTEN | G1 | -0.027067669 | 0.909812 |
| PTEN | G2 | 0.016554751 | 0.848298 |
| PTEN | G3 | -0.037161089 | 0.686979 |
| PTEN | GX | 0.173913043 | 0.41637 |
| PTEN | Stage I | 0.121057867 | 0.123722 |
| PTEN | Stage II | 0.010077214 | 0.933528 |
| PTEN | Stage III | -0.057052983 | 0.706463 |
| PTEN | Stage IV | -0.250141163 | 0.261546 |
| PTEN | overall | 0.024770306 | 0.664489 |
| PTGS2 | G1 | 0.165413534 | 0.485838 |
| PTGS2 | G2 | 0.060684851 | 0.482794 |
| PTGS2 | G3 | 0.09150497 | 0.320232 |
| PTGS2 | GX | -0.046956522 | 0.827524 |
| PTGS2 | Stage I | 0.144251365 | 0.066191 |
| PTGS2 | Stage II | 0.090694925 | 0.451933 |
| PTGS2 | Stage III | -0.225559738 | 0.131755 |
| PTGS2 | Stage IV | 0.037831733 | 0.867254 |
| PTGS2 | overall | 0.061474994 | 0.281359 |
| PTHLH | G1 | 0.027067669 | 0.909812 |
| PTHLH | G2 | 0.164354806 | 0.055873 |
| ELF5 | Stage I | -0.161076736 | 0.039966 |
| PTHLH | GX | 0.085217391 | 0.692169 |
| PTHLH | Stage I | 0.153520452 | 0.050401 |
| FLT1 | Stage I | -0.161861244 | 0.038995 |
| PTHLH | Stage III | 0.072225992 | 0.63336 |
| PTHLH | Stage IV | 0.019762846 | 0.930438 |
| MAP3K7 | Stage I | -0.167400527 | 0.03269 |
| PTK2 | G1 | 0.111278195 | 0.640455 |
| PTK2 | G2 | 0.112090455 | 0.193868 |
| PTK2 | G3 | 0.12395364 | 0.177388 |
| PTK2 | GX | -0.066956522 | 0.75591 |
| PTK2 | Stage I | 0.096636804 | 0.219764 |
| PTK2 | Stage II | 0.169317315 | 0.158072 |

| | | | |
|---|---|---|---|
| PTK2 | Stage III | 0.092086598 | 0.542744 |
| PTK2 | Stage IV | 0.044607566 | 0.843739 |
| RHOA | Stage I | -0.168159789 | 0.031897 |
| PTN | G1 | 0.335338346 | 0.148357 |
| PTN | G2 | -0.051565426 | 0.551043 |
| PTN | G3 | 0.006660589 | 0.942443 |
| ATM | G2 | -0.169946209 | 0.047927 |
| PTN | Stage I | -0.064426384 | 0.413894 |
| PTN | Stage II | 0.024966675 | 0.836267 |
| PTN | Stage III | -0.22543638 | 0.131972 |
| PTN | Stage IV | 0.194805195 | 0.384985 |
| PTN | overall | -0.037772179 | 0.508279 |
| PTPN14 | G1 | 0.270676692 | 0.248391 |
| PTPN14 | G2 | 0.129272569 | 0.133627 |
| SMAD2 | Stage I | -0.170819975 | 0.029246 |
| PTPN14 | GX | 0.062608696 | 0.771334 |
| RHOA | overall | -0.172516632 | 0.002341 |
| MTDH | Stage I | -0.172562951 | 0.027614 |
| PTPN14 | Stage III | 0.121199039 | 0.422348 |
| PTPN14 | Stage IV | 0.114624506 | 0.6115 |
| SIM2 | Stage I | -0.172637769 | 0.027545 |
| PTPRZ1 | G1 | 0.321804511 | 0.166474 |
| PTPRZ1 | G2 | 0.155546226 | 0.070569 |
| ENG | G2 | -0.172811087 | 0.044234 |
| LATS1 | overall | -0.174622125 | 0.002064 |
| CDX2 | Stage I | -0.17487872 | 0.025565 |
| MYCN | G2 | -0.175962215 | 0.040449 |
| PTPRZ1 | Stage III | 0.173323054 | 0.249351 |
| PTPRZ1 | Stage IV | 0.070581592 | 0.754948 |
| TCF7 | G2 | -0.177512732 | 0.038689 |
| RAC1 | G1 | 0.213533835 | 0.366021 |
| DAB2 | G2 | -0.177996971 | 0.038153 |
| RAC1 | G3 | 0.097224465 | 0.290786 |
| RAC1 | GX | 0.031304348 | 0.884547 |
| RAC1 | Stage I | 0.130850678 | 0.095933 |
| FBLN5 | G2 | -0.178972604 | 0.037091 |
| PROM1 | overall | -0.179276487 | 0.001555 |
| RAC1 | Stage IV | -0.329192547 | 0.134658 |
| MSX2 | G2 | -0.180640006 | 0.035334 |
| RAF1 | G1 | 0.287218045 | 0.219504 |
| RAF1 | G2 | -0.045778419 | 0.596657 |
| TCF7 | overall | -0.181458216 | 0.001358 |
| ERBB2 | Stage I | -0.181527224 | 0.020392 |
| RAF1 | Stage I | -0.087880358 | 0.264633 |

| | | | |
|---|---|---|---|
| CMTM8 | G2 | -0.182479157 | 0.033478 |
| RAF1 | Stage III | -0.109480048 | 0.468896 |
| RAF1 | Stage IV | 0.227555054 | 0.308456 |
| FLT1 | overall | -0.183488056 | 0.001196 |
| RDX | G1 | -0.132330827 | 0.578107 |
| RDX | G2 | -0.151292296 | 0.078712 |
| RDX | G3 | -0.03235838 | 0.725701 |
| RDX | GX | -0.150434783 | 0.482902 |
| RDX | Stage I | -0.138784129 | 0.077262 |
| RDX | Stage II | -0.083350799 | 0.489525 |
| RDX | Stage III | 0.059150065 | 0.696184 |
| RDX | Stage IV | 0.022021457 | 0.92251 |
| RDX | overall | -0.098764576 | 0.083038 |
| RGS3 | G1 | -0.033082707 | 0.889876 |
| RGS3 | G2 | -0.160149327 | 0.06254 |
| RGS3 | G3 | -0.129818987 | 0.157597 |
| RGS3 | GX | -0.249565217 | 0.239566 |
| RGS3 | Stage I | -0.097287996 | 0.216658 |
| RGS3 | Stage II | -0.161151586 | 0.179408 |
| RGS3 | Stage III | -0.235305004 | 0.115444 |
| RGS3 | Stage IV | -0.151891587 | 0.499815 |
| SIM2 | G3 | -0.185871425 | 0.042097 |
| RHOA | G1 | 0.145864662 | 0.539466 |
| RHOA | G2 | -0.051651301 | 0.55038 |
| FLT1 | G2 | -0.187025751 | 0.029241 |
| GLRX | G3 | -0.188850979 | 0.038854 |
| LATS1 | Stage I | -0.189011769 | 0.015675 |
| RHOA | Stage II | -0.184407985 | 0.123681 |
| RHOA | Stage III | -0.134521681 | 0.372756 |
| RHOA | Stage IV | -0.114624506 | 0.6115 |
| RAF1 | G3 | -0.189121848 | 0.03857 |
| RNF111 | G1 | -0.187969925 | 0.427421 |
| RNF111 | G2 | -0.029390647 | 0.734112 |
| RNF111 | G3 | 0.040828233 | 0.657944 |
| CXCR4 | G3 | -0.198817555 | 0.029486 |
| RNF111 | Stage I | -0.017593252 | 0.82361 |
| RNF111 | Stage II | -0.121446357 | 0.313022 |
| RNF111 | Stage III | 0.215814472 | 0.149732 |
| RNF111 | Stage IV | -0.162055336 | 0.471204 |
| RNF111 | overall | -0.044142756 | 0.439409 |
| ROCK1 | G1 | 0.163909774 | 0.489867 |
| ROCK1 | G2 | -0.105699939 | 0.220681 |
| ROCK1 | G3 | -0.117102053 | 0.202752 |
| ROCK1 | GX | -0.286956522 | 0.173974 |

| | | | |
|---|---|---|---|
| ROCK1 | Stage I | -0.107008759 | 0.17396 |
| ROCK1 | Stage II | -0.086352166 | 0.473967 |
| ROCK1 | Stage III | -0.141923149 | 0.34678 |
| ROCK1 | Stage IV | 0.043478261 | 0.84765 |
| ROCK1 | overall | -0.10709727 | 0.060056 |
| ROCK2 | G1 | -0.030075188 | 0.899837 |
| GAB2 | overall | -0.198935661 | 4.35E-04 |
| ROCK2 | G3 | 0.006087598 | 0.947387 |
| ROCK2 | GX | -0.324347826 | 0.122033 |
| ROCK2 | Stage I | -0.141621661 | 0.071343 |
| ROCK2 | Stage II | -0.19570922 | 0.10191 |
| ROCK2 | Stage III | 0.176956764 | 0.23941 |
| ROCK2 | Stage IV | -0.221908526 | 0.320937 |
| IL6R | G3 | -0.199605852 | 0.028835 |
| ROR2 | G1 | -0.210526316 | 0.37297 |
| ROR2 | G2 | 0.032470212 | 0.707461 |
| ROR2 | G3 | 0.05122542 | 0.578458 |
| ROR2 | GX | 0.403478261 | 0.050566 |
| ROR2 | Stage I | 0.082077827 | 0.2976 |
| ROR2 | Stage II | 0.041968829 | 0.728207 |
| ROR2 | Stage III | -0.00339234 | 0.982149 |
| ROR2 | Stage IV | 0.363071711 | 0.096757 |
| ROR2 | overall | 0.079713451 | 0.162179 |
| RUNX3 | G1 | -0.079699248 | 0.738374 |
| RUNX3 | G2 | 0.13871164 | 0.107287 |
| RUNX3 | G3 | 0.164361679 | 0.072834 |
| RUNX3 | GX | 0.01826087 | 0.932507 |
| RUNX3 | Stage I | 0.101716097 | 0.196363 |
| MAP3K7 | G3 | -0.203342449 | 0.025913 |
| RUNX3 | Stage III | 0.115771296 | 0.443574 |
| RUNX3 | Stage IV | -0.296442688 | 0.180368 |
| PCMT1 | G3 | -0.204439815 | 0.025104 |
| S100A4 | G1 | -0.054135338 | 0.820675 |
| S100A4 | G2 | -0.016511814 | 0.848686 |
| S100A4 | G3 | -0.095012372 | 0.301954 |
| S100A4 | GX | -0.245217391 | 0.248121 |
| S100A4 | Stage I | -0.034061467 | 0.665999 |
| S100A4 | Stage II | -0.204076158 | 0.087809 |
| S100A4 | Stage III | 0.044347129 | 0.769795 |
| S100A4 | Stage IV | -0.271597967 | 0.221446 |
| S100A4 | overall | -0.048454817 | 0.39599 |
| SCRIB | G1 | 0.115789474 | 0.626879 |
| SCRIB | G2 | -0.013842539 | 0.872922 |
| SCRIB | G3 | 0.042276338 | 0.646614 |

| | | | |
|---|---|---|---|
| SCRIB | GX | 0.089565217 | 0.677269 |
| SCRIB | Stage I | -0.00962932 | 0.902902 |
| SCRIB | Stage II | -0.084105333 | 0.485589 |
| SCRIB | Stage III | 0.265527665 | 0.074504 |
| SCRIB | Stage IV | -0.232072276 | 0.298688 |
| SCRIB | overall | 0.010194142 | 0.85835 |
| SDC1 | G1 | 0.117293233 | 0.622379 |
| SON | G3 | -0.204825281 | 0.024825 |
| TCF7 | G3 | -0.206408821 | 0.023707 |
| SDC1 | GX | 0.275652174 | 0.192317 |
| SEMA4C | G2 | -0.206652911 | 0.015786 |
| FLT1 | G3 | -0.208568824 | 0.022252 |
| SDC1 | Stage III | 0.184851664 | 0.218752 |
| SDC1 | Stage IV | -0.29079616 | 0.189208 |
| TRPS1 | Stage I | -0.216176149 | 0.005579 |
| SDC2 | G1 | 0.088721805 | 0.709921 |
| SDC2 | G2 | -0.066168911 | 0.44405 |
| SDC2 | G3 | -0.131364327 | 0.152669 |
| SDC2 | GX | -0.303478261 | 0.149409 |
| SDC2 | Stage I | -0.070977092 | 0.367941 |
| SDC2 | Stage II | -0.194686408 | 0.103748 |
| SDC2 | Stage III | -0.150804911 | 0.317119 |
| SDC2 | Stage IV | 0.162055336 | 0.471204 |
| SDC2 | overall | -0.105131695 | 0.064939 |
| SEMA4C | G1 | 0.021052632 | 0.929799 |
| NFIC | G3 | -0.216271216 | 0.017669 |
| SEMA4C | G3 | -0.112052785 | 0.223045 |
| SEMA4C | GX | -0.393913043 | 0.056835 |
| SEMA4C | Stage I | -0.059069988 | 0.453854 |
| ROCK2 | G2 | -0.217895352 | 0.010824 |
| SMAD2 | G3 | -0.218882667 | 0.016313 |
| SEMA4C | Stage IV | -0.140598532 | 0.532573 |
| CDKN1B | Stage I | -0.219182713 | 0.004938 |
| SEMA7A | G1 | 0.036090226 | 0.879931 |
| SEMA7A | G2 | -0.029922593 | 0.729485 |
| SEMA7A | G3 | 0.124009203 | 0.177193 |
| SEMA7A | GX | 0.00173913 | 0.993565 |
| TRPS1 | G3 | -0.222021965 | 0.014803 |
| SEMA7A | Stage II | 0.038715952 | 0.748547 |
| SEMA7A | Stage III | 0.005242707 | 0.972415 |
| SEMA7A | Stage IV | -0.207227555 | 0.354786 |
| SEMA7A | overall | 0.068312916 | 0.231165 |
| SHH | G1 | -0.231578947 | 0.325899 |
| SHH | G2 | -0.008688805 | 0.920031 |

| | | | |
|---|---|---|---|
| SHH | G3 | -0.016051306 | 0.861862 |
| GAB2 | Stage I | -0.222699147 | 0.004272 |
| SHH | Stage I | -0.04881439 | 0.536053 |
| SHH | Stage II | -0.184009266 | 0.12451 |
| SHH | Stage III | 0.133267524 | 0.37727 |
| SHH | Stage IV | 0.055083228 | 0.807644 |
| SHH | overall | -0.050159503 | 0.379562 |
| SIM2 | G1 | 0.196992481 | 0.405152 |
| SIM2 | G2 | -0.059291772 | 0.492917 |
| ERBB2 | G3 | -0.226585059 | 0.012825 |
| SIM2 | GX | -0.302608696 | 0.150636 |
| DAPK1 | G2 | -0.230740789 | 0.00688 |
| SIM2 | Stage II | 0.022032378 | 0.855288 |
| SIM2 | Stage III | -0.084808488 | 0.57522 |
| SIM2 | Stage IV | -0.232072276 | 0.298688 |
| SIM2 | overall | -0.098429625 | 0.084092 |
| SIX1 | G1 | -0.045112782 | 0.850206 |
| SIX1 | G2 | 0.054995647 | 0.524831 |
| SIX1 | G3 | 0.140177975 | 0.126749 |
| SIX1 | GX | -0.22 | 0.30162 |
| SIX1 | Stage I | 0.079517398 | 0.312977 |
| SIX1 | Stage II | 0.099263072 | 0.410169 |
| SIX1 | Stage III | -0.013507679 | 0.929005 |
| SIX1 | Stage IV | 0.335968379 | 0.126344 |
| SIX1 | overall | 0.065362215 | 0.251989 |
| SLC39A6 | G1 | 0.103759398 | 0.663326 |
| SLC39A6 | G2 | 0.078122205 | 0.365981 |
| SLC39A6 | G3 | -0.07253375 | 0.431113 |
| SLC39A6 | GX | 0.174782609 | 0.414007 |
| SLC39A6 | Stage I | 0.015085472 | 0.848426 |
| SLC39A6 | Stage II | 0.094081942 | 0.435149 |
| SLC39A6 | Stage III | 0.011410597 | 0.940005 |
| SLC39A6 | Stage IV | 0.099943535 | 0.65811 |
| SLC39A6 | overall | 0.018426168 | 0.746985 |
| SMAD2 | G1 | 0.163909774 | 0.489867 |
| SMAD2 | G2 | -0.010951421 | 0.899305 |
| LRP6 | Stage I | -0.233858074 | 0.00266 |
| SMAD2 | GX | -0.32173913 | 0.125239 |
| PPARG | Stage II | -0.23410659 | 0.049414 |
| SMAD2 | Stage II | -0.102381811 | 0.395543 |
| SMAD2 | Stage III | 0.126626782 | 0.401709 |
| SMAD2 | Stage IV | 0.024280068 | 0.91459 |
| SMAD2 | overall | -0.105348081 | 0.064386 |
| SMAD3 | G1 | -0.171428571 | 0.469886 |

| | | | |
|---|---|---|---|
| FBLN5 | Stage II | -0.234291032 | 0.04923 |
| SMAD4 | Stage II | -0.234509008 | 0.049013 |
| SMAD3 | GX | 0.352173913 | 0.091458 |
| KRAS | Stage I | -0.237005961 | 0.002318 |
| SMAD3 | Stage II | 0.233234685 | 0.050292 |
| HDAC6 | Stage II | -0.237577444 | 0.046044 |
| SMAD3 | Stage IV | -0.129305477 | 0.566299 |
| PROM1 | G2 | -0.240181006 | 0.004856 |
| SMAD4 | G1 | 0.057142857 | 0.81088 |
| SMAD4 | G2 | -0.026167957 | 0.762345 |
| SMAD4 | G3 | -0.015001953 | 0.870812 |
| SMAD4 | GX | -0.336521739 | 0.107856 |
| SMAD4 | Stage I | -0.041906245 | 0.595322 |
| RAF1 | Stage II | -0.240494974 | 0.043359 |
| SMAD4 | Stage III | 0.281070748 | 0.058465 |
| SMAD4 | Stage IV | 0.225296443 | 0.313412 |
| SMAD4 | overall | -0.04101797 | 0.472504 |
| SMAD7 | G1 | 0.248120301 | 0.291523 |
| SMAD7 | G2 | -0.128533092 | 0.135883 |
| SMAD7 | G3 | 0.008535834 | 0.926278 |
| SMAD7 | GX | 0.133043478 | 0.535424 |
| SMAD7 | Stage I | 0.102885471 | 0.191241 |
| SMAD7 | Stage II | -0.20924052 | 0.079905 |
| SMAD7 | Stage III | 0.032998212 | 0.827659 |
| SMAD7 | Stage IV | -0.132693394 | 0.556083 |
| SMAD7 | overall | -0.009412794 | 0.869105 |
| SNAI1 | G1 | 0.207518797 | 0.379993 |
| SNAI1 | G2 | -0.116028768 | 0.178569 |
| SNAI1 | G3 | -0.12265486 | 0.182008 |
| SNAI1 | GX | 0.247826087 | 0.242965 |
| SNAI1 | Stage I | 0.032301865 | 0.682293 |
| SNAI1 | Stage II | -0.166986645 | 0.163961 |
| SNAI1 | Stage III | -0.227286747 | 0.128745 |
| SNAI1 | Stage IV | -0.106719368 | 0.636432 |
| SNAI1 | overall | -0.068669831 | 0.228731 |
| SNAI2 | G1 | -0.073684211 | 0.757529 |
| CXCL12 | Stage II | -0.240662648 | 0.043209 |
| LOXL3 | G2 | -0.241253295 | 0.004663 |
| DDX5 | Stage II | -0.245558732 | 0.039008 |
| CDX2 | G3 | -0.248353166 | 0.006236 |
| CTGF | Stage II | -0.250052398 | 0.035456 |
| SNAI2 | Stage III | 0.092333313 | 0.541659 |
| SNAI2 | Stage IV | -0.012987013 | 0.954258 |
| SEMA4C | Stage II | -0.250303909 | 0.035265 |

| | | | |
|---|---|---|---|
| SNW1 | G1 | 0.118796992 | 0.617892 |
| SNW1 | G2 | 0.115503978 | 0.180555 |
| SNW1 | G3 | 0.017148066 | 0.852527 |
| SNW1 | GX | 0.160869565 | 0.452691 |
| SNW1 | Stage I | 0.087741807 | 0.26539 |
| SNW1 | Stage II | 0.073240051 | 0.543855 |
| SNW1 | Stage III | 0.098007772 | 0.516991 |
| SNW1 | Stage IV | 0.037831733 | 0.867254 |
| SNW1 | overall | 0.061558783 | 0.280703 |
| SON | G1 | -0.240601504 | 0.306861 |
| SON | G2 | -0.035623725 | 0.680533 |
| KIT | Stage II | -0.254294553 | 0.032354 |
| MAPK3 | Stage II | -0.256658758 | 0.030727 |
| SON | Stage I | -0.15165555 | 0.053298 |
| AGER | Stage II | -0.257379757 | 0.030244 |
| SON | Stage III | 0.019798927 | 0.896092 |
| SON | Stage IV | -0.163184641 | 0.468077 |
| TCF7 | Stage II | -0.267624645 | 0.024049 |
| SOX9 | G1 | 0.356390977 | 0.122986 |
| SOX9 | G2 | -0.147902628 | 0.085729 |
| SOX9 | G3 | -0.06192473 | 0.50165 |
| SOX9 | GX | -0.086086957 | 0.68918 |
| SOX9 | Stage I | -0.125630062 | 0.110058 |
| SOX9 | Stage II | -0.069433848 | 0.565041 |
| SOX9 | Stage III | 0.016221551 | 0.91479 |
| SOX9 | Stage IV | 0.005081875 | 0.982093 |
| SOX9 | overall | -0.077659817 | 0.173305 |
| SP1 | G1 | 0.123308271 | 0.604509 |
| SP1 | G2 | -0.009050249 | 0.916716 |
| SP1 | G3 | 0.020693667 | 0.822493 |
| SP1 | GX | -0.197391304 | 0.355208 |
| SP1 | Stage I | 0.044281015 | 0.574616 |
| SP1 | Stage II | -0.111603887 | 0.354135 |
| CMTM8 | Stage II | -0.26765818 | 0.024031 |
| SP1 | Stage IV | -0.085262564 | 0.70598 |
| SP1 | overall | 0.013619509 | 0.811532 |
| SPP1 | G1 | -0.027067669 | 0.909812 |
| SPP1 | G2 | -0.036210536 | 0.675565 |
| SPP1 | G3 | -0.046978339 | 0.61039 |
| SPP1 | GX | 0.099130435 | 0.644906 |
| SPP1 | Stage I | -0.101547065 | 0.197111 |
| SPP1 | Stage II | 0.105198736 | 0.382602 |
| SPP1 | Stage III | 0.17720348 | 0.238745 |
| SPP1 | Stage IV | -0.032185206 | 0.886933 |

| | | | |
|---|---|---|---|
| SPP1 | overall | -0.008038823 | 0.888076 |
| SPRR2A | G1 | -0.064661654 | 0.786515 |
| LEFTY1 | Stage II | -0.271570987 | 0.021968 |
| SPRR2A | G3 | 0.166851308 | 0.068542 |
| SPRR2A | GX | 0.249565217 | 0.239566 |
| ID2 | overall | -0.272180105 | 1.19E-06 |
| SPRR2A | Stage II | 0.216860197 | 0.069287 |
| SPRR2A | Stage III | 0.152901993 | 0.310357 |
| SPRR2A | Stage IV | -0.095426313 | 0.672712 |
| SON | Stage II | -0.275455026 | 0.020072 |
| SRC | G1 | -0.144360902 | 0.543701 |
| SRC | G2 | -0.1292511 | 0.133692 |
| SRC | G3 | 0.13505578 | 0.141366 |
| SRC | GX | -0.131304348 | 0.540819 |
| SRC | Stage I | -0.056418115 | 0.474408 |
| SRC | Stage II | 9.56E-04 | 0.993689 |
| SRC | Stage III | 0.117868378 | 0.435304 |
| SRC | Stage IV | -0.263692829 | 0.235716 |
| SRC | overall | -0.017364778 | 0.761104 |
| SRF | G1 | 0.181954887 | 0.442621 |
| SRF | G2 | -0.008146178 | 0.92501 |
| SRF | G3 | -0.117543083 | 0.201045 |
| SRF | GX | 0.20173913 | 0.344493 |
| SRF | Stage I | 0.042726469 | 0.588132 |
| SRF | Stage II | -0.106087409 | 0.378572 |
| SRF | Stage III | -0.16992537 | 0.258895 |
| SRF | Stage IV | 0.086391869 | 0.702256 |
| SRF | overall | -0.024796947 | 0.66415 |
| ST14 | G1 | 0.184962406 | 0.434986 |
| ST14 | G2 | 0.061603235 | 0.476182 |
| ST14 | G3 | 0.046596345 | 0.613299 |
| ST14 | GX | -0.191304348 | 0.370533 |
| ST14 | Stage I | 0.102541863 | 0.192736 |
| ST14 | Stage II | 0.049681838 | 0.680742 |
| ST14 | Stage III | 0.013631037 | 0.928359 |
| ST14 | Stage IV | -0.332580463 | 0.130454 |
| ST14 | overall | 0.064128571 | 0.26107 |
| STAT3 | G1 | 0.066165414 | 0.781664 |
| STAT3 | G2 | -0.012117887 | 0.888645 |
| STAT3 | G3 | -0.130437123 | 0.155612 |
| STAT3 | GX | -0.365217391 | 0.07928 |
| STAT3 | Stage I | -0.064559393 | 0.412929 |
| STAT3 | Stage II | 0.167456132 | 0.162762 |
| STAT3 | Stage III | -0.205205701 | 0.171287 |

| | | | |
|---|---|---|---|
| STAT3 | Stage IV | -0.237718803 | 0.286749 |
| STAT3 | overall | -0.060421739 | 0.2897 |
| STAT5A | G1 | -0.097744361 | 0.681832 |
| STAT5A | G2 | -0.015020932 | 0.862207 |
| STAT5A | G3 | -0.080496593 | 0.382126 |
| STAT5A | GX | -0.094782609 | 0.659543 |
| STAT5A | Stage I | -0.051369303 | 0.514903 |
| STAT5A | Stage II | 0.025167884 | 0.834966 |
| STAT5A | Stage III | -0.079750818 | 0.598297 |
| STAT5A | Stage IV | -0.132693394 | 0.556083 |
| STAT5A | overall | -0.032083911 | 0.574223 |
| STAT5B | G1 | 0.04962406 | 0.835414 |
| STAT5B | G2 | -0.094249967 | 0.275088 |
| STAT5B | G3 | -0.024530972 | 0.790277 |
| STAT5B | GX | -0.126086957 | 0.557154 |
| STAT5B | Stage I | -0.075748801 | 0.336535 |
| STAT5B | Stage II | -0.098843887 | 0.412159 |
| STAT5B | Stage III | 0.048047863 | 0.751174 |
| STAT5B | Stage IV | -0.017504235 | 0.938373 |
| STAT5B | overall | -0.056290067 | 0.324005 |
| STK11 | G1 | 0.045112782 | 0.850206 |
| STK11 | G2 | 0.071247451 | 0.409792 |
| STK11 | G3 | 0.040786561 | 0.658271 |
| STK11 | GX | 0.026086957 | 0.903693 |
| STK11 | Stage I | 0.134031817 | 0.088057 |
| STK11 | Stage II | -0.098558841 | 0.413515 |
| STK11 | Stage III | -0.178066985 | 0.236428 |
| STK11 | Stage IV | 0.163184641 | 0.468077 |
| STK11 | overall | 0.039989526 | 0.483688 |
| TBX2 | G1 | -0.009022556 | 0.969885 |
| TBX2 | G2 | -0.108834369 | 0.207218 |
| TBX2 | G3 | -0.071085645 | 0.440393 |
| TBX2 | GX | -0.189565217 | 0.37498 |
| TBX2 | Stage I | -0.063367851 | 0.421623 |
| ATM | Stage II | -0.282195525 | 0.017112 |
| TBX2 | Stage III | -0.014124468 | 0.925772 |
| TBX2 | Stage IV | 0.04121965 | 0.855482 |
| TBX2 | overall | -0.102732535 | 0.071337 |
| TBX3 | G1 | -0.043609023 | 0.855148 |
| TBX3 | G2 | -0.13685579 | 0.112116 |
| TBX3 | G3 | 0.016474367 | 0.858259 |
| TBX3 | GX | 0.082608696 | 0.701163 |
| TBX3 | Stage I | 0.001036364 | 0.989524 |
| TBX3 | Stage II | -0.072770563 | 0.546447 |

| | | | |
|---|---|---|---|
| TBX3 | Stage III | -0.092209956 | 0.542201 |
| TBX3 | Stage IV | -0.003952569 | 0.986072 |
| TBX3 | overall | -0.028036637 | 0.62347 |
| TCF3 | G1 | 0.201503759 | 0.394261 |
| TCF3 | G2 | -0.136068604 | 0.114215 |
| TCF3 | G3 | -0.107732778 | 0.241513 |
| TCF3 | GX | -0.169565217 | 0.428295 |
| TCF3 | Stage I | -0.093907342 | 0.233128 |
| TCF3 | Stage II | -0.137509536 | 0.25281 |
| LOXL3 | Stage II | -0.285683146 | 0.015733 |
| TCF3 | Stage IV | 0.393562959 | 0.069962 |
| ID2 | G2 | -0.289359875 | 6.34E-04 |
| TCF4 | G1 | 0.136842105 | 0.565096 |
| TCF4 | G2 | 0.025676562 | 0.766678 |
| TCF4 | G3 | -0.0173217 | 0.851051 |
| TCF4 | GX | -0.350434783 | 0.093182 |
| TCF4 | Stage I | 0.037397784 | 0.635533 |
| TCF4 | Stage II | -0.119652244 | 0.320277 |
| TCF4 | Stage III | 0.059766854 | 0.693171 |
| TCF4 | Stage IV | 0.04121965 | 0.855482 |
| TCF4 | overall | -6.61E-04 | 0.990773 |
| TCF7 | G1 | -0.082706767 | 0.728851 |
| ID2 | Stage I | -0.291354118 | 1.61E-04 |
| MTA1 | Stage II | -0.292306274 | 0.013377 |
| TCF7 | GX | -0.26 | 0.219832 |
| TCF7 | Stage I | -0.137442951 | 0.080194 |
| ID2 | G3 | -0.292708252 | 0.001178 |
| TCF7 | Stage III | -0.235428362 | 0.115247 |
| TCF7 | Stage IV | -0.255787691 | 0.250573 |
| ID2 | Stage II | -0.298208402 | 0.01154 |
| TEAD1 | G1 | 0.237593985 | 0.313131 |
| TEAD1 | G2 | -0.044590485 | 0.606227 |
| TEAD1 | G3 | -0.057368581 | 0.533694 |
| TEAD1 | GX | -0.195652174 | 0.359548 |
| TEAD1 | Stage I | -0.027103418 | 0.731272 |
| TEAD1 | Stage II | -0.009339448 | 0.938385 |
| TEAD1 | Stage III | -0.097514341 | 0.519114 |
| TEAD1 | Stage IV | -0.17108978 | 0.44649 |
| TEAD1 | overall | -0.045167539 | 0.428851 |
| TFCP2 | G1 | 0.264661654 | 0.259469 |
| TFCP2 | G2 | -0.069253247 | 0.423055 |
| TFCP2 | G3 | -0.019547684 | 0.832175 |
| TFCP2 | GX | -0.213043478 | 0.317545 |
| TFCP2 | Stage I | -0.052300368 | 0.507304 |

| | | | |
|---|---|---|---|
| TFCP2 | Stage II | 0.009993377 | 0.93408 |
| TFCP2 | Stage III | 0.067538396 | 0.655618 |
| TFCP2 | Stage IV | 0.081874647 | 0.717189 |
| TFCP2 | overall | -0.023257434 | 0.683842 |
| TGFA | G1 | -0.172932331 | 0.46594 |
| TGFA | G2 | 0.100072755 | 0.246383 |
| TGFA | G3 | 0.130596866 | 0.155102 |
| TGFA | GX | 0.364347826 | 0.080051 |
| GAB2 | G3 | -0.299282025 | 8.98E-04 |
| TGFA | Stage II | 0.146396264 | 0.223134 |
| TGFA | Stage III | 0.188675755 | 0.209203 |
| TGFA | Stage IV | -0.219649915 | 0.326013 |
| SEMA4C | Stage III | -0.29945106 | 0.043203 |
| TGFB1 | G1 | -0.189473684 | 0.423665 |
| HAS2 | Stage III | -0.303768583 | 0.040137 |
| TGFB1 | G3 | 0.098630898 | 0.28383 |
| TGFB1 | GX | 0.4 | 0.05278 |
| AXIN2 | Stage II | -0.305351319 | 0.009614 |
| MCAM | Stage III | -0.308456179 | 0.037012 |
| TGFB1 | Stage III | -0.108123112 | 0.474457 |
| TGFB1 | Stage IV | -0.159796725 | 0.477489 |
| VIM | Stage III | -0.310429904 | 0.035757 |
| TGFB1I1 | G1 | 0.133834586 | 0.573756 |
| TGFB1I1 | G2 | 0.019526973 | 0.82148 |
| TGFB1I1 | G3 | 0.055139124 | 0.549739 |
| TGFB1I1 | GX | -0.183478261 | 0.390787 |
| TGFB1I1 | Stage I | 0.113661996 | 0.148554 |
| TGFB1I1 | Stage II | -0.064252718 | 0.594488 |
| TGFB1I1 | Stage III | -0.084068341 | 0.578572 |
| TGFB1I1 | Stage IV | 0.120271033 | 0.593941 |
| TGFB1I1 | overall | 0.03592395 | 0.529264 |
| TGFB2 | G1 | 0.245112782 | 0.297601 |
| TGFB2 | G2 | 0.028524743 | 0.741665 |
| TGFB2 | G3 | 0.159642315 | 0.081565 |
| TGFB2 | GX | -0.070434783 | 0.743634 |
| RHOA | G3 | -0.316478709 | 4.29E-04 |
| TGFB2 | Stage II | 0.078404413 | 0.515742 |
| TGFB2 | Stage III | -0.119965461 | 0.427121 |
| TGFB2 | Stage IV | 0.328063241 | 0.136081 |
| TGFB2 | overall | 0.083224437 | 0.144412 |
| TGFB3 | G1 | -0.02556391 | 0.914804 |
| TGFB3 | G2 | -0.025032501 | 0.772369 |
| TGFB3 | G3 | -0.077642054 | 0.399286 |
| TGFB3 | GX | 0.02173913 | 0.919689 |

| | | | |
|---|---|---|---|
| TGFB3 | Stage I | 0.012167581 | 0.877487 |
| TGFB3 | Stage II | -0.078119367 | 0.517275 |
| TGFB3 | Stage III | -0.166101278 | 0.269925 |
| TGFB3 | Stage IV | 0.191417278 | 0.393465 |
| TGFB3 | overall | -0.031417669 | 0.582203 |
| TGFBR1 | G1 | -0.052631579 | 0.825581 |
| TGFBR1 | G2 | -0.017160645 | 0.842816 |
| TGFBR1 | G3 | 0.069561141 | 0.450283 |
| TGFBR1 | GX | 0.173913043 | 0.41637 |
| TGFBR1 | Stage I | 0.077319973 | 0.326579 |
| TGFBR1 | Stage II | 0.016079947 | 0.894118 |
| TGFBR1 | Stage III | 0.089496084 | 0.554201 |
| TGFBR1 | Stage IV | -0.067193676 | 0.766386 |
| TGFBR1 | overall | 0.02739114 | 0.63149 |
| TGFBR3 | G1 | -0.237593985 | 0.313131 |
| TGFBR3 | G2 | 0.046362845 | 0.591975 |
| TGFBR3 | G3 | 0.003677562 | 0.968201 |
| TGFBR3 | GX | -0.16173913 | 0.450219 |
| TGFBR3 | Stage I | 0.001102869 | 0.988852 |
| TGFBR3 | Stage II | -0.099430746 | 0.409375 |
| TGFBR3 | Stage III | 0.10713625 | 0.478523 |
| TGFBR3 | Stage IV | 0.107848673 | 0.632846 |
| TGFBR3 | overall | -0.001006277 | 0.985944 |
| TGM2 | G1 | -0.297744361 | 0.20232 |
| TGM2 | G2 | -0.121858712 | 0.157579 |
| TGM2 | G3 | -0.003865087 | 0.966581 |
| TGM2 | GX | 0.046956522 | 0.827524 |
| TGM2 | Stage I | 0.052416751 | 0.506358 |
| TGM2 | Stage II | -0.166081205 | 0.166291 |
| TGM2 | Stage III | -0.27083205 | 0.068681 |
| TGM2 | Stage IV | 0.017504235 | 0.938373 |
| TGM2 | overall | -0.064706143 | 0.256791 |
| THBD | G1 | -0.009022556 | 0.969885 |
| THBD | G2 | 0.145987143 | 0.08991 |
| MAGED1 | Stage III | -0.317461299 | 0.031569 |
| CDX2 | Stage III | -0.323719191 | 0.028191 |
| TBX2 | Stage II | -0.324164354 | 0.005818 |
| FGFR1 | Stage II | -0.324650609 | 0.005741 |
| THBD | Stage III | 0.191512985 | 0.20231 |
| THBD | Stage IV | -0.201581028 | 0.368337 |
| COL8A2 | Stage III | -0.335224822 | 0.022764 |
| TIAM1 | G1 | 0.040601504 | 0.865047 |
| TIAM1 | G2 | 0.107438904 | 0.213138 |
| TIAM1 | G3 | 0.033174459 | 0.719069 |

| | | | |
|---|---|---|---|
| TIAM1 | GX | -0.169565217 | 0.428295 |
| TIAM1 | Stage I | 0.106011189 | 0.178031 |
| TIAM1 | Stage II | 0.094400523 | 0.433589 |
| TIAM1 | Stage III | 0.135508544 | 0.369227 |
| TIAM1 | Stage IV | -0.343873518 | 0.117117 |
| TIAM1 | overall | 0.05862964 | 0.304269 |
| TIMP1 | G1 | -0.243609023 | 0.300668 |
| TIMP1 | G2 | -0.055081521 | 0.524183 |
| TIMP1 | G3 | -0.10965664 | 0.233162 |
| TIMP1 | GX | -0.022608696 | 0.916487 |
| TIMP1 | Stage I | -0.008720423 | 0.912029 |
| TIMP1 | Stage II | -0.151728301 | 0.206546 |
| TIMP1 | Stage III | -0.124776415 | 0.408679 |
| TIMP1 | Stage IV | -0.193675889 | 0.3878 |
| TIMP1 | overall | -0.076500606 | 0.179831 |
| TM4SF5 | G1 | -0.036510099 | 0.878544 |
| TM4SF5 | G2 | -0.059895049 | 0.48852 |
| TM4SF5 | G3 | -0.065361353 | 0.478162 |
| TM4SF5 | GX | -0.26822536 | 0.205063 |
| TM4SF5 | Stage I | -0.145738796 | 0.063415 |
| TM4SF5 | Stage II | 0.051808783 | 0.667859 |
| TM4SF5 | Stage III | 0.073019942 | 0.629621 |
| TM4SF5 | Stage IV | -0.216815721 | 0.332451 |
| TM4SF5 | overall | -0.086142271 | 0.13081 |
| TMPRSS4 | G1 | -0.001503759 | 0.99498 |
| TMPRSS4 | G2 | 0.144002481 | 0.094412 |
| GAB2 | Stage II | -0.337075261 | 0.004047 |
| TMPRSS4 | GX | -0.133043478 | 0.535424 |
| LOXL2 | Stage III | -0.360636529 | 0.013812 |
| TMPRSS4 | Stage II | 0.08328373 | 0.489876 |
| TNF | Stage III | -0.376796401 | 0.009848 |
| TMPRSS4 | Stage IV | -0.121400339 | 0.590455 |
| TCF3 | Stage III | -0.376919759 | 0.009822 |
| TNC | G1 | -0.105263158 | 0.658728 |
| TNC | G2 | 0.111026562 | 0.19816 |
| TNC | G3 | -0.024430264 | 0.791118 |
| TNC | GX | -0.085217391 | 0.692169 |
| TWIST1 | Stage III | -0.405045337 | 0.005233 |
| TNC | Stage II | -0.008668751 | 0.942802 |
| TNC | Stage III | -0.2882255 | 0.052078 |
| TNC | Stage IV | -0.284020327 | 0.200198 |
| TNC | overall | 0.026156478 | 0.64695 |
| TNF | G1 | -0.019548872 | 0.934803 |
| TNF | G2 | -0.041324857 | 0.632876 |

| | | | |
|---|---|---|---|
| TNF | G3 | 0.054833529 | 0.551956 |
| TNF | GX | 0.066956522 | 0.75591 |
| TNF | Stage I | 0.120880522 | 0.124278 |
| TNF | Stage II | -0.046663704 | 0.699181 |
| RHOA | GX | -0.407826087 | 0.047901 |
| TNF | Stage IV | 0.139469226 | 0.535903 |
| TNF | overall | 0.014293885 | 0.802388 |
| TP53 | G1 | 0.418045113 | 0.066622 |
| TP53 | G2 | -0.018329497 | 0.832262 |
| TP53 | G3 | -0.055194687 | 0.549336 |
| TP53 | GX | -0.3 | 0.154363 |
| TP53 | Stage I | -0.040232545 | 0.610116 |
| TP53 | Stage II | 0.010982654 | 0.927571 |
| TP53 | Stage III | -0.181274287 | 0.227954 |
| TP53 | Stage IV | 0.215132693 | 0.336309 |
| TP53 | overall | -0.029165343 | 0.609554 |
| TP73 | G1 | 0.413533835 | 0.069921 |
| SHH | GX | -0.407864151 | 0.047878 |
| TP73 | G3 | 0.012234232 | 0.894493 |
| TP73 | GX | 0.174782609 | 0.414007 |
| PROM1 | Stage II | -0.409574513 | 3.90E-04 |
| RAF1 | GX | -0.420869565 | 0.04056 |
| TP73 | Stage III | -0.179547278 | 0.232491 |
| TP73 | Stage IV | 0.075098814 | 0.739774 |
| FHL2 | Stage III | -0.421451925 | 0.003534 |
| TRPS1 | G1 | 0.040601504 | 0.865047 |
| TRPS1 | G2 | -0.048972484 | 0.571271 |
| KHDRBS1 | GX | -0.431304348 | 0.03535 |
| TRPS1 | GX | -0.393043478 | 0.057433 |
| RNF111 | GX | -0.431304348 | 0.03535 |
| TRPS1 | Stage II | -0.220608825 | 0.064496 |
| TRPS1 | Stage III | -0.043360267 | 0.774782 |
| TRPS1 | Stage IV | 0.217391304 | 0.331137 |
| CDH1 | GX | -0.435652174 | 0.033343 |
| TWIST1 | G1 | -0.121804511 | 0.608957 |
| TWIST1 | G2 | -0.064181864 | 0.457881 |
| TWIST1 | G3 | 0.123175761 | 0.180145 |
| TWIST1 | GX | 0.30173913 | 0.151871 |
| TWIST1 | Stage I | 0.144035225 | 0.066603 |
| TWIST1 | Stage II | 0.010664073 | 0.929666 |
| ZYX | Stage III | -0.435884787 | 0.002462 |
| TWIST1 | Stage IV | 0.166572558 | 0.458761 |
| TWIST1 | overall | 0.026392998 | 0.643976 |
| TXNIP | G1 | -0.058646617 | 0.805993 |

| | | | |
|---|---|---|---|
| TXNIP | G2 | 0.058795606 | 0.49655 |
| TXNIP | G3 | 0.139476495 | 0.128679 |
| TXNIP | GX | 0.053043478 | 0.805559 |
| IL6R | Stage IV | -0.436476567 | 0.042264 |
| TXNIP | Stage II | -0.079410458 | 0.510353 |
| TXNIP | Stage III | 0.151668415 | 0.314323 |
| TXNIP | Stage IV | -0.108977979 | 0.629268 |
| TXNIP | overall | 0.082441868 | 0.148238 |
| VCAN | G1 | -0.016541353 | 0.944817 |
| VCAN | G2 | 0.013205634 | 0.878723 |
| VCAN | G3 | 0.02912532 | 0.75217 |
| VCAN | GX | -0.242608696 | 0.253348 |
| VCAN | Stage I | 0.054752727 | 0.487571 |
| VCAN | Stage II | -0.069752429 | 0.563253 |
| VCAN | Stage III | -0.120828965 | 0.423777 |
| VCAN | Stage IV | 0.129305477 | 0.566299 |
| VCAN | overall | 0.003637686 | 0.94922 |
| VDR | G1 | 0.126315789 | 0.595654 |
| VDR | G2 | 0.09902556 | 0.251386 |
| VDR | G3 | 0.146442679 | 0.110483 |
| VDR | GX | 0.096521739 | 0.653673 |
| CLDN4 | GX | -0.437391304 | 0.032566 |
| NFIC | Stage III | -0.437488439 | 0.002363 |
| VDR | Stage III | 0.085671992 | 0.571321 |
| ELF5 | GX | -0.44173913 | 0.030687 |
| PARP1 | GX | -0.447826087 | 0.028201 |
| VEGFA | G1 | 0.216541353 | 0.359146 |
| VEGFA | G2 | -0.129861765 | 0.13185 |
| VEGFA | G3 | -0.101287494 | 0.270997 |
| PTN | GX | -0.447826087 | 0.028201 |
| VEGFA | Stage I | -0.110774585 | 0.159212 |
| VEGFA | Stage II | -0.218915316 | 0.066626 |
| VEGFA | Stage III | -0.069758836 | 0.645036 |
| VEGFA | Stage IV | -0.169960474 | 0.449542 |
| ZNF217 | GX | -0.449565217 | 0.027522 |
| VIM | G1 | -0.010526316 | 0.964869 |
| VIM | G2 | -0.041627804 | 0.630383 |
| VIM | G3 | -0.043245214 | 0.639078 |
| VIM | GX | -0.085217391 | 0.692169 |
| VIM | Stage I | 0.051710139 | 0.512114 |
| VIM | Stage II | -0.187392584 | 0.117614 |
| ZBTB33 | Stage IV | -0.453416149 | 0.034061 |
| VIM | Stage IV | 0.280632411 | 0.205851 |
| VIM | overall | -0.051011183 | 0.371513 |

| | | | |
|---|---|---|---|
| VSNL1 | G1 | 0.022556391 | 0.924798 |
| PROM1 | GX | -0.453913043 | 0.02588 |
| ITGA6 | Stage IV | -0.454545455 | 0.033563 |
| ID2 | GX | -0.474782609 | 0.01906 |
| VDR | Stage IV | -0.477131564 | 0.024744 |
| BCL2 | GX | -0.47826087 | 0.018081 |
| VSNL1 | Stage III | 0.180534141 | 0.229891 |
| VSNL1 | Stage IV | -0.238848108 | 0.284397 |
| LATS1 | GX | -0.480869565 | 0.017373 |
| VTN | G1 | -0.287218045 | 0.219504 |
| VTN | G2 | -0.055048191 | 0.524435 |
| VTN | G3 | -0.106013804 | 0.249149 |
| VTN | GX | -0.004347826 | 0.983913 |
| VTN | Stage I | -0.055345727 | 0.482862 |
| VTN | Stage II | -0.031170617 | 0.796369 |
| VTN | Stage III | -0.200394746 | 0.181768 |
| VTN | Stage IV | -0.263692829 | 0.235716 |
| VTN | overall | -0.095299562 | 0.094479 |
| WISP2 | G1 | -0.114285714 | 0.631392 |
| WISP2 | G2 | 0.129902317 | 0.131729 |
| WISP2 | G3 | -0.064897339 | 0.481298 |
| WISP2 | GX | 0.166956522 | 0.435538 |
| WISP2 | Stage I | 0.131548977 | 0.094158 |
| WISP2 | Stage II | -0.093461548 | 0.438197 |
| WISP2 | Stage III | -0.021032505 | 0.889657 |
| WISP2 | Stage IV | 0.242236025 | 0.277415 |
| WISP2 | overall | 0.040939266 | 0.473355 |
| WNT3A | G1 | 0.085714286 | 0.719366 |
| CTNNBIP1 | Stage IV | -0.487295313 | 0.021434 |
| NF1 | GX | -0.519130435 | 0.009336 |
| WNT3A | GX | 0.29826087 | 0.156883 |
| VEGFA | GX | -0.52 | 0.009198 |
| SON | GX | -0.520869565 | 0.009061 |
| WNT3A | Stage III | 0.234071426 | 0.117419 |
| ERBB2 | GX | -0.526086957 | 0.008276 |
| BRAF | GX | -0.537391304 | 0.006767 |
| WNT5A | G1 | -0.2 | 0.397873 |
| WNT5A | G2 | 0.014904047 | 0.863269 |
| WNT5A | G3 | 0.059726527 | 0.516984 |
| WNT5A | GX | -0.393043478 | 0.057433 |
| WNT5A | Stage I | 0.055808489 | 0.479204 |
| WNT5A | Stage II | -0.112894977 | 0.34856 |
| WNT5A | Stage III | 0.021402578 | 0.887727 |
| WNT5A | Stage IV | 0.01863354 | 0.934405 |

| | | | |
|---|---|---|---|
| WNT5A | overall | 0.013099491 | 0.818601 |
| WWTR1 | G1 | 0.356390977 | 0.122986 |
| WWTR1 | G2 | 0.027148361 | 0.753721 |
| WWTR1 | G3 | 0.017380735 | 0.850549 |
| WWTR1 | GX | -0.362608696 | 0.081612 |
| WWTR1 | Stage I | 0.08165386 | 0.300111 |
| WWTR1 | Stage II | -0.114487881 | 0.341757 |
| WWTR1 | Stage III | -0.04878801 | 0.747467 |
| WWTR1 | Stage IV | 0.238848108 | 0.284397 |
| WWTR1 | overall | 0.021063678 | 0.712272 |
| YAP1 | G1 | 0.120300752 | 0.613418 |
| YAP1 | G2 | 0.025209022 | 0.770808 |
| YAP1 | G3 | 0.175370057 | 0.055384 |
| YAP1 | GX | 0.017391304 | 0.935715 |
| KRT19 | Stage IV | -0.551665726 | 0.007777 |
| YAP1 | Stage II | 0.095272429 | 0.429335 |
| YAP1 | Stage III | 0.071855919 | 0.635106 |
| YAP1 | Stage IV | -0.140598532 | 0.532573 |
| YAP1 | overall | 0.097217741 | 0.087999 |
| YBX1 | G1 | -0.013533835 | 0.95484 |
| YBX1 | G2 | 0.019822764 | 0.818822 |
| YBX1 | G3 | -0.009876286 | 0.914742 |
| BMI1 | GX | -0.56173913 | 0.004283 |
| YBX1 | Stage I | -0.004012447 | 0.959458 |
| YBX1 | Stage II | 0.056422338 | 0.640245 |
| YBX1 | Stage III | -0.121445754 | 0.421397 |
| YBX1 | Stage IV | 0.185770751 | 0.407828 |
| YBX1 | overall | 0.019183519 | 0.736962 |
| YWHAZ | G1 | 0.284210526 | 0.224585 |
| YWHAZ | G2 | 0.123270875 | 0.152785 |
| YWHAZ | G3 | 0.074763207 | 0.417047 |
| YWHAZ | GX | 0.129565217 | 0.546239 |
| YWHAZ | Stage I | 0.065529252 | 0.40593 |
| PLAUR | G1 | -0.577443609 | 0.007673 |
| MAPK1 | GX | -0.612173913 | 0.001476 |
| YWHAZ | Stage IV | -0.328063241 | 0.136081 |
| LRP6 | GX | -0.633043478 | 9.00E-04 |
| YY1 | G1 | 0.273684211 | 0.242966 |
| YY1 | G2 | 0.015948857 | 0.853787 |
| YY1 | G3 | 0.021148587 | 0.818658 |
| YY1 | GX | 0.047826087 | 0.824378 |
| YY1 | Stage I | -0.029746978 | 0.706215 |
| YY1 | Stage II | 0.001626439 | 0.98926 |
| YY1 | Stage III | 0.10824647 | 0.47395 |

| | | | |
|---|---|---|---|
| YY1 | Stage IV | 0.043478261 | 0.84765 |
| YY1 | overall | -0.004298029 | 0.940019 |
| ZBTB33 | G1 | 0.160902256 | 0.497974 |
| ZBTB33 | G2 | 0.056014217 | 0.517171 |
| ZBTB33 | G3 | 0.096300734 | 0.295416 |
| ZBTB33 | GX | -0.142608696 | 0.506209 |
| ZBTB33 | Stage I | 0.074180399 | 0.346663 |
| ZBTB33 | Stage II | 0.087089932 | 0.470183 |
| ZBTB33 | Stage III | 0.255412325 | 0.086683 |
| HDAC6 | GX | -0.64 | 7.57E-04 |
| ZBTB33 | overall | 0.072249556 | 0.20532 |
| ZEB1 | G1 | -0.045112782 | 0.850206 |
| ZEB1 | G2 | -0.061932421 | 0.473824 |
| ZEB1 | G3 | -0.056917133 | 0.536924 |
| ZEB1 | GX | -0.191304348 | 0.370533 |
| ZEB1 | Stage I | -0.0094686 | 0.904515 |
| ZEB1 | Stage II | -0.104695713 | 0.384894 |
| ZEB1 | Stage III | -0.091346451 | 0.546006 |
| ZEB1 | Stage IV | 0.050254094 | 0.824241 |
| ZEB1 | overall | -0.062960208 | 0.269876 |
| ZFYVE9 | G1 | 0.039097744 | 0.870004 |
| ZFYVE9 | G2 | 0.165301814 | 0.054456 |
| ZFYVE9 | G3 | 0.042248557 | 0.64683 |
| ZFYVE9 | GX | 0.223478261 | 0.293847 |
| ZFYVE9 | Stage I | 0.081055318 | 0.30368 |
| ZFYVE9 | Stage II | 0.094920313 | 0.43105 |
| ZFYVE9 | Stage III | 0.214334178 | 0.152613 |
| ZFYVE9 | Stage IV | 0.239977414 | 0.282058 |
| ZFYVE9 | overall | 0.107419815 | 0.059285 |
| ZNF217 | G1 | 0.172932331 | 0.46594 |
| ZNF217 | G2 | -0.008787853 | 0.919123 |
| ZNF217 | G3 | -0.054826583 | 0.552006 |
| OCLN | GX | -0.66173913 | 4.29E-04 |
| ZNF217 | Stage I | -0.129296131 | 0.09998 |
| ZNF217 | Stage II | -0.048575189 | 0.687482 |
| ZNF217 | Stage III | 0.210510086 | 0.160245 |
| ZNF217 | Stage IV | 0.00621118 | 0.978115 |
| ZNF217 | overall | -0.04074403 | 0.475469 |
| ZYX | G1 | 0.266165414 | 0.256671 |
| ZYX | G2 | -0.064336915 | 0.456794 |
| ZYX | G3 | -0.029344099 | 0.750369 |
| ZYX | GX | 0.064347826 | 0.765154 |
| ZYX | Stage I | 0.01985441 | 0.801383 |
| ZYX | Stage II | 0.076191115 | 0.5277 |

| | | | |
|---|---|---|---|
| MYCN | GX | -0.691304348 | 1.83E-04 |
| ZYX | Stage IV | 0.328063241 | 0.136081 |
| ZYX | overall | -0.031633852 | 0.579608 |

#### Supplementary File 3: EMT Scores for the TCGA-CESC

| Sample_ID | EMT z score | CDH1 z.score | DSP z.score | TJP1 z.score | VIM z.score | FN1 z.score | CDH2 z.score | ITBG6 z.score | **FOXC2_z.score** | **MMP2_z.score** | **SNAI1_z.score** | **MMP3_z.score** | **SNAI2_z.score** | **MMP9_z.score** | **TWIST1_z.score** | **SOX10_z.score** | **GSC_z.score** |
|---|---|---|---|---|---|---|---|---|---|---|---|---|---|---|---|---|---|
| TCGA-VS-A9V2-01A-11R | 120.474 | -0.834 | 2.314 | -0.394 | -0.307 | -0.441 | -0.287 | 0.000 | -0.504 | -0.426 | -0.317 | -0.306 | 0.702 | 0.046 | -0.297 | NA | NA |
| TCGA-C5-A8ZZ-01A-11R | 123.075 | 2.922 | 1.886 | 0.670 | 0.210 | 1.757 | 0.110 | 0.000 | -0.471 | 2.087 | -0.299 | -0.301 | 0.855 | 0.023 | -0.019 | NA | NA |
| TCGA-FU-A3HZ-01A-11R | 26.444 | 0.043 | -0.509 | -0.918 | 0.582 | 0.089 | 3.669 | 0.000 | -0.079 | -0.571 | -0.035 | -0.335 | -1.012 | -0.414 | 1.897 | NA | NA |
| TCGA-C5-A1MK-01A-11R | 39.706 | 0.067 | 0.064 | 0.684 | -0.110 | 0.615 | -0.184 | 0.000 | -0.034 | 1.894 | 0.244 | 0.054 | -0.219 | -0.207 | 0.027 | NA | NA |
| TCGA-VS-A94Z-01A-11R | 41.528 | -0.189 | -0.410 | 0.020 | -0.254 | -0.294 | -0.284 | 0.000 | 0.117 | -0.435 | -0.321 | -0.270 | 0.688 | -0.437 | -0.004 | NA | NA |
| TCGA-MU-A51Y-01A-11R | 59.318 | 0.108 | 0.280 | 1.342 | 0.126 | -0.067 | -0.280 | 0.000 | -0.414 | 0.415 | -0.320 | -0.287 | -0.195 | -0.096 | -0.158 | NA | NA |
| TCGA-C5-A7X8-01A-11R | -18.206 | -0.409 | -1.180 | -0.612 | 1.436 | -0.291 | 0.598 | 0.000 | 1.611 | 2.006 | 0.277 | -0.315 | -0.223 | -0.420 | -0.025 | NA | NA |
| TCGA-MU-A5YI-01A-11R | 20.409 | -0.741 | -0.156 | 0.128 | 0.614 | 0.279 | -0.185 | 0.000 | -0.192 | 1.287 | -0.288 | -0.283 | 0.407 | -0.335 | -0.112 | NA | NA |
| TCGA-VS-A9V5-01A-11R | 16.201 | -1.084 | -1.046 | 0.968 | -0.480 | -0.419 | -0.257 | 0.000 | 0.214 | -0.360 | -0.105 | -0.311 | -1.129 | -0.465 | -0.293 | NA | NA |
| TCGA-FU-A3TQ-01A-11R | 101.723 | 0.022 | 1.427 | 2.500 | -0.338 | -0.234 | -0.226 | 0.000 | -0.499 | -0.099 | -0.251 | 0.064 | -0.440 | 0.414 | -0.195 | NA | NA |
| TCGA-LP-A4AX-01A-12R | -34.401 | -1.390 | -0.153 | -0.672 | 0.721 | 5.970 | 0.157 | 0.000 | 1.089 | -0.072 | 3.683 | 0.477 | -0.289 | 0.100 | 0.322 | NA | NA |
| TCGA-C5-A8YR-01A-12R | 6.570 | -0.602 | -0.595 | -1.149 | -0.105 | 1.037 | 0.870 | 0.000 | 1.273 | -0.719 | 15.464 | -0.205 | -1.158 | -0.195 | -0.223 | NA | NA |
| TCGA-VS-A959-01A-11R | 56.595 | 1.424 | -0.515 | 0.236 | -0.243 | 0.070 | -0.252 | 0.000 | -0.508 | 0.033 | 0.348 | -0.303 | -0.980 | -0.371 | -0.243 | NA | NA |
| TCGA-C5-A907-01A-11R | 36.779 | -0.477 | -0.570 | 0.659 | -0.435 | -0.292 | 0.225 | 0.000 | 0.555 | -0.521 | -0.198 | -0.267 | -0.381 | -0.416 | -0.254 | NA | NA |
| TCGA-EK-A2H0-01A-11R | 92.643 | -0.369 | 1.104 | 0.007 | -0.566 | -0.520 | -0.276 | 0.000 | 0.734 | -0.649 | -0.296 | -0.284 | 1.473 | -0.279 | -0.190 | NA | NA |
| TCGA-MA-AA3X-01A-22R | 123.487 | 0.478 | 2.193 | 0.016 | -0.021 | 0.112 | -0.281 | 0.000 | 3.862 | -0.468 | -0.102 | -0.321 | -0.520 | 0.677 | -0.294 | NA | NA |
| TCGA-Q1-A73P-01A-11R | 33.584 | 0.483 | -0.936 | -0.339 | -0.289 | -0.465 | -0.217 | 0.000 | 2.969 | -0.064 | -0.271 | -0.313 | -1.108 | -0.315 | -0.281 | NA | NA |

| | | | | | | | | | | | | | | | | | |
|---|---|---|---|---|---|---|---|---|---|---|---|---|---|---|---|---|---|
| TCGA-EK-A2RK-01A-11R | 70.032 | -0.730 | 0.723 | -1.692 | -0.414 | -0.496 | -0.261 | 0.000 | -0.478 | -0.756 | -0.321 | -0.334 | 3.186 | -0.360 | 0.206 | NA | NA |
| TCGA-VS-A8EB-01A-11R | 49.158 | -0.493 | -0.017 | -0.686 | -0.511 | -0.389 | -0.284 | 0.000 | 0.022 | -0.440 | -0.277 | 0.057 | 1.635 | -0.331 | 0.363 | NA | NA |
| TCGA-Q1-A73O-01A-11R | 85.483 | 0.315 | 0.870 | 0.057 | -0.443 | -0.439 | -0.268 | 0.000 | -0.487 | -0.260 | -0.154 | 1.179 | 1.208 | -0.372 | -0.253 | NA | NA |
| TCGA-C5-A7X3-01A-11R | 23.508 | 0.680 | -0.779 | 0.487 | -0.216 | 0.692 | -0.120 | 0.000 | 2.434 | 0.694 | 0.066 | 0.572 | -0.185 | -0.167 | -0.230 | NA | NA |
| TCGA-BI-A0VR-01A-11R | 98.483 | -0.488 | 1.656 | 0.101 | -0.132 | -0.130 | -0.261 | 0.000 | -0.289 | -0.145 | 1.135 | -0.280 | 0.097 | -0.021 | -0.087 | NA | NA |
| TCGA-LP-A7HU-01A-11R | 9.835 | -0.901 | -1.060 | -0.952 | -0.191 | -0.285 | -0.293 | 0.000 | -0.498 | -0.335 | -0.228 | -0.286 | -1.123 | -0.280 | 0.020 | NA | NA |
| TCGA-Q1-A6DV-01A-11R | 50.396 | 0.670 | -0.715 | 1.172 | -0.445 | -0.509 | -0.012 | 0.000 | 1.670 | -0.508 | -0.337 | -0.280 | -1.177 | -0.380 | -0.256 | NA | NA |
| TCGA-HG-A2PA-01A-11R | 25.855 | -0.244 | -0.662 | 0.550 | -0.096 | -0.294 | -0.276 | 0.000 | -0.373 | 0.594 | 0.131 | -0.044 | 0.045 | -0.331 | -0.188 | NA | NA |
| TCGA-DG-A2KL-01A-11R | 19.083 | 0.054 | 0.557 | 2.913 | 0.381 | 3.166 | 0.095 | 0.000 | -0.025 | 3.883 | -0.161 | -0.220 | 0.328 | 0.718 | 0.652 | NA | NA |
| TCGA-JW-A5VI-01A-11R | 65.178 | -0.095 | 0.405 | 2.629 | -0.115 | -0.145 | -0.226 | 0.000 | -0.175 | 0.321 | -0.285 | -0.080 | 0.374 | -0.190 | -0.244 | NA | NA |
| TCGA-Q1-A73S-01A-11R | 95.246 | 2.459 | 0.068 | -0.472 | -0.298 | -0.332 | -0.201 | 0.000 | -0.510 | -0.598 | -0.214 | -0.281 | -1.111 | -0.360 | -0.207 | NA | NA |
| TCGA-FU-A57G-01A-11R | -16.705 | 1.672 | -0.980 | 0.040 | 3.042 | 1.857 | 2.507 | 0.000 | -0.449 | 1.593 | 0.293 | -0.231 | -0.543 | -0.146 | -0.074 | NA | NA |
| TCGA-EK-A3GN-01A-11R | 23.993 | -1.315 | -0.682 | -0.942 | -0.634 | -0.485 | -0.282 | 0.000 | 0.937 | -0.661 | -0.017 | -0.325 | 0.141 | -0.466 | -0.319 | NA | NA |
| TCGA-EA-A50E-01A-21R | 42.915 | -0.883 | 0.837 | 0.238 | 0.531 | 1.296 | -0.161 | 0.000 | -0.330 | 0.751 | 0.234 | -0.298 | 0.804 | 0.186 | 0.024 | NA | NA |
| TCGA-C5-A3HF-01A-11R | 20.651 | -0.317 | -0.832 | 0.050 | -0.080 | -0.439 | -0.032 | 0.000 | -0.233 | 0.690 | -0.201 | -0.226 | -0.957 | -0.248 | -0.225 | NA | NA |
| TCGA-VS-A9UH-01A-11R | 0.931 | -0.631 | -0.646 | -1.434 | 0.482 | -0.070 | -0.056 | 0.000 | -0.139 | 0.375 | 0.019 | 0.205 | 0.920 | 0.898 | 0.035 | NA | NA |
| TCGA-EK-A2RM-01A-21R | -20.650 | 0.540 | -1.172 | -0.954 | -0.076 | 0.468 | 0.228 | 0.000 | 0.336 | 1.806 | 0.614 | 6.245 | -0.870 | -0.310 | 0.115 | NA | NA |
| TCGA-VS-A958-01A-11R | 45.392 | -0.055 | -0.399 | 0.621 | -0.218 | -0.260 | -0.303 | 0.000 | -0.507 | -0.661 | 0.061 | -0.001 | -0.975 | -0.290 | -0.312 | NA | NA |
| TCGA-EA-A3HT-01A-61R | 7.419 | -0.265 | -0.078 | 0.801 | 0.102 | 3.036 | 0.071 | 0.000 | 0.205 | 2.498 | -0.135 | -0.146 | 0.699 | -0.226 | 0.247 | NA | NA |
| TCGA-IR-A3LF-01A-21R | 21.018 | -0.292 | -0.959 | 0.743 | -0.099 | -0.393 | -0.211 | 0.000 | -0.503 | 0.012 | -0.214 | -0.315 | -1.035 | -0.141 | -0.223 | NA | NA |

| | | | | | | | | | | | | | | | | | |
|---|---|---|---|---|---|---|---|---|---|---|---|---|---|---|---|---|---|
| TCGA-VS-A9UY-01A-11R | 50.539 | -0.253 | -0.148 | 0.969 | -0.416 | -0.425 | -0.270 | 0.000 | -0.498 | -0.684 | -0.093 | 0.000 | 0.781 | 0.045 | -0.298 | NA | NA |
| TCGA-VS-A954-01A-11R | 82.208 | -1.528 | 1.675 | -0.824 | 0.090 | -0.161 | -0.269 | 0.000 | -0.498 | -0.416 | -0.292 | -0.326 | 3.202 | -0.356 | 0.067 | NA | NA |
| TCGA-ZJ-AAXI-01A-11R | 48.699 | -0.691 | -0.166 | -0.075 | -0.543 | -0.524 | -0.304 | 0.000 | -0.347 | -0.757 | -0.364 | -0.332 | 1.597 | -0.439 | -0.299 | NA | NA |
| TCGA-FU-A3TX-01A-11R | 26.898 | -0.914 | -0.396 | -0.590 | -0.318 | -0.016 | -0.187 | 0.000 | 1.226 | -0.741 | 0.004 | -0.331 | -0.285 | -0.010 | 2.859 | NA | NA |
| TCGA-EK-A2RN-01A-12R | 37.483 | -0.457 | -0.642 | -0.403 | -0.536 | -0.521 | -0.294 | 0.000 | -0.498 | -0.787 | -0.258 | -0.321 | 0.415 | -0.460 | -0.331 | NA | NA |
| TCGA-DG-A2KJ-01A-11R | -1.454 | -1.131 | -1.226 | -2.037 | -0.391 | -0.521 | -0.218 | 0.000 | -0.519 | -0.576 | -0.351 | -0.171 | -1.173 | 0.701 | -0.281 | NA | NA |
| TCGA-JX-A3PZ-01A-11R | -2.159 | -0.234 | -0.742 | -0.256 | -0.055 | 0.479 | -0.149 | 0.000 | 0.417 | 0.081 | 1.641 | 0.477 | -0.386 | 2.431 | -0.208 | NA | NA |
| TCGA-HM-A6W2-01A-21R | -123.318 | -0.546 | -1.202 | -1.607 | 11.152 | -0.365 | 0.220 | 0.000 | -0.501 | 0.232 | -0.109 | -0.331 | -0.210 | -0.437 | 4.483 | NA | NA |
| TCGA-FU-A3NI-01A-11R | 95.601 | -0.164 | 1.263 | 0.801 | -0.381 | -0.363 | -0.265 | 0.000 | 1.478 | 0.407 | 0.103 | -0.321 | -0.081 | -0.357 | -0.261 | NA | NA |
| TCGA-ZJ-AAXD-01A-21R | 52.478 | 0.742 | -0.010 | -0.786 | 0.082 | 0.056 | -0.142 | 0.000 | 0.935 | 0.657 | -0.207 | -0.192 | 1.097 | -0.391 | 0.047 | NA | NA |
| TCGA-JX-A5QV-01A-22R | 59.768 | -0.361 | 0.263 | 0.988 | -0.291 | -0.506 | -0.287 | 0.000 | -0.362 | -0.631 | -0.307 | -0.212 | 2.153 | 0.027 | 0.731 | NA | NA |
| TCGA-C5-A7CO-01A-11R | 28.314 | -0.992 | -0.657 | -0.979 | -0.521 | -0.515 | -0.297 | 0.000 | -0.083 | -0.678 | -0.359 | -0.331 | -0.092 | -0.424 | 0.131 | NA | NA |
| TCGA-PN-A8MA-01A-11R | 91.436 | 4.159 | -0.609 | -0.118 | -0.177 | -0.147 | 0.760 | 0.000 | 1.028 | -0.330 | -0.182 | -0.322 | -0.823 | -0.455 | -0.219 | NA | NA |
| TCGA-DS-A1OD-01A-11R | 56.276 | 1.447 | -0.541 | -0.011 | 0.037 | -0.395 | -0.239 | 0.000 | -0.335 | -0.245 | -0.019 | -0.326 | -0.022 | -0.367 | -0.201 | NA | NA |
| TCGA-C5-A1ME-01A-11R | 10.218 | -0.899 | -1.150 | 0.545 | -0.060 | -0.528 | -0.251 | 0.000 | 1.436 | -0.600 | -0.279 | -0.300 | -1.144 | -0.250 | -0.254 | NA | NA |
| TCGA-C5-A3HD-01B-11R | 69.080 | -0.728 | 0.855 | -0.880 | -0.062 | -0.242 | -0.194 | 0.000 | -0.279 | -0.522 | -0.356 | 0.123 | 1.326 | -0.332 | 0.147 | NA | NA |
| TCGA-EA-A6QX-01A-12R | 18.445 | -0.730 | -0.890 | -0.564 | -0.325 | -0.392 | -0.279 | 0.000 | -0.488 | 0.034 | -0.279 | -0.277 | -0.426 | -0.353 | -0.228 | NA | NA |
| TCGA-IR-A3LC-01A-11R | 57.388 | -0.842 | 0.299 | -0.082 | -0.385 | -0.503 | -0.259 | 0.000 | -0.514 | -0.266 | -0.033 | -0.320 | 0.346 | -0.194 | -0.283 | NA | NA |
| TCGA-VS-A950-01A-11R | 121.564 | 1.152 | 1.796 | 0.032 | -0.357 | 0.203 | -0.241 | 0.000 | 6.809 | -0.367 | -0.085 | 0.046 | 0.302 | 0.174 | -0.277 | NA | NA |
| TCGA-C5-A8YT-01A-11R | -48.112 | -1.047 | -1.089 | 0.653 | 2.571 | -0.160 | 1.767 | 0.000 | -0.506 | 4.452 | 0.588 | -0.314 | -0.233 | -0.391 | 0.451 | NA | NA |

| | | | | | | | | | | | | | | | | | |
|---|---|---|---|---|---|---|---|---|---|---|---|---|---|---|---|---|---|
| TCGA-EK-A2PL-01A-11R | 96.270 | 1.547 | 0.417 | 1.172 | -0.623 | -0.507 | -0.293 | 0.000 | -0.414 | 0.370 | -0.360 | -0.304 | 0.160 | -0.440 | -0.306 | NA | NA |
| TCGA-Q1-A73Q-01A-21R | 91.901 | 0.404 | 0.770 | 0.219 | -0.493 | -0.492 | -0.296 | 0.000 | -0.108 | -0.560 | -0.309 | -0.217 | 0.477 | -0.349 | 0.512 | NA | NA |
| TCGA-Q1-A73R-01A-11R | 19.041 | 1.111 | -0.919 | 2.605 | 1.946 | -0.283 | -0.262 | 0.000 | -0.483 | 0.710 | -0.255 | -0.297 | -1.128 | -0.453 | -0.295 | NA | NA |
| TCGA-RA-A741-01A-11R | 23.164 | -0.023 | -0.981 | -0.905 | -0.356 | -0.505 | -0.197 | 0.000 | -0.380 | -0.006 | -0.291 | 0.153 | -0.440 | -0.243 | -0.267 | NA | NA |
| TCGA-C5-A2LZ-01A-11R | 84.778 | -0.464 | 0.796 | 1.155 | -0.503 | -0.500 | -0.285 | 0.000 | -0.496 | -0.688 | -0.319 | -0.290 | 0.211 | -0.306 | -0.287 | NA | NA |
| TCGA-UC-A7PD-01A-11R | 44.137 | -0.082 | 0.816 | 0.391 | -0.049 | -0.059 | -0.207 | 0.000 | 0.273 | -0.612 | 0.027 | 8.241 | -0.030 | -0.206 | -0.073 | NA | NA |
| TCGA-JX-A3Q8-01A-11R | 13.047 | -0.512 | -0.987 | -1.262 | 0.524 | -0.534 | -0.302 | 0.000 | 0.504 | -0.617 | -0.125 | -0.324 | -1.166 | -0.397 | -0.259 | NA | NA |
| TCGA-LP-A4AU-01A-32R | 96.790 | -0.406 | 1.316 | 0.120 | -0.267 | -0.328 | -0.262 | 0.000 | -0.026 | -0.455 | 0.028 | -0.263 | -0.399 | -0.386 | -0.274 | NA | NA |
| TCGA-IR-A3LL-01A-11R | 20.228 | -1.228 | -0.614 | -0.686 | -0.211 | -0.427 | -0.289 | 0.000 | -0.512 | -0.686 | -0.340 | -0.244 | 0.157 | -0.029 | -0.227 | NA | NA |
| TCGA-EA-A3QE-01A-21R | 73.483 | 1.044 | 0.030 | 0.918 | -0.212 | -0.405 | -0.282 | 0.000 | -0.479 | -0.323 | -0.326 | -0.236 | -0.182 | -0.242 | -0.229 | NA | NA |
| TCGA-VS-A8Q9-01A-12R | 60.211 | -1.084 | 0.374 | -0.925 | -0.545 | -0.465 | -0.306 | 0.000 | 4.797 | -0.743 | -0.261 | -0.333 | -0.498 | -0.404 | -0.316 | NA | NA |
| TCGA-C5-A8XJ-01A-11R | 84.360 | 2.173 | 0.392 | 0.199 | -0.100 | 0.401 | 0.089 | 0.000 | 0.113 | 1.044 | -0.207 | -0.236 | -0.181 | -0.349 | 1.006 | NA | NA |
| TCGA-VS-A9UC-01A-11R | 66.966 | 0.822 | 0.121 | 0.249 | -0.272 | -0.176 | -0.186 | 0.000 | -0.512 | 0.646 | 0.468 | 0.129 | -1.054 | -0.405 | -0.293 | NA | NA |
| TCGA-LP-A5U2-01A-11R | 38.969 | -0.787 | -0.514 | 0.141 | -0.538 | -0.493 | -0.286 | 0.000 | -0.513 | -0.710 | 0.048 | -0.303 | -0.882 | -0.466 | -0.275 | NA | NA |
| TCGA-JW-A5VJ-01A-11R | 68.577 | 0.431 | -0.026 | 0.441 | -0.533 | -0.435 | -0.280 | 0.000 | -0.404 | -0.570 | -0.312 | -0.291 | -0.237 | -0.324 | -0.154 | NA | NA |
| TCGA-ZJ-AAXB-01A-11R | -5.203 | -0.622 | -0.867 | -1.161 | 0.008 | 0.221 | 0.194 | 0.000 | -0.486 | 0.266 | -0.004 | 3.571 | -0.904 | -0.436 | 0.073 | NA | NA |
| TCGA-C5-A7UC-01A-11R | 51.292 | 0.988 | 0.114 | 0.482 | -0.305 | 2.843 | -0.221 | 0.000 | -0.219 | -0.628 | -0.232 | 0.026 | 0.614 | -0.245 | -0.213 | NA | NA |
| TCGA-C5-A8XI-01A-11R | 48.320 | 0.216 | -0.232 | -0.553 | -0.154 | -0.360 | -0.226 | 0.000 | 0.080 | -0.428 | -0.067 | -0.246 | 0.856 | -0.077 | 0.022 | NA | NA |
| TCGA-C5-A7UE-01A-11R | 98.209 | -0.152 | 1.082 | -0.323 | -0.565 | -0.532 | -0.285 | 0.000 | -0.439 | -0.679 | -0.250 | -0.318 | -0.251 | -0.419 | -0.145 | NA | NA |
| TCGA-EA-A3HS-01A-11R | 111.318 | -0.949 | 2.131 | 1.397 | -0.366 | -0.361 | -0.272 | 0.000 | -0.503 | 0.002 | -0.240 | -0.290 | 2.257 | 0.043 | -0.262 | NA | NA |

| | | | | | | | | | | | | | | | | | |
|---|---|---|---|---|---|---|---|---|---|---|---|---|---|---|---|---|---|
| TCGA-EK-A2PI-01A-11R | 21.466 | -0.512 | -0.761 | -0.277 | -0.328 | -0.306 | -0.191 | 0.000 | -0.455 | 0.419 | -0.306 | -0.125 | 0.717 | -0.438 | 0.115 | NA | NA |
| TCGA-EK-A2RJ-01A-11R | 75.431 | -0.572 | 0.696 | -0.322 | -0.369 | -0.283 | -0.185 | 0.000 | -0.487 | -0.742 | 0.502 | -0.237 | -0.972 | -0.243 | -0.299 | NA | NA |
| TCGA-MA-AA43-01A-11R | -83.802 | -1.604 | -0.205 | -0.448 | 3.057 | 6.443 | 11.873 | 0.000 | 5.462 | 0.917 | 0.352 | -0.157 | 1.031 | 0.986 | 0.155 | NA | NA |
| TCGA-DS-A0VL-01A-21R | 86.354 | 0.652 | 0.497 | 0.203 | -0.450 | -0.483 | -0.300 | 0.000 | -0.505 | -0.433 | -0.265 | -0.307 | 0.131 | -0.280 | -0.234 | NA | NA |
| TCGA-ZJ-AAXN-01A-11R | 74.685 | 0.390 | 0.693 | -0.481 | -0.235 | 0.133 | -0.077 | 0.000 | 0.452 | -0.289 | -0.034 | 0.375 | -0.011 | 0.091 | 0.120 | NA | NA |
| TCGA-ZJ-AAXJ-01A-11R | 49.942 | -0.055 | -0.379 | 0.580 | -0.479 | -0.518 | -0.282 | 0.000 | -0.503 | -0.590 | -0.317 | -0.199 | 0.539 | -0.413 | 0.005 | NA | NA |
| TCGA-C5-A1BQ-01C-11R | 99.558 | 3.776 | -0.354 | 0.290 | -0.428 | -0.331 | -0.282 | 0.000 | -0.495 | -0.441 | -0.367 | -0.271 | -0.078 | -0.379 | -0.266 | NA | NA |
| TCGA-VS-A8EG-01A-11R | 102.645 | 0.226 | 0.960 | 1.781 | -0.547 | -0.483 | -0.277 | 0.000 | -0.463 | -0.657 | -0.358 | -0.332 | -0.826 | -0.410 | -0.308 | NA | NA |
| TCGA-C5-A1ML-01A-11R | 55.089 | -0.334 | 0.166 | -0.912 | -0.282 | -0.374 | -0.007 | 0.000 | 0.466 | -0.541 | -0.193 | -0.320 | 0.150 | -0.002 | 1.091 | NA | NA |
| TCGA-C5-A8YQ-01A-11R | 71.174 | 0.854 | -0.017 | -1.202 | -0.535 | -0.412 | -0.303 | 0.000 | -0.505 | -0.597 | -0.158 | -0.324 | 1.222 | -0.456 | -0.312 | NA | NA |
| TCGA-C5-A7XC-01A-11R | 64.604 | 0.261 | 0.368 | 1.115 | 0.117 | -0.333 | -0.174 | 0.000 | 0.132 | 0.593 | 0.109 | -0.247 | 0.376 | -0.251 | -0.204 | NA | NA |
| TCGA-EK-A2GZ-01A-11R | 61.812 | -0.343 | 0.423 | -0.889 | -0.442 | 0.109 | -0.291 | 0.000 | 0.475 | -0.344 | -0.279 | -0.281 | 1.267 | -0.313 | 0.516 | NA | NA |
| TCGA-EK-A3GJ-01A-21R | 53.518 | -0.619 | 0.195 | -0.971 | -0.353 | -0.501 | -0.303 | 0.000 | -0.500 | -0.736 | -0.010 | 0.900 | -0.422 | -0.190 | -0.272 | NA | NA |
| TCGA-EA-A5ZD-01A-11R | 62.473 | -0.228 | 0.341 | 1.049 | -0.214 | -0.313 | -0.264 | 0.000 | -0.442 | -0.177 | -0.273 | 0.542 | 0.341 | -0.362 | -0.290 | NA | NA |
| TCGA-EA-A97N-01A-11R | 154.864 | 0.959 | 3.178 | 3.258 | -0.002 | 0.663 | 0.011 | 0.000 | -0.471 | 0.525 | -0.212 | -0.131 | 0.839 | 0.892 | -0.138 | NA | NA |
| TCGA-VS-A9V4-01A-12R | 40.279 | 0.521 | -0.808 | -1.130 | -0.529 | -0.061 | -0.271 | 0.000 | -0.452 | -0.684 | 0.389 | -0.186 | -1.120 | -0.347 | -0.317 | NA | NA |
| TCGA-FU-A3HY-01A-11R | 59.909 | 0.449 | -0.028 | -0.411 | -0.622 | -0.380 | -0.266 | 0.000 | -0.084 | -0.384 | -0.311 | -0.289 | 1.474 | 0.436 | -0.210 | NA | NA |
| TCGA-JW-A5VK-01A-11R | 41.136 | -0.416 | -0.433 | -1.672 | -0.539 | -0.510 | -0.285 | 0.000 | -0.517 | -0.696 | -0.292 | -0.231 | 0.263 | -0.275 | -0.277 | NA | NA |
| TCGA-VS-A9UO-01A-11R | 19.676 | -0.901 | -1.019 | -0.184 | -0.677 | -0.529 | -0.303 | 0.000 | -0.516 | -0.772 | -0.357 | 0.604 | -1.193 | -0.478 | -0.317 | NA | NA |
| TCGA-VS-A9V0-01A-11R | 28.572 | -0.244 | -0.879 | 0.712 | -0.202 | -0.123 | -0.276 | 0.000 | -0.472 | -0.589 | -0.276 | -0.328 | -1.082 | -0.449 | -0.291 | NA | NA |

| | | | | | | | | | | | | | | | | | |
|---|---|---|---|---|---|---|---|---|---|---|---|---|---|---|---|---|---|
| TCGA-VS-A8QH-01A-11R | 64.337 | 1.292 | -0.485 | -0.968 | -0.484 | -0.428 | -0.248 | 0.000 | -0.494 | -0.573 | -0.198 | -0.295 | -1.139 | -0.435 | -0.290 | NA | NA |
| TCGA-DS-A0VK-01A-21R | 26.152 | 0.190 | -0.923 | 0.094 | -0.264 | -0.315 | -0.249 | 0.000 | -0.502 | 0.429 | -0.198 | -0.248 | 0.255 | -0.356 | -0.124 | NA | NA |
| TCGA-EA-A4BA-01A-21R | -56.734 | -0.522 | -1.032 | -0.198 | 5.857 | -0.192 | 0.212 | 0.000 | -0.517 | 0.373 | -0.244 | -0.202 | -0.786 | -0.001 | -0.093 | NA | NA |
| TCGA-EK-A2R9-01A-11R | -13.196 | -0.952 | -1.046 | -1.167 | 0.237 | 0.319 | -0.149 | 0.000 | 1.838 | 1.559 | -0.279 | -0.293 | 0.491 | -0.224 | 0.781 | NA | NA |
| TCGA-EK-A2RO-01A-11R | 4.861 | -1.012 | -0.320 | -0.961 | 0.270 | 0.808 | 0.189 | 0.000 | -0.029 | 1.944 | 0.064 | -0.249 | 0.602 | -0.267 | 0.262 | NA | NA |
| TCGA-IR-A3L7-01A-21R | 47.952 | -0.056 | -0.377 | 0.280 | -0.336 | -0.482 | -0.287 | 0.000 | -0.503 | -0.117 | -0.366 | -0.333 | -0.605 | -0.449 | -0.161 | NA | NA |
| TCGA-C5-A7CM-01A-11R | 5.433 | -0.783 | -0.985 | 0.188 | -0.317 | 0.252 | -0.197 | 0.000 | 3.688 | 0.075 | -0.155 | 0.677 | -0.995 | -0.354 | 0.303 | NA | NA |
| TCGA-VS-A9U6-01A-11R | 47.012 | -1.032 | 0.340 | 0.146 | -0.305 | -0.209 | -0.201 | 0.000 | 0.624 | 0.872 | -0.147 | 0.074 | 0.176 | -0.342 | -0.116 | NA | NA |
| TCGA-DR-A0ZM-01A-12R | 47.319 | 0.650 | -0.570 | -0.415 | -0.205 | -0.466 | -0.301 | 0.000 | -0.511 | -0.559 | -0.364 | -0.310 | -0.297 | -0.004 | -0.266 | NA | NA |
| TCGA-DR-A0ZL-01A-11R | 76.709 | 0.740 | -0.007 | 1.046 | -0.574 | -0.512 | -0.294 | 0.000 | -0.498 | -0.719 | -0.155 | -0.306 | -0.452 | -0.403 | -0.200 | NA | NA |
| TCGA-EX-A69L-01A-11R | 40.248 | 0.010 | -0.376 | -0.637 | -0.042 | -0.337 | -0.284 | 0.000 | -0.304 | -0.277 | 0.169 | -0.279 | -0.360 | -0.012 | -0.227 | NA | NA |
| TCGA-FU-A23L-01A-11R | 104.425 | 1.412 | 0.749 | 0.735 | -0.391 | -0.514 | -0.288 | 0.000 | -0.455 | -0.261 | -0.206 | -0.314 | 0.278 | -0.369 | -0.170 | NA | NA |
| TCGA-WL-A834-01A-11R | 81.217 | -0.088 | 0.719 | 0.009 | -0.429 | -0.160 | -0.282 | 0.000 | -0.280 | -0.532 | 0.363 | -0.320 | 0.234 | -0.353 | -0.171 | NA | NA |
| TCGA-VS-A9UL-01A-11R | 36.266 | 0.540 | -0.575 | 0.320 | -0.120 | 0.248 | 1.115 | 0.000 | -0.513 | 0.832 | -0.244 | -0.321 | -0.997 | -0.445 | 0.143 | NA | NA |
| TCGA-C5-A1MJ-01A-11R | -15.817 | -0.591 | -1.025 | -0.296 | 0.403 | 0.992 | 0.016 | 0.000 | 2.583 | 1.727 | 0.003 | -0.036 | -0.768 | 0.206 | -0.116 | NA | NA |
| TCGA-FU-A40J-01A-11R | 12.105 | -0.879 | -1.082 | 1.378 | -0.227 | -0.454 | -0.205 | 0.000 | -0.496 | 0.308 | -0.044 | -0.213 | -1.111 | -0.446 | -0.238 | NA | NA |
| TCGA-EX-A69M-01A-11R | 47.720 | -1.237 | 0.071 | -0.496 | -0.397 | -0.453 | -0.302 | 0.000 | -0.144 | -0.695 | -0.319 | -0.333 | -0.580 | -0.240 | -0.311 | NA | NA |
| TCGA-C5-A1BK-01B-11R | 53.983 | 0.019 | 0.016 | 0.344 | -0.064 | -0.468 | -0.285 | 0.000 | -0.514 | -0.548 | -0.249 | -0.334 | 0.100 | 0.479 | -0.276 | NA | NA |
| TCGA-DS-A0VM-01A-11R | 4.536 | -0.472 | -1.048 | -0.151 | -0.254 | 0.225 | -0.121 | 0.000 | 0.190 | 0.559 | -0.162 | 0.232 | 1.354 | -0.381 | -0.055 | NA | NA |
| TCGA-LP-A4AW-01A-11R | 27.344 | 1.064 | -0.379 | -0.665 | 0.676 | 0.450 | -0.143 | 0.000 | -0.420 | 1.303 | 0.339 | 0.424 | 0.710 | 0.067 | 0.076 | NA | NA |

| | | | | | | | | | | | | | | | | | |
|---|---|---|---|---|---|---|---|---|---|---|---|---|---|---|---|---|---|
| TCGA-C5-A1BJ-01A-11R | 77.971 | 0.402 | 0.622 | 0.234 | -0.219 | -0.354 | -0.266 | 0.000 | -0.449 | 0.323 | 0.061 | -0.148 | 0.708 | -0.244 | -0.155 | NA | NA |
| TCGA-EA-A439-01A-11R | 5.887 | 0.759 | -0.718 | -0.517 | 0.187 | 2.633 | 0.052 | 0.000 | -0.047 | 2.027 | 0.065 | -0.327 | -0.384 | -0.371 | -0.093 | NA | NA |
| TCGA-EA-A44S-01A-12R | 55.299 | 0.401 | -0.137 | -0.044 | -0.170 | -0.487 | -0.237 | 0.000 | -0.508 | -0.485 | -0.303 | -0.316 | -0.322 | 0.422 | -0.244 | NA | NA |
| TCGA-VS-A9UZ-01A-11R | 33.132 | 0.437 | -1.040 | -0.109 | -0.397 | -0.481 | -0.273 | 0.000 | -0.505 | -0.385 | -0.244 | -0.331 | -1.138 | -0.161 | -0.281 | NA | NA |
| TCGA-C5-A7X5-01A-11R | 49.141 | -0.619 | 0.275 | -0.243 | 0.195 | -0.331 | -0.084 | 0.000 | 0.217 | -0.269 | -0.112 | -0.205 | 2.555 | -0.459 | 0.232 | NA | NA |
| TCGA-VS-A9UU-01A-11R | 119.325 | 0.471 | 1.644 | 0.024 | -0.552 | -0.520 | -0.303 | 0.000 | -0.438 | -0.578 | -0.229 | 0.754 | 0.097 | -0.382 | -0.058 | NA | NA |
| TCGA-C5-A8XH-01A-11R | 43.262 | -0.759 | 0.124 | -0.008 | -0.395 | -0.400 | -0.287 | 0.000 | -0.237 | -0.486 | -0.224 | 0.094 | 0.219 | 1.110 | -0.080 | NA | NA |
| TCGA-EK-A2PM-01A-11R | 61.053 | -1.267 | 0.708 | -1.438 | -0.541 | -0.316 | 1.221 | 0.000 | -0.502 | -0.513 | -0.333 | -0.322 | 3.055 | -0.405 | 0.373 | NA | NA |
| TCGA-C5-A1BM-01A-11R | 143.333 | 1.942 | 1.790 | 0.996 | -0.204 | -0.367 | -0.233 | 0.000 | -0.515 | -0.504 | -0.198 | -0.182 | -0.600 | -0.182 | -0.235 | NA | NA |
| TCGA-EK-A2RE-01A-11R | 158.661 | 0.178 | 3.239 | 0.599 | -0.310 | -0.256 | -0.249 | 0.000 | -0.020 | -0.509 | -0.269 | 0.012 | 1.818 | 0.365 | 0.508 | NA | NA |
| TCGA-IR-A3LH-01A-21R | -40.254 | 1.110 | -0.666 | -1.055 | 0.520 | 4.587 | 1.990 | 0.000 | -0.155 | 2.294 | 0.225 | 0.030 | 0.461 | 4.043 | 0.062 | NA | NA |
| TCGA-EX-A1H5-01A-31R | 94.636 | 0.942 | 0.703 | -0.851 | -0.442 | -0.461 | -0.301 | 0.000 | -0.476 | -0.461 | -0.279 | -0.312 | 0.821 | -0.410 | 0.330 | NA | NA |
| TCGA-VS-A94Y-01A-11R | -42.935 | 1.782 | -0.693 | 0.512 | -0.368 | 0.605 | -0.245 | 0.000 | 0.071 | 1.882 | 2.197 | 2.026 | 0.037 | 11.238 | -0.131 | NA | NA |
| TCGA-EA-A1QT-01A-11R | 69.869 | -0.432 | 0.447 | -0.677 | -0.536 | -0.441 | -0.299 | 0.000 | -0.515 | -0.470 | -0.290 | -0.264 | 0.137 | -0.311 | -0.139 | NA | NA |
| TCGA-VS-A953-01A-11R | 68.256 | -0.100 | 0.740 | -0.786 | 0.526 | -0.470 | -0.202 | 0.000 | -0.481 | -0.803 | -0.284 | -0.327 | 4.580 | -0.460 | 0.546 | NA | NA |
| TCGA-EK-A2R8-01A-21R | 12.340 | -0.077 | -0.667 | -0.359 | 0.472 | 1.338 | -0.008 | 0.000 | 1.018 | -0.405 | 1.036 | 0.124 | -0.953 | -0.309 | -0.106 | NA | NA |
| TCGA-C5-A2LS-01A-22R | 23.620 | -0.834 | -0.989 | -0.069 | -0.545 | -0.529 | -0.298 | 0.000 | -0.455 | -0.731 | -0.261 | -0.307 | -1.180 | -0.463 | -0.315 | NA | NA |
| TCGA-EK-A2RA-01A-11R | 52.622 | -0.258 | -0.132 | 0.169 | -0.201 | -0.528 | -0.301 | 0.000 | -0.473 | -0.741 | -0.337 | -0.334 | 0.603 | -0.315 | -0.308 | NA | NA |
| TCGA-MY-A5BF-01A-11R | 107.476 | -1.123 | 1.999 | -0.220 | -0.313 | -0.473 | -0.303 | 0.000 | -0.517 | -0.715 | 0.244 | -0.323 | 0.316 | 0.198 | -0.243 | NA | NA |
| TCGA-C5-A1MN-01A-11R | 42.580 | 0.285 | 0.203 | 0.955 | -0.618 | -0.177 | 0.105 | 0.000 | 0.824 | 1.522 | -0.223 | 0.852 | 3.703 | 1.298 | -0.214 | NA | NA |

| | | | | | | | | | | | | | | | | | |
|---|---|---|---|---|---|---|---|---|---|---|---|---|---|---|---|---|---|
| TCGA-ZJ-A8QR-01A-11R | 51.923 | 0.158 | -0.344 | -0.072 | -0.535 | -0.472 | -0.288 | 0.000 | -0.446 | -0.677 | -0.276 | -0.267 | 0.281 | -0.156 | 0.658 | NA | NA |
| TCGA-EX-A1H6-01B-11R | 0.685 | -1.076 | -1.246 | 0.253 | -0.133 | -0.494 | -0.238 | 0.000 | -0.360 | 0.442 | -0.189 | -0.218 | -0.877 | -0.460 | 0.051 | NA | NA |
| TCGA-VS-A9V3-01A-11R | 64.088 | 0.442 | -0.138 | 0.626 | -0.429 | -0.428 | -0.292 | 0.000 | 1.139 | -0.560 | -0.139 | -0.324 | -0.682 | -0.288 | -0.312 | NA | NA |
| TCGA-VS-A94X-01A-11R | 93.392 | 1.183 | 0.422 | 0.065 | -0.607 | -0.444 | -0.289 | 0.000 | 0.054 | -0.655 | -0.286 | -0.085 | -0.366 | -0.354 | -0.306 | NA | NA |
| TCGA-ZJ-AAXA-01A-11R | 131.361 | 0.394 | 1.967 | 1.840 | -0.313 | -0.224 | -0.248 | 0.000 | -0.349 | -0.496 | -0.276 | -0.317 | -0.012 | -0.423 | -0.281 | NA | NA |
| TCGA-C5-A905-01A-11R | 173.734 | 0.770 | 2.903 | 1.918 | -0.545 | -0.504 | -0.286 | 0.000 | -0.511 | -0.702 | -0.096 | -0.319 | -0.718 | -0.451 | -0.314 | NA | NA |
| TCGA-Q1-A6DW-01A-11R | 69.694 | -0.663 | 0.517 | 1.150 | -0.468 | -0.334 | -0.199 | 0.000 | 0.007 | -0.440 | -0.284 | -0.319 | 0.205 | -0.246 | -0.270 | NA | NA |
| TCGA-C5-A7UI-01A-11R | 29.186 | -1.608 | 0.008 | -1.600 | -0.260 | -0.509 | -0.227 | 0.000 | -0.510 | -0.416 | -0.183 | 1.258 | -0.130 | -0.224 | -0.156 | NA | NA |
| TCGA-C5-A2LT-01A-11R | -5.027 | -0.984 | -0.683 | -0.273 | 0.091 | 0.436 | -0.195 | 0.000 | 1.403 | 0.872 | 0.566 | 1.805 | 1.068 | -0.296 | -0.012 | NA | NA |
| TCGA-ZJ-AB0I-01A-11R | 57.593 | 0.008 | 1.507 | -0.148 | -0.455 | -0.404 | -0.269 | 0.000 | 0.193 | 0.198 | 0.239 | 1.354 | 0.916 | 5.977 | 0.099 | NA | NA |
| TCGA-DS-A7WF-01A-11R | -0.932 | 0.588 | -0.974 | 1.617 | 0.830 | 0.049 | -0.272 | 0.000 | 0.668 | 4.170 | -0.192 | 0.112 | -1.036 | -0.363 | -0.297 | NA | NA |
| TCGA-EA-A411-01A-11R | 21.499 | -0.029 | 0.039 | 0.408 | 0.371 | 1.537 | -0.098 | 0.000 | -0.326 | 2.419 | 0.389 | -0.210 | -0.392 | 0.179 | -0.076 | NA | NA |
| TCGA-EA-A3QD-01A-32R | 16.971 | -0.233 | -0.417 | -0.276 | 0.738 | 0.001 | -0.278 | 0.000 | -0.455 | 1.185 | -0.066 | -0.116 | -0.813 | 0.160 | -0.131 | NA | NA |
| TCGA-ZX-AA5X-01A-11R | 12.730 | -0.511 | -0.274 | 0.817 | 0.606 | 0.028 | -0.207 | 0.000 | -0.396 | 2.881 | 0.192 | -0.137 | 0.390 | -0.163 | -0.039 | NA | NA |
| TCGA-IR-A3LB-01A-11R | -39.257 | -0.982 | -1.107 | -0.581 | 0.714 | 0.702 | -0.129 | 0.000 | 1.646 | 4.688 | 0.311 | 0.240 | -0.574 | -0.188 | 0.620 | NA | NA |
| TCGA-DS-A1OA-01A-11R | 49.397 | -0.339 | 0.121 | 1.602 | -0.051 | 0.114 | -0.246 | 0.000 | 0.503 | 0.247 | -0.234 | -0.251 | -0.175 | -0.083 | -0.191 | NA | NA |
| TCGA-C5-A8XK-01A-11R | 56.618 | 0.098 | 0.002 | -1.118 | -0.403 | -0.253 | -0.291 | 0.000 | 0.957 | -0.639 | 0.588 | -0.250 | 0.676 | -0.106 | -0.320 | NA | NA |
| TCGA-DS-A5RQ-01A-11R | 81.501 | 0.239 | 0.543 | 0.356 | -0.372 | -0.425 | -0.287 | 0.000 | 0.369 | -0.402 | -0.182 | -0.305 | -0.207 | -0.326 | -0.177 | NA | NA |
| TCGA-VS-A8QM-01A-11R | 63.022 | 0.148 | 0.526 | 0.724 | 0.527 | -0.356 | -0.194 | 0.000 | -0.081 | 0.474 | -0.269 | -0.322 | -0.203 | 0.107 | -0.176 | NA | NA |
| TCGA-C5-A1BE-01B-11R | 117.104 | 1.436 | 1.303 | -0.260 | -0.397 | -0.351 | -0.283 | 0.000 | -0.501 | -0.408 | -0.220 | -0.244 | 0.222 | 0.208 | -0.204 | NA | NA |

| | | | | | | | | | | | | | | | | | |
|---|---|---|---|---|---|---|---|---|---|---|---|---|---|---|---|---|---|
| TCGA-DS-A3LQ-01A-21R | 60.830 | 1.354 | -0.151 | 1.320 | -0.050 | 0.081 | -0.225 | 0.000 | -0.481 | -0.054 | -0.130 | -0.019 | 0.469 | 0.171 | -0.031 | NA | NA |
| TCGA-C5-A7CK-01A-11R | 72.304 | 0.501 | 0.755 | 0.092 | -0.178 | 0.985 | -0.043 | 0.000 | -0.435 | 0.350 | -0.130 | -0.068 | 1.804 | -0.428 | -0.046 | NA | NA |
| TCGA-C5-A3HL-01A-11R | 145.559 | 1.451 | 1.992 | 0.264 | -0.510 | -0.442 | -0.251 | 0.000 | -0.045 | -0.488 | -0.351 | -0.260 | 0.735 | -0.323 | -0.166 | NA | NA |
| TCGA-C5-A0TN-01A-21R | -40.089 | -0.720 | -0.024 | 0.152 | 0.458 | 6.569 | 2.970 | 0.000 | -0.017 | -0.691 | -0.256 | 4.161 | 2.448 | 0.613 | 0.942 | NA | NA |
| TCGA-EK-A2RB-01A-11R | 51.026 | -0.760 | -0.087 | -0.631 | -0.567 | -0.495 | -0.293 | 0.000 | -0.495 | -0.681 | -0.356 | -0.331 | 0.274 | -0.459 | -0.147 | NA | NA |
| TCGA-VS-A8EH-01A-11R | 81.652 | -0.148 | 0.645 | -0.287 | -0.512 | -0.489 | -0.278 | 0.000 | -0.189 | -0.336 | -0.372 | -0.328 | -0.170 | -0.429 | -0.317 | NA | NA |
| TCGA-EA-A5FO-01A-21R | 66.474 | 0.335 | -0.086 | 0.133 | -0.400 | -0.528 | -0.304 | 0.000 | -0.514 | -0.814 | -0.372 | -0.334 | -0.127 | -0.465 | -0.316 | NA | NA |
| TCGA-LP-A5U3-01A-11R | 90.245 | 1.432 | 0.217 | 0.733 | -0.476 | -0.497 | -0.290 | 0.000 | -0.513 | -0.754 | -0.348 | -0.328 | 0.046 | -0.223 | -0.328 | NA | NA |
| TCGA-VS-A9UR-01A-11R | 79.785 | 1.484 | -0.066 | 0.611 | -0.194 | -0.432 | -0.158 | 0.000 | -0.505 | -0.654 | -0.213 | -0.329 | -1.167 | -0.386 | -0.162 | NA | NA |
| TCGA-EA-A3HR-01A-11R | 179.009 | 1.398 | 2.965 | 1.264 | -0.377 | -0.514 | -0.232 | 0.000 | -0.512 | -0.500 | -0.307 | -0.333 | 0.206 | -0.410 | -0.289 | NA | NA |
| TCGA-DG-A2KK-01A-11R | 17.025 | 0.682 | -1.146 | -1.641 | 0.319 | 0.295 | -0.297 | 0.000 | -0.341 | -0.571 | 0.999 | -0.164 | -1.179 | -0.252 | -0.301 | NA | NA |
| TCGA-ZJ-A8QQ-01A-11R | 108.033 | -0.238 | 1.586 | -0.407 | -0.528 | -0.461 | -0.269 | 0.000 | 0.444 | -0.581 | -0.213 | -0.116 | 0.897 | -0.216 | -0.212 | NA | NA |
| TCGA-EX-A449-01A-11R | 29.831 | 0.061 | -1.138 | -0.604 | -0.592 | -0.507 | -0.278 | 0.000 | -0.505 | -0.716 | -0.366 | -0.334 | -1.162 | -0.397 | -0.319 | NA | NA |
| TCGA-VS-A9UB-01A-22R | 42.186 | 0.204 | -0.591 | 2.672 | -0.052 | -0.267 | -0.219 | 0.000 | -0.435 | -0.807 | -0.285 | -0.330 | 0.199 | -0.026 | 0.225 | NA | NA |
| TCGA-C5-A7CJ-01A-11R | 81.763 | 0.646 | 0.983 | 1.078 | 0.247 | -0.100 | -0.192 | 0.000 | -0.466 | 1.332 | 0.087 | -0.147 | 0.182 | -0.065 | -0.111 | NA | NA |
| TCGA-R2-A69V-01A-11R | 14.535 | -0.077 | -0.483 | -0.563 | 1.033 | -0.073 | -0.193 | 0.000 | -0.494 | -0.367 | 0.147 | -0.138 | 0.029 | 1.026 | -0.190 | NA | NA |
| TCGA-VS-A94W-01A-12R | 63.557 | -0.190 | 0.353 | -0.703 | -0.066 | -0.485 | -0.270 | 0.000 | -0.471 | -0.620 | 0.658 | -0.327 | -0.493 | 0.110 | -0.169 | NA | NA |
| TCGA-C5-A1MP-01A-11R | 88.085 | 0.470 | 1.326 | -0.521 | 0.649 | -0.251 | -0.099 | 0.000 | -0.482 | 0.666 | 0.090 | -0.151 | -0.568 | 0.108 | -0.118 | NA | NA |
| TCGA-EX-A8YF-01A-11R | 30.618 | -1.321 | -0.533 | 1.073 | -0.446 | -0.451 | -0.285 | 0.000 | -0.454 | -0.589 | -0.329 | -0.333 | -1.136 | -0.402 | 0.120 | NA | NA |
| TCGA-C5-A7CH-01A-11R | 136.631 | 0.426 | 2.910 | -0.211 | -0.006 | -0.203 | -0.284 | 0.000 | 0.133 | -0.386 | 0.602 | 0.469 | 0.757 | 1.666 | 0.093 | NA | NA |

| | | | | | | | | | | | | | | | | | |
|---|---|---|---|---|---|---|---|---|---|---|---|---|---|---|---|---|---|
| TCGA-HM-A3JK-01A-11R | -14.535 | -0.861 | -0.406 | -0.534 | 0.893 | 1.002 | 0.079 | 0.000 | -0.444 | 3.108 | -0.267 | -0.283 | 0.580 | 0.644 | 0.259 | NA | NA |
| TCGA-C5-A1MF-01A-11R | 5.269 | -1.425 | -1.013 | -1.729 | -0.298 | -0.468 | 0.024 | 0.000 | -0.504 | -0.315 | -0.277 | -0.256 | -0.815 | -0.303 | -0.076 | NA | NA |
| TCGA-C5-A1BL-01A-11R | 73.186 | -0.226 | 0.489 | -0.019 | -0.465 | -0.480 | -0.293 | 0.000 | -0.513 | -0.719 | -0.341 | -0.316 | 0.364 | 0.088 | -0.305 | NA | NA |
| TCGA-EA-A78R-01A-11R | 19.317 | -0.278 | -0.639 | -1.291 | 0.217 | -0.155 | -0.270 | 0.000 | -0.391 | 0.071 | -0.170 | -0.069 | 0.388 | -0.007 | -0.189 | NA | NA |
| TCGA-EK-A3GM-01A-11R | 55.064 | 0.968 | -0.965 | 4.211 | -0.649 | -0.473 | -0.281 | 0.000 | -0.137 | -0.784 | -0.267 | -0.297 | -1.172 | -0.430 | -0.301 | NA | NA |
| TCGA-C5-A1BI-01B-11R | 81.556 | -0.573 | 1.093 | 0.704 | -0.167 | -0.355 | -0.247 | 0.000 | -0.462 | 0.112 | -0.104 | -0.222 | -0.112 | 0.015 | -0.252 | NA | NA |
| TCGA-VS-A9UQ-01A-21R | 35.783 | 0.713 | -0.911 | 1.381 | -0.123 | -0.346 | -0.065 | 0.000 | -0.415 | 0.525 | -0.251 | -0.265 | -1.025 | -0.422 | -0.285 | NA | NA |
| TCGA-HM-A6W2-06A-22R | -0.472 | 0.850 | -0.816 | 1.030 | 3.175 | -0.274 | 4.651 | 0.000 | 2.492 | -0.751 | -0.202 | -0.334 | -1.061 | 0.555 | -0.273 | NA | NA |
| TCGA-UC-A7PG-06A-11R | 97.850 | 0.034 | 1.146 | 0.617 | -0.306 | -0.082 | -0.238 | 0.000 | -0.470 | -0.734 | -0.212 | -0.334 | -0.294 | -0.322 | -0.289 | NA | NA |
| TCGA-EX-A3L1-01A-11R | 26.790 | 1.992 | -0.008 | 0.819 | 0.436 | 3.061 | 0.249 | 0.000 | 0.035 | -0.220 | 0.210 | 0.038 | 0.256 | 3.176 | 0.072 | NA | NA |
| TCGA-C5-A1MQ-01A-11R | 0.362 | -0.723 | -0.768 | -1.480 | 0.193 | -0.027 | -0.023 | 0.000 | -0.263 | 0.167 | -0.231 | 0.094 | -0.704 | 1.251 | -0.193 | NA | NA |
| TCGA-VS-A9UP-01A-11R | 46.465 | 1.498 | -1.045 | 1.045 | -0.517 | -0.375 | 0.144 | 0.000 | -0.258 | 0.147 | -0.211 | -0.208 | -1.096 | -0.423 | 0.574 | NA | NA |
| TCGA-VS-A9UI-01A-11R | 138.281 | 2.521 | 1.195 | 1.093 | -0.512 | -0.507 | -0.304 | 0.000 | -0.516 | -0.779 | -0.091 | -0.330 | -0.725 | -0.182 | -0.319 | NA | NA |
| TCGA-UC-A7PF-01A-11R | 35.615 | -0.192 | -0.359 | -0.528 | -0.185 | -0.322 | -0.242 | 0.000 | -0.412 | 0.305 | -0.330 | -0.214 | 0.548 | 0.005 | -0.150 | NA | NA |
| TCGA-DS-A1O9-01A-11R | 4.623 | -1.178 | -0.485 | -0.925 | 0.040 | 1.023 | -0.182 | 0.000 | -0.470 | 0.677 | -0.046 | -0.333 | 1.275 | -0.393 | 0.792 | NA | NA |
| TCGA-VS-A9UD-01A-11R | 8.677 | -0.689 | -0.998 | -1.551 | 0.079 | -0.440 | -0.303 | 0.000 | -0.455 | -0.779 | -0.287 | -0.279 | -0.957 | 0.530 | 0.013 | NA | NA |
| TCGA-2W-A8YY-01A-11R | 11.920 | 0.259 | -1.094 | -0.617 | -0.114 | -0.006 | 1.128 | 0.000 | -0.460 | 0.957 | -0.114 | -0.332 | -1.095 | 0.087 | 0.260 | NA | NA |
| TCGA-MY-A5BE-01A-21R | -3.706 | -1.163 | -0.872 | -1.016 | 0.503 | -0.481 | -0.259 | 0.000 | -0.506 | -0.178 | -0.252 | -0.253 | 0.323 | 0.815 | 0.018 | NA | NA |
| TCGA-ZJ-AAXU-01A-11R | 44.067 | -0.270 | -0.236 | -1.133 | -0.369 | -0.510 | -0.300 | 0.000 | -0.513 | -0.356 | -0.251 | 0.464 | 0.209 | -0.318 | -0.162 | NA | NA |
| TCGA-C5-A1BF-01B-11R | -1.446 | -1.423 | -0.181 | -0.088 | 1.362 | 0.308 | 0.846 | 0.000 | 5.895 | 0.831 | 0.760 | -0.310 | 1.239 | -0.248 | 0.306 | NA | NA |

| | | | | | | | | | | | | | | | | | |
|---|---|---|---|---|---|---|---|---|---|---|---|---|---|---|---|---|---|
| TCGA-MY-A5BD-01A-11R | 58.902 | -0.052 | 0.025 | -1.199 | 0.010 | -0.528 | -0.286 | 0.000 | 0.129 | -0.703 | -0.236 | -0.326 | -1.174 | -0.447 | -0.303 | NA | NA |
| TCGA-4J-AA1J-01A-21R | 54.379 | 1.291 | -0.488 | 2.207 | -0.314 | -0.278 | -0.295 | 0.000 | -0.469 | 0.830 | -0.310 | -0.193 | 0.810 | -0.179 | -0.190 | NA | NA |
| TCGA-UC-A7PG-01A-11R | 193.523 | 0.114 | 3.999 | 1.020 | -0.450 | -0.364 | -0.242 | 0.000 | -0.467 | -0.386 | -0.132 | -0.217 | 0.252 | -0.354 | -0.285 | NA | NA |
| TCGA-EK-A2RL-01A-11R | 28.484 | -0.195 | -1.039 | 0.419 | -0.488 | -0.441 | -0.243 | 0.000 | 0.754 | -0.509 | -0.368 | -0.316 | -1.140 | -0.476 | -0.303 | NA | NA |
| TCGA-C5-A3HE-01A-21R | 15.895 | -0.921 | -0.986 | -0.790 | -0.295 | -0.435 | -0.192 | 0.000 | -0.258 | -0.497 | -0.017 | -0.238 | -1.138 | -0.408 | -0.272 | NA | NA |
| TCGA-EA-A43B-01A-81R | 15.905 | -1.09 | -0.443 | -1.501 | -0.025 | -0.137 | -0.292 | 0.000 | 0.007 | -0.483 | 0.800 | 0.019 | -0.114 | 0.537 | -0.206 | NA | NA |
| TCGA-ZJ-A8QO-01A-11R | -0.502 | -1.26 | -0.439 | -1.700 | 0.185 | 0.684 | -0.265 | 0.000 | 1.614 | 0.081 | -0.188 | 0.159 | 1.035 | 0.645 | 0.078 | NA | NA |
| TCGA-EA-A410-01A-11R | -22.615 | -0.030 | -0.964 | -0.271 | 0.952 | 1.530 | 0.332 | 0.000 | 0.504 | -0.440 | 0.230 | -0.332 | -0.727 | 0.628 | 15.251 | NA | NA |
| TCGA-FU-A3EO-11A-13R | -49.157 | -1.894 | -1.299 | 2.294 | 2.833 | -0.332 | -0.054 | 0.000 | -0.455 | 0.748 | -0.037 | -0.335 | 0.605 | -0.467 | 3.206 | NA | NA |
| TCGA-EK-A2PK-01A-11R | 17.047 | -1.468 | -0.589 | -0.153 | -0.044 | -0.417 | 0.994 | 0.000 | -0.513 | -0.384 | -0.079 | -0.084 | -1.046 | -0.317 | -0.201 | NA | NA |
| TCGA-C5-A1BN-01B-11R | 62.779 | -0.25 | 0.784 | 1.043 | -0.400 | 0.723 | -0.240 | 0.000 | -0.392 | -0.316 | -0.321 | 0.244 | 2.360 | 0.787 | -0.085 | NA | NA |
| TCGA-MA-AA41-01A-11R | 33.945 | -0.99 | -0.359 | -1.011 | -0.525 | -0.469 | -0.287 | 0.000 | -0.413 | -0.665 | -0.349 | -0.304 | 0.912 | -0.003 | -0.169 | NA | NA |
| TCGA-VS-A9UM-01A-11R | 97.429 | 0.253 | 1.192 | 0.900 | -0.265 | -0.240 | -0.203 | 0.000 | -0.494 | 0.265 | -0.239 | -0.222 | -0.235 | -0.246 | -0.249 | NA | NA |
| TCGA-Q1-A5R2-01A-11R | 33.675 | -0.83 | -0.209 | -1.548 | -0.114 | -0.336 | -0.288 | 0.000 | -0.458 | -0.090 | -0.313 | -0.265 | 0.428 | -0.192 | -0.256 | NA | NA |
| TCGA-DS-A7WI-01A-12R | 24.827 | -0.67 | -0.883 | -0.879 | -0.494 | -0.519 | -0.304 | 0.000 | 0.807 | -0.727 | -0.165 | -0.019 | -0.700 | -0.443 | -0.243 | NA | NA |
| TCGA-VS-A8QA-01A-11R | 50.536 | 1.403 | -1.095 | -0.497 | -0.644 | -0.537 | -0.304 | 0.000 | -0.502 | -0.820 | -0.362 | -0.335 | -1.199 | -0.397 | -0.324 | NA | NA |
| TCGA-GH-A9DA-01A-21R | 31.696 | 1.062 | -0.282 | -0.145 | 0.566 | 1.562 | -0.264 | 0.000 | 0.330 | -0.294 | -0.036 | 1.025 | 0.185 | -0.089 | -0.229 | NA | NA |
| TCGA-MA-AA42-01A-12R | 19.595 | 0.026 | -0.98 | -1.761 | -0.061 | -0.481 | -0.302 | 0.000 | 0.059 | -0.632 | -0.370 | -0.317 | -0.270 | 0.173 | 1.377 | NA | NA |
| TCGA-ZJ-AB0H-01A-11R | 77.799 | 0.029 | 0.808 | 0.362 | -0.085 | 0.035 | -0.233 | 0.000 | -0.374 | -0.228 | -0.154 | -0.294 | 0.135 | -0.014 | -0.260 | NA | NA |
| TCGA-FU-A3WB-01A-11R | 87.990 | 0.914 | 0.444 | 0.570 | -0.430 | -0.378 | -0.278 | 0.000 | -0.336 | -0.257 | -0.318 | -0.255 | -0.338 | -0.460 | 0.116 | NA | NA |

| | | | | | | | | | | | | | | | | | |
|---|---|---|---|---|---|---|---|---|---|---|---|---|---|---|---|---|---|
| TCGA-IR-A3LK-01A-12R | 79.587 | 1.219 | -0.02 | 0.645 | -0.449 | -0.483 | -0.303 | 0.000 | -0.478 | -0.763 | 0.488 | -0.296 | -0.729 | -0.258 | -0.293 | NA | NA |
| TCGA-VS-A8Q8-01A-11R | 42.559 | -1.21 | -0.02 | -0.296 | -0.394 | -0.505 | -0.289 | 0.000 | -0.336 | -0.555 | -0.285 | -0.186 | 0.807 | -0.231 | -0.148 | NA | NA |
| TCGA-EA-A3Y4-01A-51R | 17.725 | -0.14 | -0.70 | -1.411 | -0.249 | -0.199 | 0.041 | 0.000 | -0.267 | -0.264 | 2.774 | 1.584 | 0.600 | -0.214 | -0.176 | NA | NA |
| TCGA-VS-A9UT-01A-11R | -56.912 | -1.61 | -1.09 | -0.796 | 2.515 | 1.268 | 6.485 | 0.000 | -0.298 | 0.349 | -0.111 | -0.334 | -1.061 | 0.952 | -0.047 | NA | NA |
| TCGA-EK-A2RC-01A-11R | 31.978 | -0.07 | -0.68 | 0.926 | -0.064 | -0.424 | -0.178 | 0.000 | -0.503 | 0.368 | -0.144 | -0.320 | -0.389 | -0.374 | -0.209 | NA | NA |
| TCGA-C5-A1MH-01A-11R | 79.457 | 0.939 | 0.761 | -0.208 | 0.123 | -0.166 | -0.281 | 0.000 | -0.504 | 0.124 | -0.219 | 0.089 | -0.207 | 0.660 | -0.134 | NA | NA |
| TCGA-MA-AA3Y-01A-11R | 56.646 | -0.29 | 0.953 | 0.126 | 0.067 | 0.459 | -0.098 | 0.000 | -0.255 | -0.479 | 0.383 | 3.562 | 1.387 | -0.137 | -0.041 | NA | NA |
| TCGA-DG-A2KH-01A-21R | 24.181 | -0.51 | -1.157 | 0.708 | -0.616 | -0.530 | -0.271 | 0.000 | 1.240 | -0.799 | -0.361 | -0.322 | -1.192 | -0.469 | -0.192 | NA | NA |
| TCGA-JX-A3Q0-01A-11R | 41.955 | -0.47 | -0.371 | -0.893 | -0.422 | -0.444 | -0.292 | 0.000 | -0.509 | -0.626 | -0.142 | -0.299 | -0.533 | -0.146 | -0.318 | NA | NA |
| TCGA-EK-A2IP-01A-11R | 30.202 | 0.239 | -0.532 | 0.674 | -0.513 | -0.487 | -0.275 | 0.000 | -0.166 | 0.123 | -0.295 | 0.184 | 1.643 | 1.501 | -0.076 | NA | NA |
| TCGA-VS-A9U7-01A-11R | 53.797 | -0.68 | 0.263 | -0.466 | -0.183 | -0.317 | -0.285 | 0.000 | -0.494 | -0.493 | -0.275 | -0.176 | -0.421 | 0.151 | -0.295 | NA | NA |
| TCGA-JW-A5VL-01A-11R | 70.861 | 0.545 | 0.120 | 0.268 | -0.352 | -0.441 | -0.289 | 0.000 | -0.448 | -0.735 | 0.522 | -0.313 | -0.010 | -0.025 | -0.326 | NA | NA |
| TCGA-HM-A4S6-01A-11R | 77.391 | 0.854 | 0.608 | 0.449 | -0.156 | 0.188 | -0.234 | 0.000 | -0.345 | -0.286 | 1.031 | -0.311 | -0.577 | 0.766 | -0.278 | NA | NA |
| TCGA-EA-A5ZE-01A-11R | 20.088 | -0.580 | -0.58 | -0.799 | 0.141 | -0.226 | -0.244 | 0.000 | 2.605 | 0.143 | -0.171 | -0.324 | 0.920 | -0.305 | -0.208 | NA | NA |
| TCGA-ZJ-AAXF-01A-31R | 89.870 | 0.387 | 1.179 | 0.002 | 0.052 | -0.312 | -0.208 | 0.000 | -0.320 | 0.556 | -0.221 | -0.265 | -0.281 | 0.234 | 0.034 | NA | NA |
| TCGA-IR-A3LA-01A-11R | 52.224 | -0.117 | -0.30 | 0.817 | -0.420 | -0.427 | -0.135 | 0.000 | -0.498 | -0.382 | -0.039 | -0.331 | -1.100 | -0.371 | -0.290 | NA | NA |
| TCGA-C5-A2LX-01A-11R | 0.762 | -1.293 | -0.63 | -1.495 | 0.687 | -0.453 | -0.253 | 0.000 | -0.481 | -0.365 | 0.237 | -0.066 | -0.218 | 0.693 | -0.017 | NA | NA |
| TCGA-EK-A2R7-01A-11R | 13.317 | -0.355 | -1.07 | -0.978 | -0.181 | -0.486 | 0.967 | 0.000 | -0.163 | -0.073 | -0.150 | 0.153 | -0.882 | -0.266 | 0.343 | NA | NA |
| TCGA-ZJ-AAX8-01A-11R | 41.469 | -0.071 | 0.33 | 0.561 | -0.023 | 0.529 | 0.023 | 0.000 | -0.445 | 2.428 | -0.177 | 0.417 | 0.435 | -0.183 | -0.105 | NA | NA |
| TCGA-VS-A9UJ-01A-11R | 21.196 | 1.470 | -0.24 | 0.480 | 3.447 | -0.144 | -0.208 | 0.000 | 0.507 | 0.610 | -0.287 | 0.358 | -1.050 | -0.258 | -0.279 | NA | NA |

| | | | | | | | | | | | | | | | | | |
|---|---|---|---|---|---|---|---|---|---|---|---|---|---|---|---|---|---|
| TCGA-MA-AA3Z-01A-11R | 104.184 | 1.579 | 0.604 | -0.800 | -0.477 | -0.486 | -0.300 | 0.000 | -0.502 | -0.769 | -0.328 | -0.334 | -0.364 | -0.383 | -0.328 | NA | NA |
| TCGA-LP-A4AV-01A-11R | -5.822 | -1.496 | -1.03 | -1.852 | -0.130 | -0.317 | -0.245 | 0.000 | -0.465 | 0.154 | 0.230 | -0.261 | -0.060 | 0.189 | -0.226 | NA | NA |
| TCGA-VS-A8EJ-01A-11R | 14.697 | -0.863 | -0.99 | -1.044 | -0.495 | -0.496 | 1.830 | 0.000 | -0.506 | -0.253 | -0.215 | -0.202 | -1.170 | -0.303 | 0.605 | NA | NA |
| TCGA-HM-A3JJ-01A-21R | 36.448 | 0.604 | -0.08 | -0.975 | -0.166 | 0.182 | -0.177 | 0.000 | 1.778 | 3.101 | -0.127 | 0.032 | 1.271 | -0.342 | -0.088 | NA | NA |
| TCGA-VS-A8EK-01A-12R | 62.112 | -0.662 | 0.237 | -0.451 | -0.553 | -0.461 | -0.303 | 0.000 | -0.422 | -0.666 | -0.269 | -0.307 | -0.053 | -0.316 | -0.254 | NA | NA |
| TCGA-C5-A901-01A-11R | 167.406 | 0.557 | 3.078 | 2.026 | -0.246 | -0.401 | -0.177 | 0.000 | -0.493 | -0.345 | 0.017 | -0.241 | -0.055 | -0.122 | -0.222 | NA | NA |
| TCGA-VS-A8QF-01A-21R | 67.438 | 0.051 | 0.287 | 1.076 | -0.179 | -0.301 | -0.262 | 0.000 | -0.510 | -0.457 | 0.090 | -0.218 | -0.344 | -0.088 | -0.298 | NA | NA |
| TCGA-JW-A852-01A-11R | 13.870 | -1.262 | -0.87 | -1.809 | -0.538 | -0.529 | -0.302 | 0.000 | -0.494 | -0.761 | -0.351 | -0.305 | -0.457 | 0.240 | -0.304 | NA | NA |
| TCGA-EA-A556-01A-11R | -7.586 | 0.046 | -0.97 | -0.681 | 0.692 | 0.788 | 0.435 | 0.000 | -0.509 | 2.316 | -0.015 | 0.203 | -0.915 | -0.320 | 0.052 | NA | NA |
| TCGA-DG-A2KM-01A-11R | 10.361 | -0.976 | 0.113 | -0.854 | 0.862 | -0.003 | -0.219 | 0.000 | -0.513 | 0.342 | -0.280 | -0.316 | -0.726 | 2.759 | 0.036 | NA | NA |
| TCGA-VS-A9UV-01A-11R | 70.637 | -0.443 | 0.457 | -1.351 | -0.541 | -0.425 | -0.294 | 0.000 | -0.480 | -0.612 | -0.338 | -0.311 | 0.298 | -0.435 | -0.281 | NA | NA |
| TCGA-C5-A2M1-01A-11R | 15.179 | -0.951 | -1.037 | 0.307 | -0.171 | -0.463 | -0.293 | 0.000 | -0.516 | -0.658 | 0.761 | -0.331 | -1.147 | -0.426 | -0.309 | NA | NA |
| TCGA-C5-A2LV-01A-11R | -25.719 | -0.444 | -0.510 | -0.921 | 1.000 | 2.912 | 4.813 | 0.000 | 0.124 | -0.669 | -0.204 | -0.235 | 4.662 | 1.724 | 0.396 | NA | NA |
| TCGA-EK-A2IR-01A-11R | 32.783 | -0.891 | -0.510 | -0.690 | -0.546 | -0.456 | -0.273 | 0.000 | -0.511 | -0.655 | -0.312 | -0.243 | 1.554 | -0.472 | -0.231 | NA | NA |
| TCGA-DS-A1OB-01A-11R | 62.882 | -0.460 | 0.326 | 0.132 | -0.316 | -0.143 | -0.253 | 0.000 | -0.504 | -0.541 | -0.226 | -0.280 | -0.448 | -0.242 | -0.211 | NA | NA |
| TCGA-EA-A5ZF-01A-11R | 26.703 | 1.950 | -1.086 | -0.613 | -0.043 | 0.725 | 0.614 | 0.000 | -0.468 | 1.549 | -0.081 | 0.092 | -0.796 | -0.325 | -0.097 | NA | NA |
| TCGA-XS-A8TJ-01A-11R | 104.564 | 1.853 | 0.689 | -0.816 | -0.336 | -0.367 | -0.271 | 0.000 | -0.506 | -0.277 | -0.351 | -0.321 | -0.061 | -0.277 | -0.298 | NA | NA |
| TCGA-ZJ-AAX4-01A-11R | 98.706 | 0.863 | 1.087 | 1.249 | 0.096 | -0.352 | -0.285 | 0.000 | -0.456 | -0.355 | -0.171 | -0.322 | -0.735 | 0.503 | -0.265 | NA | NA |
| TCGA-C5-A1M6-01A-11R | -10.913 | -0.610 | -1.209 | -0.864 | -0.184 | 0.749 | -0.158 | 0.000 | -0.408 | 0.321 | 0.100 | 1.573 | -1.110 | -0.166 | 0.204 | NA | NA |
| TCGA-FU-A3EO-01A-11R | -1.073 | -1.456 | -1.229 | -1.223 | -0.264 | -0.485 | -0.266 | 0.000 | 1.652 | -0.238 | -0.277 | -0.325 | -1.162 | -0.397 | -0.275 | NA | NA |

| | | | | | | | | | | | | | | | | | |
|---|---|---|---|---|---|---|---|---|---|---|---|---|---|---|---|---|---|
| TCGA-BI-A20A-01A-11R | 25.862 | -0.764 | -0.620 | -0.253 | -0.181 | -0.427 | -0.267 | 0.000 | -0.438 | -0.231 | -0.307 | -0.330 | -0.072 | -0.179 | -0.240 | NA | NA |
| TCGA-JW-A5VG-01A-11R | 60.964 | 0.563 | -0.128 | 2.090 | -0.333 | -0.060 | 0.030 | 0.000 | 1.365 | -0.667 | -0.101 | 0.222 | 0.105 | -0.243 | -0.300 | NA | NA |
| TCGA-C5-A7CG-01A-11R | 26.759 | -0.846 | -0.302 | -1.170 | 0.304 | -0.402 | -0.277 | 0.000 | -0.506 | -0.428 | -0.328 | -0.291 | -0.020 | 0.176 | -0.237 | NA | NA |
| TCGA-VS-A8EC-01A-11R | 61.819 | 0.159 | -0.160 | -0.146 | -0.631 | -0.489 | -0.299 | 0.000 | -0.509 | -0.692 | -0.301 | -0.300 | 0.155 | -0.373 | -0.309 | NA | NA |
| TCGA-FU-A23K-01A-11R | -1.731 | -0.949 | -1.218 | -1.565 | 0.210 | -0.432 | -0.013 | 0.000 | -0.502 | -0.238 | -0.032 | -0.282 | -1.029 | -0.200 | 0.103 | NA | NA |
| TCGA-C5-A2LY-01A-31R | -12.565 | -1.287 | -0.058 | -1.106 | -0.005 | -0.505 | -0.278 | 0.000 | -0.511 | -0.664 | -0.360 | 10.657 | 0.134 | 0.358 | -0.306 | NA | NA |
| TCGA-C5-A1M9-01A-11R | 14.005 | 0.136 | -0.908 | -0.421 | -0.364 | 2.177 | 1.598 | 0.000 | -0.503 | -0.651 | -0.158 | -0.238 | -1.129 | -0.345 | -0.282 | NA | NA |
| TCGA-UC-A7PI-01A-11R | 57.619 | 1.900 | -0.975 | 0.301 | -0.541 | -0.480 | -0.210 | 0.000 | 2.639 | -0.426 | -0.376 | -0.327 | -1.157 | -0.419 | -0.300 | NA | NA |
| TCGA-C5-A1MI-01A-11R | 173.844 | 6.488 | 0.816 | 2.205 | -0.266 | -0.254 | -0.206 | 0.000 | -0.283 | -0.354 | 0.086 | -0.312 | -1.075 | -0.236 | -0.294 | NA | NA |
| TCGA-C5-A1M5-01A-11R | 40.949 | -0.570 | -0.189 | -0.270 | -0.208 | -0.420 | -0.283 | 0.000 | -0.480 | -0.226 | -0.327 | -0.269 | 0.621 | -0.088 | -0.232 | NA | NA |
| TCGA-MA-AA3W-01A-11R | 75.039 | 0.428 | 0.441 | -0.512 | -0.147 | -0.428 | -0.289 | 0.000 | -0.422 | -0.688 | -0.195 | -0.314 | 0.049 | 0.197 | -0.236 | NA | NA |
| TCGA-DS-A0VN-01A-21R | 43.080 | -0.925 | 0.545 | 0.034 | 0.427 | -0.070 | -0.245 | 0.000 | -0.237 | 0.043 | 0.258 | 0.698 | -0.186 | 0.389 | -0.075 | NA | NA |
| TCGA-EK-A2PG-01A-11R | 59.350 | -0.591 | 0.069 | -0.733 | -0.635 | -0.537 | -0.300 | 0.000 | -0.239 | -0.817 | -0.342 | -0.329 | 0.624 | -0.431 | -0.328 | NA | NA |
| TCGA-MU-A8JM-01A-11R | 49.960 | 0.005 | -0.077 | 0.463 | -0.203 | -0.288 | -0.208 | 0.000 | 0.038 | -0.192 | -0.049 | -0.057 | 1.686 | -0.211 | -0.048 | NA | NA |
| TCGA-JW-AAVH-01A-11R | 81.461 | 1.043 | 0.015 | 0.671 | -0.582 | -0.497 | -0.280 | 0.000 | -0.497 | -0.648 | -0.363 | -0.328 | -0.677 | -0.462 | -0.223 | NA | NA |
| TCGA-C5-A902-01A-11R | 52.124 | -1.408 | 0.321 | -1.357 | -0.452 | -0.505 | -0.292 | 0.000 | 0.355 | -0.684 | 2.601 | -0.144 | -1.038 | -0.442 | -0.303 | NA | NA |
| TCGA-VS-A957-01A-11R | 62.551 | 0.011 | -0.068 | 0.188 | -0.553 | -0.486 | -0.140 | 0.000 | -0.513 | -0.788 | -0.278 | -0.327 | -0.581 | -0.174 | -0.213 | NA | NA |
| TCGA-DS-A7WH-01A-22R | -17.304 | -1.014 | -1.020 | 1.258 | 0.851 | 0.998 | 0.491 | 0.000 | -0.435 | 0.978 | 0.963 | 0.218 | -0.656 | -0.457 | -0.073 | NA | NA |
| TCGA-EK-A2H1-01A-11R | -55.745 | -0.034 | -0.87 | -0.904 | 1.569 | 5.111 | 3.243 | 0.000 | 3.378 | -0.203 | 0.443 | 0.511 | 0.329 | 1.853 | 0.733 | NA | NA |
| TCGA-MY-A5BF-11A-11R | -70.464 | -1.591 | -1.24 | -0.731 | 4.173 | -0.099 | 0.162 | 0.000 | -0.092 | 2.289 | 2.068 | -0.314 | 0.566 | -0.436 | -0.099 | NA | NA |

| | | | | | | | | | | | | | | | | | |
|---|---|---|---|---|---|---|---|---|---|---|---|---|---|---|---|---|---|
| TCGA-Q1-A5R1-01A-11R | 29.914 | 0.143 | -1.03 | -0.230 | -0.230 | -0.525 | -0.277 | 0.000 | -0.495 | -0.584 | -0.043 | -0.322 | -1.115 | -0.398 | -0.251 | NA | NA |
| TCGA-VS-A952-01A-11R | 61.382 | 1.020 | -0.58 | 0.861 | -0.546 | -0.483 | -0.267 | 0.000 | -0.508 | -0.605 | -0.259 | -0.329 | -1.134 | -0.393 | -0.315 | NA | NA |
| TCGA-IR-A3LI-01A-11R | 37.944 | 0.323 | -1.01 | -0.514 | -0.537 | -0.524 | -0.298 | 0.000 | -0.514 | -0.805 | -0.327 | -0.335 | -1.191 | -0.475 | -0.323 | NA | NA |
| TCGA-VS-A8EL-01A-11R | 20.561 | -0.319 | -0.90 | -0.458 | -0.227 | -0.245 | -0.264 | 0.000 | -0.279 | -0.434 | -0.085 | -0.258 | -0.395 | 0.100 | -0.226 | NA | NA |
| TCGA-Q1-A5R3-01A-11R | -21.089 | -0.286 | -0.94 | -1.202 | -0.448 | -0.513 | -0.200 | 0.000 | -0.270 | -0.466 | -0.247 | -0.246 | -1.012 | 7.119 | -0.256 | NA | NA |
| TCGA-BI-A0VS-01A-11R | 129.205 | -0.011 | 3.095 | -1.985 | -0.104 | 1.083 | -0.204 | 0.000 | -0.495 | 0.422 | -0.162 | -0.185 | 3.944 | 0.454 | -0.127 | NA | NA |
| TCGA-ZJ-AAXT-01A-11R | 74.222 | -0.637 | 0.789 | 0.013 | -0.483 | -0.390 | -0.225 | 0.000 | -0.401 | -0.671 | -0.229 | 0.003 | 0.861 | 0.140 | -0.140 | NA | NA |
| TCGA-VS-A9V1-01A-11R | 23.917 | -0.143 | -1.17 | -0.386 | -0.588 | -0.500 | -0.301 | 0.000 | -0.021 | -0.498 | -0.285 | -0.001 | -1.168 | -0.457 | -0.302 | NA | NA |
| TCGA-FU-A5XV-01A-11R | 131.228 | 0.614 | 1.955 | 0.540 | -0.545 | -0.427 | -0.274 | 0.000 | -0.480 | -0.383 | -0.283 | 0.356 | 0.644 | -0.338 | -0.213 | NA | NA |
| TCGA-HM-A3JJ-11A-12R | -75.284 | -1.910 | -1.30 | 1.475 | 2.900 | 0.429 | -0.266 | 0.000 | -0.394 | 4.841 | -0.061 | -0.334 | 1.030 | -0.479 | 1.340 | NA | NA |
| TCGA-EA-A1QS-01A-61R | -10.178 | 0.111 | -0.20 | -0.101 | 1.613 | 1.431 | 0.964 | 0.000 | 1.297 | 4.706 | 0.042 | -0.250 | 1.545 | -0.257 | 0.355 | NA | NA |
| TCGA-C5-A1M8-01A-21R | 69.614 | 0.823 | -0.10 | 0.274 | -0.369 | -0.431 | -0.269 | 0.000 | -0.509 | -0.549 | -0.113 | -0.326 | -0.602 | -0.258 | -0.289 | NA | NA |
| TCGA-FU-A770-01A-11R | 16.402 | -1.017 | -1.06 | 1.341 | -0.413 | -0.471 | -0.131 | 0.000 | -0.517 | -0.547 | -0.201 | -0.181 | -1.099 | -0.290 | -0.157 | NA | NA |
| TCGA-EA-A5O9-01A-11R | 21.949 | -1.291 | -0.79 | 0.442 | -0.468 | -0.489 | -0.300 | 0.000 | -0.513 | -0.711 | -0.255 | -0.209 | -0.266 | -0.436 | -0.279 | NA | NA |
| TCGA-EK-A3GK-01A-11R | 51.817 | 0.534 | -0.72 | 1.525 | -0.570 | -0.490 | -0.279 | 0.000 | -0.510 | -0.760 | 0.461 | -0.327 | -1.183 | -0.338 | -0.303 | NA | NA |
| TCGA-EA-A3HU-01A-11R | 62.494 | -0.196 | 0.239 | -0.930 | -0.288 | -0.392 | -0.291 | 0.000 | -0.508 | -0.195 | -0.239 | -0.277 | -0.567 | -0.339 | -0.096 | NA | NA |
| TCGA-C5-A7CL-01A-11R | 66.973 | -0.326 | 0.848 | 0.022 | -0.086 | 0.223 | -0.220 | 0.000 | -0.205 | 0.602 | -0.057 | -0.086 | 0.803 | -0.097 | -0.104 | NA | NA |
| TCGA-C5-A1M7-01A-11R | 101.735 | 1.177 | 0.829 | 1.501 | -0.402 | -0.482 | -0.294 | 0.000 | -0.513 | -0.143 | -0.039 | 0.319 | -0.440 | -0.227 | -0.229 | NA | NA |
| TCGA-C5-A7UH-01A-11R | 200.719 | 0.785 | 4.149 | 0.097 | -0.349 | -0.161 | -0.280 | 0.000 | -0.406 | -0.491 | -0.142 | -0.064 | 0.390 | 0.006 | -0.282 | NA | NA |
| TCGA-C5-A2M2-01A-21R | 37.684 | 0.409 | -1.07 | 0.353 | -0.613 | -0.460 | -0.289 | 0.000 | -0.515 | -0.623 | -0.354 | -0.263 | -1.169 | -0.466 | -0.300 | NA | NA |

| | | | | | | | | | | | | | | | | | |
|---|---|---|---|---|---|---|---|---|---|---|---|---|---|---|---|---|---|
| TCGA-MY-A913-01A-11R | 44.423 | -0.009 | 0.838 | -0.848 | -0.030 | 3.002 | 0.082 | 0.000 | -0.376 | -0.519 | -0.157 | 1.244 | 0.906 | 0.525 | -0.167 | NA | NA |
| TCGA-VS-A8QC-01A-11R | 128.917 | -0.140 | 2.542 | -0.200 | -0.253 | 0.048 | -0.222 | 0.000 | 0.128 | 0.489 | -0.164 | -0.256 | 0.482 | -0.045 | -0.145 | NA | NA |
| TCGA-VS-A9U5-01A-11R | 50.900 | -0.161 | -0.26 | -0.162 | -0.401 | -0.467 | -0.289 | 0.000 | -0.501 | -0.623 | -0.259 | -0.322 | -0.640 | -0.138 | -0.301 | NA | NA |
| TCGA-DS-A1OC-01A-11R | 58.707 | -0.082 | 0.115 | -0.431 | -0.255 | -0.372 | -0.231 | 0.000 | -0.304 | -0.495 | 0.034 | -0.216 | 0.662 | -0.227 | -0.116 | NA | NA |
| TCGA-EA-A3HQ-01A-11R | 68.582 | -0.358 | 0.724 | 2.431 | -0.054 | 0.426 | -0.253 | 0.000 | -0.508 | 0.269 | -0.302 | -0.292 | 0.855 | -0.331 | -0.285 | NA | NA |
| TCGA-JW-A5VH-01A-11R | 22.421 | 0.406 | -0.56 | -0.626 | 1.045 | 0.243 | 1.073 | 0.000 | -0.208 | -0.401 | 0.408 | -0.287 | -0.771 | -0.283 | 2.479 | NA | NA |
| TCGA-FU-A2QG-01A-11R | 111.838 | 0.162 | 1.434 | 0.325 | -0.618 | -0.484 | -0.295 | 0.000 | -0.265 | -0.620 | -0.308 | 0.244 | -0.222 | -0.328 | -0.211 | NA | NA |
| TCGA-FU-A3YQ-01A-11R | 43.197 | -0.261 | -0.07 | 1.132 | -0.131 | -0.220 | -0.275 | 0.000 | -0.256 | 0.419 | -0.190 | -0.273 | 0.168 | 0.330 | -0.092 | NA | NA |
| TCGA-Q1-A6DT-01A-11R | 62.524 | 0.229 | 0.062 | 0.478 | -0.381 | -0.450 | -0.300 | 0.000 | 0.632 | -0.638 | 1.302 | 0.176 | 0.418 | -0.203 | -0.229 | NA | NA |
| TCGA-VS-A8EI-01A-11R | 51.486 | -0.810 | 0.144 | 1.571 | -0.347 | -0.443 | -0.277 | 0.000 | 1.192 | 0.192 | -0.227 | -0.277 | 0.513 | -0.247 | -0.183 | NA | NA |
| TCGA-VS-AA62-01A-11R | 89.324 | 0.527 | 1.145 | 0.063 | -0.112 | 0.595 | -0.188 | 0.000 | -0.264 | -0.223 | -0.066 | -0.044 | -0.568 | 0.232 | -0.247 | NA | NA |
| TCGA-JW-A69B-01A-11R | 43.106 | 2.049 | -1.12 | -1.456 | -0.053 | -0.273 | -0.067 | 0.000 | -0.512 | -0.030 | -0.207 | -0.310 | -0.991 | -0.277 | -0.269 | NA | NA |

#### Supplementary File 4: Primers used in this study.

| Oligo Name | Sequence 5' →3' | |
|---|---|---|
| **BMP7 NM_001719.3** | For | ACCAGAGGCAGGCCTGTAAGA |
| | Rev | CTCACAGTTAGTAGGCGGCGTAG |
| **CAV1 NM_001753.5** | For | AAGGGACACACAGTTTTGACG |
| | Rev | TTGGCACCAGGAAAATTAAAA |
| **ID2 NM_002166.5** | For | TGGACTCGCATCCCACTATT |
| | Rev | ATTCAGAAGCCTGCAAGGAC |
| **CDH1 (NM_004360.3)** | For | CCCGCCTTATGATTCTCTGCTCGTG |
| | Rev | TCCGTACATGTCAGCCAGCTTCTTG |
| **VIM (NM_003380.5)** | For | GGCTCAGATTCAGGAACAGC |
| | Rev | AGCCTCAGAGAGGTCAGCAA |
| **β-Actin (NM_001101.5)** | For | TCATGAAGATCCTCACCGAG |
| | Rev | TTGCCAATGGTGATGACCTG |
